# Connecting the dots: deep learning-based automated model building methods in cryo-EM

**DOI:** 10.64898/2025.12.28.696775

**Authors:** Harsh Bansia, Amedee des Georges

## Abstract

The resolution revolution in single particle cryo-electron microscopy (cryo-EM) has dramatically expanded our structural knowledge of large biomolecular complexes. While high-resolution cryo-EM density maps enable atomic model building, lower-resolution maps can still reveal secondary structures, folds, and domains. When combined with integrative modeling approaches, such data can provide meaningful insights into biomolecular structure and function. Constructing accurate models, however, remains challenging: at low resolutions it is difficult to interpret density maps features reliably, and at high resolutions traditional model-building workflows can become a time-consuming bottleneck. Deep learning, which is transforming problem-solving across scientific domains, offers powerful new tools to automate and accelerate this process. In this review, we discuss deep learning-based methods developed to automate model building in cryo-EM density maps, assessing their impact on streamlining structure determination. Recognizing that biomacromolecular structures exhibit hierarchical organization, we classify these methods according to their ability to model primary, secondary, tertiary, and quaternary structures of biomolecules. Deep learning tools for building atomic models in cryo-EM density maps are further grouped as *de novo*, where the model is predicted directly from features learned from the cryo-EM density, or hybrid, where it is derived by integrating structural templates with these features. We outline current limitations, including the challenge of obtaining sufficiently large and diverse datasets for training networks to model different types of biomolecules in the cryo-EM density maps, and the open challenge of constructing training sets that capture the conformational heterogeneity often observed in the cryo-EM maps. We conclude by highlighting emerging directions for this rapidly advancing field, which promise to make automated, data-driven model building an integral part of structural biology.

## 1 Introduction

Prominent architect Louis Sullivan coined the phrase “*form follows function*” asserting that an object’s shape should primarily relate to its intended purpose. He applied this design philosophy to pioneer the modern skyscraper. The design principle quickly found applications in other domains besides architecture, including automobile design, product design, software engineering, and even describes the hierarchy of biological organization at cellular, tissue, organ, individual, and ecosystem levels. At the molecular level “*form follows function*” implies how a particular arrangement of atoms, residues, and higher order motifs in a biomacromolecule allows it to perform a specific function e.g. enzyme active-site catalysis, membrane protein assisted transport of solutes, flow of genetic information, underscoring the need to accurately determine macromolecular structure to elucidate its function [1]. Unlike skyscrapers, which are static, biomolecules can dynamically adapt their *forms* by spatially rearranging their constituent elements to perform a diverse array of *functions* in rapidly changing cellular environments, the basis for allosteric mechanisms [2].

Structure determination of protein–protein and protein–nucleic acid complexes is crucial for understanding cellular processes and developing therapeutics for challenging disorders [3–5]. Single-particle cryo-electron microscopy (cryo-EM) has become the method of choice for determining high-resolution structures of biomacromolecules, ranging from small proteins to large complex assemblies in their native context [6–9]. A three-dimensional (**3D) density map** in cryo-EM is represented as a 3D grid, with each voxel assigned to a density value. A **voxel** is the 3D equivalent of a two-dimensional (2D) pixel and represents a value in a 3D grid, analogous to how a pixel represents a value in a 2D grid. A 3D cryo-EM density map is reconstructed from thousands of 2D projection images of randomly oriented, flash-frozen biomolecules (single particles) embedded in vitreous ice, providing an approximation of the electron scattering potential of the biomolecule [10,11]. These density maps provide the structural framework for building and refining models of biomacromolecules, revealing molecular mechanisms and conformational states that may be difficult to capture with other structural techniques [12]. The structure of biomacromolecules is often described in terms of hierarchical levels of complexity and organization ranging from local, regular secondary structures such as β-sheets and α-helices to global, complex 3D arrangement of atoms capturing interactions between regions that may be far apart in the primary structure. The resolution of the density map determines the level of detail, from overall molecular shapes and secondary structures at lower resolutions to 3D arrangement of individual residues and atomic features at near-atomic resolution. Model building involves the construction of plausible models of biomacromolecule within the interpretable density of the map while maintaining correct chemical, stereochemical, and geometrical properties of the biomacromolecule. The level of detail achievable in the model depends on the map resolution. Therefore, successful model building ultimately depends on the quality and resolution of the density maps. At modest to lower resolutions, model building becomes increasingly challenging, often leading to errors in models deposited in the Protein Data Bank (PDB) [13–16]. Moreover, as the biomolecule studied by cryo-EM grow in size and complexity, a now common situation, maps frequently contain densities corresponding to regions without homologous templates in the PDB. This lack of structural precedent makes *de novo* atomic model reconstruction particularly difficult and time-consuming [17]. Figure 1 shows that as of the end of the year 2024, the Electron Microscopy Data Bank (EMDB) [18,19] contained 9,329 entries, of which 6,877 (73.7 %) fell within the model building range (0 – 4.0 Å). However, only 5,791 entries had corresponding atomic models deposited in the PDB, representing just 62.1 % of all EMDB submissions and 84.2 % of those in the 0 – 4.0 Å resolution range [19] (Figure 1). Similar trend can be observed for the year 2025 where deposited models (5,571) in PDB corresponds to 62.6 % (8,893) of all EMDB submissions and 83.2 % (6,696) of those in the 0 – 4.0 Å resolution range [19] (Figure 1). While the reasons for this discrepancy may be multifaceted, it underscores the persistent challenge in generating complete atomic models even for high-resolution data. As such, accurate model building of target macromolecular assemblies from density maps is one of the most demanding tasks in any structure determination pipeline, requiring labor-intensive manual inputs reliant on complex judgements, clearly necessitating development of advanced model building tools in structural biology.

**Figure 1.**
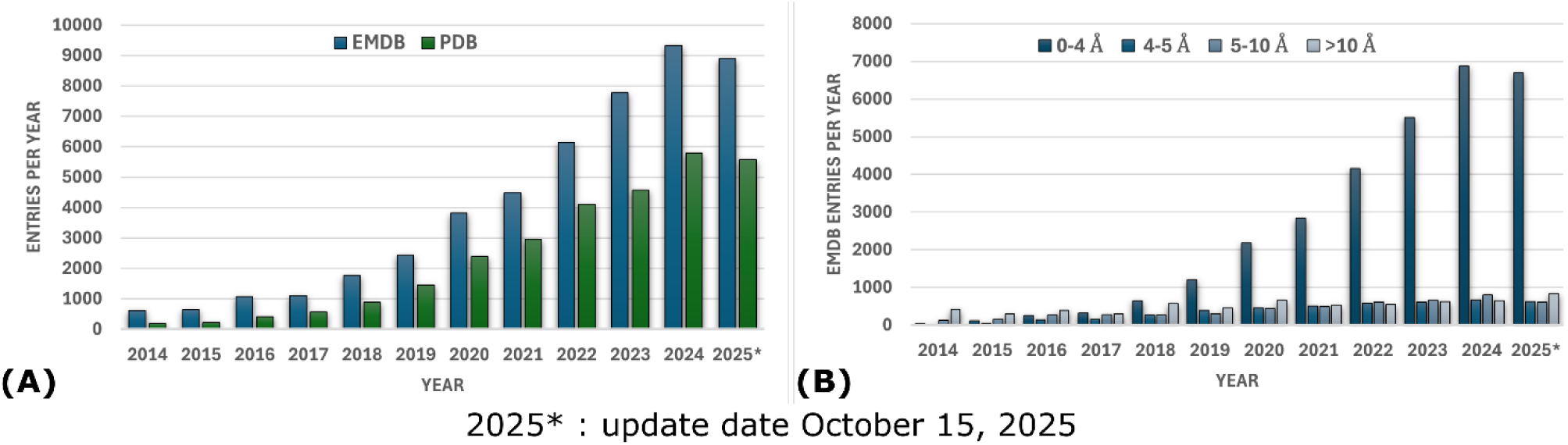
(A) Yearly distribution of total EMDB entries and associated PDB entries for the last decade. (B) Distribution of EMDB entries by resolution per year. Data sourced from EMDataResource [19].

Although methods for reconstructing models [20–24], which iteratively optimize and refine structures by minimizing an energy function, have accelerated cryo-EM structure determination, they often require expert, informed manual input to derive accurate models. Traditional machine learning algorithms have been used to develop methods aimed at automating structure determination from cryo-EM density maps. These approaches are generally rule-based or rely on statistical learning techniques to guide macromolecular modeling from cryo-EM maps [25–27]. SSELearner [25] automatically identifies helices and β-strands by using *Support Vector Machine* (SVM) to classify voxels in cryo-EM density maps. RENNSH [26] only identifies α-helices using nested *K Nearest Neighbors* (KNN) classifiers to distinguish helix from non-helix voxels. Pathwalking [27] combines *k-means clustering*, an unsupervised machine learning algorithm, with a Traveling Salesman Problem (TSP) solver, a combinatorial optimization algorithm, to automatically trace the protein backbone directly from cryo-EM maps at 3.0–6.0 Å resolution. However, the above methods are mostly limited to either identifying secondary structure elements or tracing the minimal protein backbone, thereby highlighting the need for effective atomic model-building methods from cryo-EM density maps to provide a complete modeling solution.

The resolution revolution in cryo-EM, driven by hardware and software advances, has led to the generation of large number of high-resolution density maps [8,9,28]. These advances, combined with parallel developments in computational hardware, software, and machine learning methodologies, have facilitated the development of deep learning-based tools to automate critical steps in the cryo-EM structure determination workflow. Deep learning is a branch of machine learning that employs artificial neural networks, modeled after biological neural networks, for prediction and classification tasks, with the term ‘deep’ referring to the use of multiple hidden layers in the network [29]. Deep learning has revolutionized problem solving across science [30], exemplified by AlphaFold2’s Nobel Prize winning success in protein structure prediction [31]. Within the cryo-EM workflow itself, deep learning has already significantly improved steps like particle picking, 3D density map reconstruction and structural heterogeneity analysis [32–34]. Building on this momentum, a new wave of deep learning methods now targets the critical challenge of automating reconstruction of models from cryo-EM density maps [35–38] – the focus of this review.

We begin with a brief primer on key aspects of biomolecular structures, followed by an overview of the fundamentals of deep neural networks, with the aim of providing a general and accessible introduction for readers from diverse fields to the terminology and concepts used throughout the review. We then examine approximately 50 deep learning-based tools for modeling proteins, nucleic acids, and protein-nucleic acid complexes in cryo-EM density maps, analyzing their underlying architectures to identify common conceptual strategies, outlining their main stages from data preprocessing and training to feature learning from cryo-EM density maps, model building, and subsequent refinement. Recognizing the hierarchical organization of biomolecular structures, we classify these tools according to their ability to model the primary, secondary, tertiary, and quaternary structures of biomolecules. Tools for building atomic models in cryo-EM density maps are further grouped as *de novo*, where the atomic model is predicted directly from features learned from the cryo-EM density, or hybrid, where it is derived by integrating structural templates with these features. Subsequent sections address the assessment and validation of structural models built by these tools in cryo-EM density maps, the availability and applications of these tools, and their current limitations and potential future improvements. We complement the sections with figures and tables summarizing, for each tool, its training datasets, neural network architecture, prediction tasks, the types of biomolecules it builds, and its availability as servers or publicly accessible code. Our comprehensive review of deep learning-based methods for automated model building in cryo-EM density maps provides a timely, useful, and one-stop resource for structural biologists, researchers interested in applying these methods in cryo-EM workflows, and method developers aiming to design next-generation model-building tools.

## 2 A primer on biomacromolecular structure

Biomacromolecules are large, complex molecules typically formed by the polymerization of smaller repeating units called monomers. In biological systems, biomacromolecules are essential for life and most commonly include proteins, nucleic acids and carbohydrates. Nucleic acids - deoxyribonucleic acid (DNA) and ribonucleic acid (RNA) - store and transmit the genetic information essential for life. Proteins are the workhorses of biological systems, playing critical roles in virtually every cellular process including catalyzing biochemical reactions as enzymes, providing structural support, transporting molecules, and regulating cellular processes. The structure of biomacromolecules is often described in terms of hierarchical levels of complexity and organization. For proteins and nucleic acids, these are commonly referred to as **primary, secondary, tertiary,** and sometimes **quaternary structures**.

Proteins are linear biopolymers of repeating units called **amino acids.** There are **20 different types of amino acids** commonly found in proteins. An amino acid has a central **α-carbon** (**Cα**) atom to which four different groups are covalently attached: an amino group (-NH₂), a carboxylic acid group (-COOH), a hydrogen atom (-H), and a **side chain** (-R group) unique to each amino acid. The side chain imparts distinct physicochemical properties to the amino acids (such as polar, non-polar, aromatic, aliphatic, acidic, basic) and thereby contributes to structural and functional diversity in proteins. Successive amino acids in a protein are covalently linked by **peptide bonds**, formed between the carboxyl group of one amino acid and the amino group of the next, creating the **main chain or backbone (repeating -N-Cα-C**-), to which distinctive side chains are attached (Figure 2A). **Polypeptides** are polymers of a series of amino acids linked by peptide bonds, and each amino acid within a polypeptide chain is referred to as a **residue** (Figure 2A). Proteins consist of one or more polypeptide chains. A polypeptide chain has directionality, with an amino-terminal (N-terminal), where the terminal residue has a free amino group, and a carboxy-terminal (C-terminal), where the terminal residue has a free carboxyl group. By convention, the sequence of amino acids in a polypeptide is written starting with the N-terminal residue. In successive amino acid residues within a polypeptide, the peptide bond is planar, restricting free rotation around the C–N bond. However, free rotation is permitted around the N–Cα and Cα–C bonds, which are quantified by the backbone **dihedral (torsion) angles φ (phi) and ψ (psi)**, respectively (Figure 2A). These angles define the backbone conformation of a polypeptide, allowing proteins to fold into a variety of distinct structures. The polypeptide backbone is rich in hydrogen-bonding potential because each peptide bond contains both a hydrogen-bond donor (the NH group) and a hydrogen-bond acceptor (the CO group), allowing the backbone to stabilize structural elements within a protein. The **primary structure** of a protein refers to its linear **sequence** of amino acid **residues**, defined by the peptide bonds linking them along the polypeptide chain(s) (Figure 2A). For any given segment of a polypeptide chain, **secondary structure** refers to the local spatial arrangement of its main-chain (backbone) atoms. The secondary structure of a polypeptide segment is completely determined when the backbone dihedral angles, φ (phi) and ψ (psi), are known for each residue. When these angles adopt a repeating, consistent value throughout the segment, a recurring, periodic structural pattern arises, leading to regular secondary structures such as **α-helices** and **β-strands** (Figure 2B). In an α-helix, the polypeptide backbone adopts a right-handed helical conformation, with amino acid side chains projecting outward from the helical axis. The α-helix is stabilized by intra-helix hydrogen bonds between the backbone carbonyl oxygen of residue *i* and the backbone amide hydrogen of residue *i+4* (Figure 2B). Compared to an α-helix, a β-strand represents an extended conformation of the polypeptide backbone. β-Sheets are composed of two or more β-strands arranged side-by-side in an antiparallel or parallel manner, having opposite or the same amino-to-carboxyl orientations, respectively (Figure 2B). In β-strands, the side chains alternate above and below the plane of the backbone, giving β-sheets their characteristic pleated appearance. Unlike the intra-helix backbone hydrogen bonding in α-helices, β-sheets are stabilized by inter-strand hydrogen bonds between the backbones of adjacent β-strands (Figure 2B). Random coils are examples of secondary structures in which no regular pattern exists. Turns or loops are secondary structure elements that connect successive runs of α-helices or β-strands, often facilitating a reversal of the polypeptide chain direction. An example is a β-turn, which connects the ends of two adjacent β-strands in an antiparallel β-sheet. Unlike secondary structure, which defines the local spatial arrangement of adjacent amino acid residues, **tertiary structure** describes the overall three-dimensional folding of the entire polypeptide chain, including interactions between residues that may be far apart in the primary structure and between different secondary structure elements (Figure 2E). Thus, tertiary structure captures long-range interactions. Secondary structure elements such as α-helices and β-sheets can combine in specific ways through connecting segments to form supersecondary structures, or motifs, such as the helix–turn–helix. Tertiary structure is characterized by distinct overall folding patterns, or folds, that describe the three-dimensional arrangement of these motifs within a single polypeptide chain. A protein **domain** is a region of a polypeptide chain that can fold independently into a stable structure and can often move as a single unit relative to the rest of the chain (Figure 2E). In large proteins, different domains may have distinct structural and functional properties. Many proteins are composed of more than one polypeptide chain, identical or different, associated non-covalently or through covalent disulfide bonds. Each polypeptide chain is called a subunit, and such proteins are referred to as multisubunit or multimeric proteins (Figure 2E). The spatial arrangement and interactions of these subunits define **quaternary structure**, which can range from simple dimers composed of identical subunits to large complexes containing many different subunits (Figure 2E).

**Figure 2.**
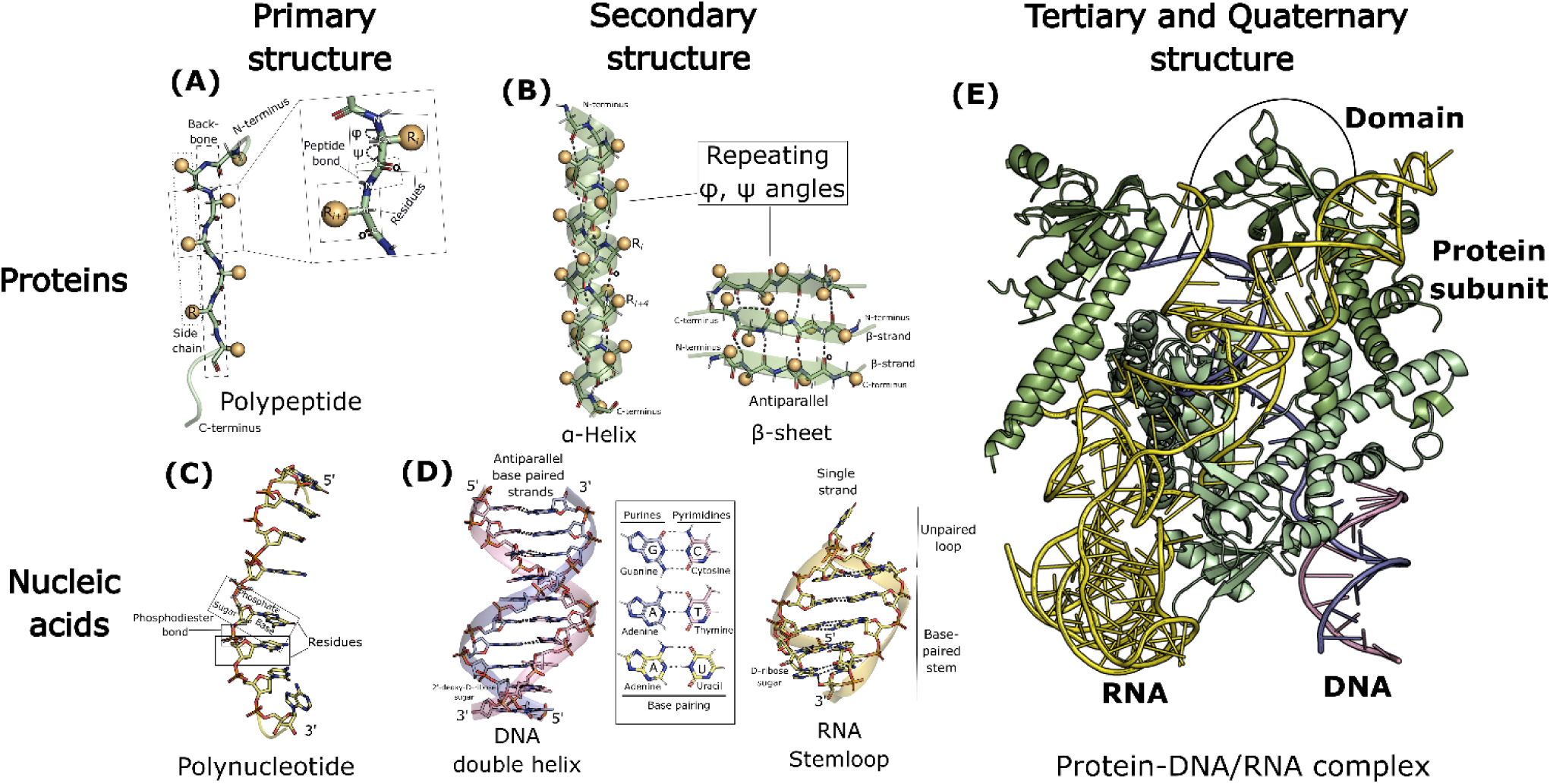
Schematic illustrating the hierarchical organization of biomolecular structures (proteins and nucleic acids), as described in **Section 2**. **(A, C)** Primary structure**. (B, D)** Secondary structure**. (E)** Tertiary and quaternary structure. The illustration was generated in PyMOL [39] using the cryo-EM structure of AsCas12f-sgRNA-target DNA ternary complex (PDB ID: 8J12).

Nucleic acids - deoxyribonucleic acid (DNA) and ribonucleic acid (RNA) - are linear biopolymers of nucleotides. A nucleotide consists of three components – a pentose **sugar**, a nitrogenous (nitrogen-containing) **base**, and a **phosphate** group (Figure 2C). In RNA the sugar is D-ribose whereas in DNA the sugar is 2’-deoxy-D-ribose which lacks the 2’-hydroxyl group (Figure 2D). In a nucleotide, the phosphate group is attached to the 5’ carbon of sugar through a phosphoester bond, whereas the nitrogenous base is covalently attached to the 1’ carbon of sugar via a glycosidic bond (Figure 2C). A nucleotide without a phosphate group is called a nucleoside. In nucleic acids, there are five major nitrogenous bases, which are derivatives of either purines or pyrimidines (Figure 2D). Adenine (A) and Guanine (G) are the major purine bases in both RNA and DNA (Figure 2D). Among the three pyrimidine bases, Uracil (U) occurs in RNA, Thymine (T) is found in DNA while Cytosine (C) is present in both DNA and RNA (Figure 2D). The successive nucleotides in nucleic acids are covalently linked through **phosphodiester linkages** where phosphate group bridges the 3’ and 5’ positions of successive sugar moieties in the adjacent nucleotides, forming the sugar-phosphate backbone to which nitrogenous bases are attached as side groups (Figure 2C). Nucleotides as part of nucleic acids are referred to as **nucleotide residues**. The **primary structure** of a nucleic acid refers to its linear sequence of nucleotide residues, distinguished by their nitrogenous bases, and defined by the phosphodiester bonds linking them along the polynucleotide chain. Polynucleotide chains in nucleic acids have directionality, with the 5′ end having a free phosphate group attached to the 5′ carbon of the terminal sugar, while the 3′ end having a free hydroxyl group on the 3′ carbon of the terminal sugar (Figure 2C, D). The **secondary structure** of a nucleic acid refers to the regular, stable structures like double helices, hairpins, and loops formed by hydrogen bonding patterns between nitrogenous bases within a single polynucleotide chain or between two such chains. The most common form of DNA, known as B-DNA, consists of two helical polynucleotide chains wound around a common axis to form a right-handed double helix, with the nitrogenous bases positioned on the inside and the sugar–phosphate backbone on the outside of the double helix (Figure 2D). The two chains of the double helix have opposite directionality making them antiparallel and are held together by hydrogen bonding between complementary base pairs: A pairs with T and G pairs with C (Figure 2D). This specific base pairing ensures that the two strands are complementary: whenever G occurs in one chain, C is found in the other and likewise for A and T. RNA molecules are mostly single-stranded and can fold into local double-helical regions or stem loop secondary structures, also referred to as hairpin loops (Figure 2D). Similar to DNA, G pairs with C, and A pairs with U in RNA. Self-complementary sequences in RNA form hairpin loops, consisting of a complementary base-paired stem and an unpaired loop at the end (Figure 2D). The **tertiary structure** of a nucleic acid refers to the three-dimensional folding and spatial arrangement of its secondary structure elements, such as helices and hairpin loops, resulting in complex overall shapes (Figure 2E). Common examples include supercoiled cellular DNA and tRNA (transfer RNA).

## 3 A primer on deep neural networks (DNNs)

This primer aims to provide a concise introduction to deep learning concepts and neural network architectures employed in automated model-building methods for cryo-EM. For a more comprehensive understanding of machine learning concepts, readers are referred to the reference [29].

### Artificial neurons and artificial neural networks

Artificial neural networks (ANNs), inspired by biological neural networks, are universal function approximators and can learn to model any mathematical function to a desired degree of accuracy [29]. The fundamental information processing units of ANNs are interconnected artificial neurons, also inspired by biological neurons. A single artificial neuron is a mathematical function that processes and transforms input data (Figure 3A). Each neuron receives one or more inputs (*xi*), multiplies each input by a corresponding learnable weight term (*wi*), sums these weighted inputs, adds a learnable bias term (*b*), and passes the result through a non-linear activation function (*f*) to produce its output (*y*) (Figure 3A) [29]. In other words, an artificial neuron computes a non-linear function of a weighted sum of its inputs.

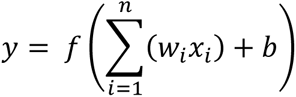

**Figure 3.**
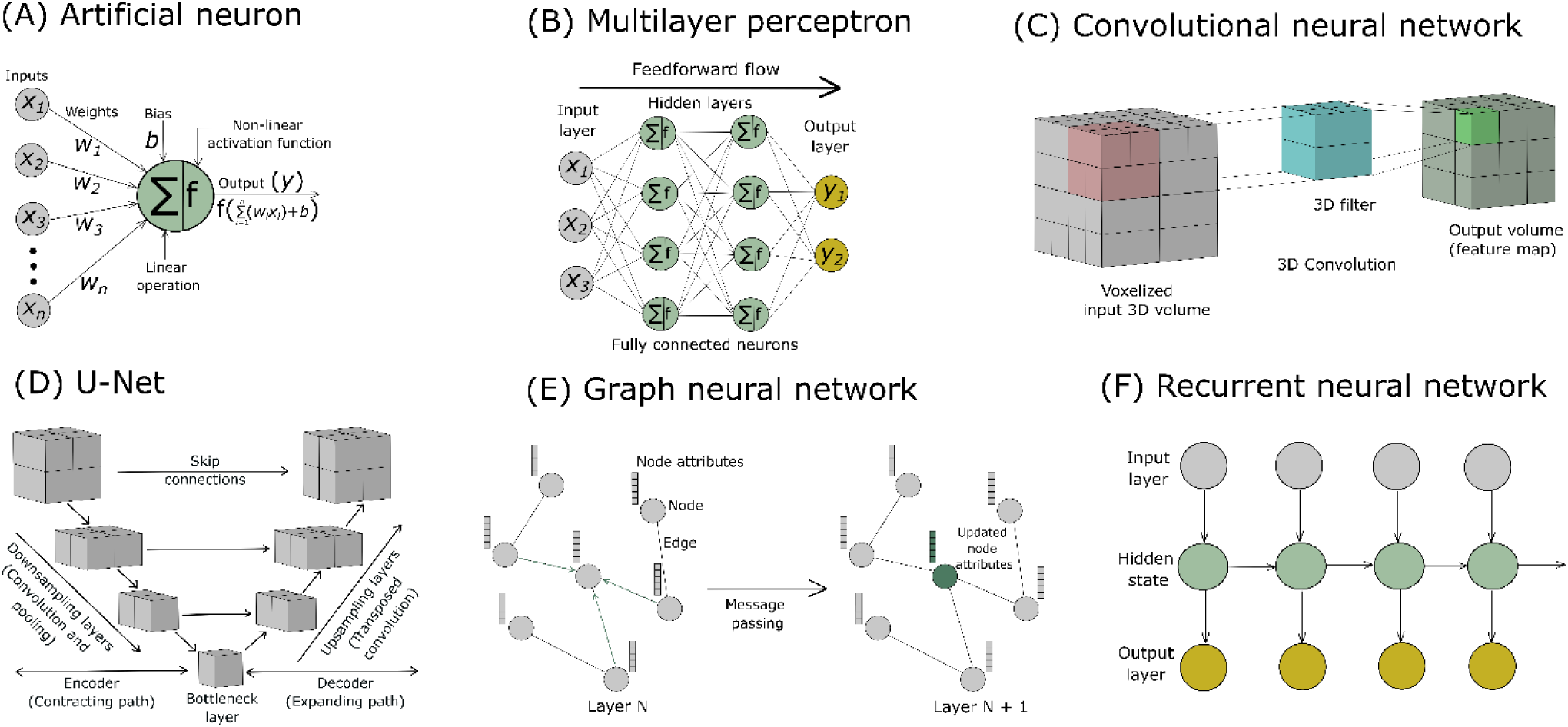
**(A)** Illustration of an artificial neuron showing its inputs, weights, bias, activation function, and output. Schematic overview of representative neural network architectures used by deep learning tools for model building in cryo-EM: **(B)** Multilayer perceptron, **(C)** Convolutional neural network, **(D)** U-Net, **(E)** Graph neural network, and **(F)** Recurrent neural network. These architectures are described in **Section 3**.

Weights control the strength or importance of the connections between neurons, and the bias allows the activation function to shift its output, enabling the model to learn diverse patterns. The activation function introduces non-linearity, transforming the linear operations (weighted sums and biases) into non-linear ones. This allows the network to model complex relationships and effectively act as a universal function approximator. Non-linear patterns are inherent in most real-world data, and activation functions enable neural networks to capture this non-linear structure. In an ANN, artificial neurons are arranged in layers, with neurons in one layer connected to neurons in adjacent layers, while neurons within a layer does not communicate with each other. Layers correspond to different stages of computation in a neural network with the output of one layer serving as the input to the next.

### Neural network architecture

A basic network typically consists of three types of layers: an input layer, one or more hidden layers, and an output layer (Figure 3B). **Deep learning** (DL) is a branch of machine learning that employs ANNs for prediction and classification tasks, with the term ‘**deep’** referring to the use of **multiple hidden layers** in the network (Figure 3B). Each neuron in the input layer typically corresponds to one input feature value calculated from the input data. Hidden layers, located between the input and output layers, process and transform the data: with multiple hidden layers, the network can learn hierarchical patterns and extract higher-level features. The output layer generates the network prediction, with the number of neurons determined by the specific task such as the number of classes in a classification problem. A mathematical function that quantifies the disagreement between the predicted output from a neural network and the ground truth values is known as **loss function** [29]. Essentially, a loss function quantifies the error in predictions from a network. Weights and biases are the adjustable, learnable parameters of a neural network, and they are optimized during the network training process to minimize the chosen loss function. **Backpropagation** [30] is a fundamental algorithm used to train neural networks. It employs the chain rule of calculus to compute the gradients of the loss function with respect to all weights and biases, indicating the direction and magnitude by which they should be adjusted to reduce the error. Subsequently, an **optimization** algorithm utilizes these gradients to update the weights and biases to minimize the loss function and improve network performance. TensorFlow [40] and PyTorch [41] are mainstream software platforms for training neural networks. Backpropagation–optimization is an iterative process, a feedback loop involving the calculation of loss function gradients and subsequent parameter adjustments and is fundamental to the neural network learning process. However, the **vanishing gradient problem** may arise during the training of deep neural networks, where gradients become extremely small as they propagate through layers, making learning slow or ineffective.

Each neuron in a fully connected (**FC**) layer, also known as dense layer, is connected to every neuron in the previous layer, and its output is passed to every neuron in the next layer (Figure 3B). A **feedforward neural network** (**FNN**) is a type of ANN where information flows in only one direction, from the input layer, through one or more hidden layers, to the output layer, without feedback loops or cycles (Figure 3B). **Multilayer perceptrons** (MLPs) are a class of FNNs identified by at least three layers: an input layer, one or more hidden layers, and an output layer, with neurons typically connected in a dense or fully connected (FC) manner (Figure 3B).

The need to handle the unique characteristics of different data modalities has driven the development of specialized neural network architectures [42] (Table 1). For example, the need for efficient local feature extraction in image processing led to the development of convolutional neural networks (CNNs) [43]. Complex neural network architectures often incorporate fully connected layers and MLPs as sub-components. Neural network architectures commonly used in automated model-building methods for cryo-EM are described below.

**Table 1:** Neural network architectures to handle specific data types.

| Data type | Structure | Examples | Architecture |
| --- | --- | --- | --- |
| **Grids** | Regular arrays (1D, 2D, 3D) with fixed spatial relationships | 2D images, 3D cryo-EM density maps | CNNs, U-Nets, Vision Transformers |
| **Graphs** | Irregular connectivity, nodes and edges define relationships | Biomolecular structure, Interaction networks | GNNs, Graph Transformers |
| **Sequential** | Ordered list of elements | Protein/DNA sequences | RNNs, LSTMs, Transformers |

### Grid-structured data and Convolutional Neural Networks (CNNs)

Grid-structured data can be naturally represented as a regular, multi-dimensional array or grid, where each grid element has a fixed spatial or temporal relationship to its neighbors, thereby enabling spatial or temporal locality in the structure. Common examples include images, represented as 2D grids of pixels containing intensity values, and volumetric data such as 3D cryo-EM density maps, represented as 3D grids of voxels containing density values. **Convolutional neural networks (CNNs)** are specifically designed to process grid-structured data by exploiting inherent spatial or temporal locality in the data to detect complex patterns and extract hierarchical features [43,44]. A CNN comprises one or more convolutional layers, where the values in the next layer are computed by applying learnable convolutional filters across the input grid, producing another grid-like layer as output (Figure 3C). Each such output is known as a feature map, which contains specific features extracted from the input data (Figure 3C). The convolution operation enforces local connectivity, such that each neuron in the convolutional layer connects only to a small, localized region of the input such as a patch of pixels in an image or a patch of voxels in a volumetric input (Figure 3C). The extent of this local connectivity is referred to as the receptive field of a neuron in the convolutional layer. **Dilated convolutions** are convolution filters with spacing between their elements, which increases the receptive field. To extract a particular feature, the same filter is applied across the entire input grid. This parameter sharing reduces the number of trainable network parameters and allows CNNs to detect the same feature regardless of its location in the input. In cryo-EM, 3D CNNs are the go-to architecture for processing volumetric 3D density maps, as they can automatically and adaptively learn hierarchical patterns and features. 3D CNNs use 3D convolutional layers, where a 3D filter is applied across the volumetric input along all three spatial dimensions (x, y, z) producing a 3D output volume as a feature map that captures patterns across all dimensions (Figure 3C). Pooling layers are typically inserted between successive convolutional layers to reduce computational load and improve network robustness. They downsample the feature maps generated by convolutional layers by reducing their spatial dimensions. This dimensionality reduction makes detected features more robust to variations in the input (e.g., translations) and decreases computational complexity. Max pooling is commonly used, where a filter slides over the feature map and selects the maximum value from each local region, reducing its size and emphasizing important features. A CNN classifier, located at the end of the CNN architecture after several convolutional and pooling layers, uses fully connected (FC) layers to classify the high-level features extracted from the input grid into specific categories. In an FC layer, each neuron is connected to all neurons of the previous layer, like a conventional multilayer perceptron (MLP) (Figure 3B).

While traditional CNN architecture is well-suited for image classification, many tasks require pixel or voxel-wise classification, such as semantic segmentation, where a label is assigned to every pixel or voxel, and multi-scale feature extraction. The **U-Net** architecture, with its characteristic ‘U’ shape, was specifically designed for this purpose [45]. The architecture of U-Nets consists of an encoder–decoder structure with skip connections (Figure 3D). The **encoder (contracting path)**, like a standard CNN, uses convolution and pooling (**downsampling**) layers to reduce spatial dimension, processing the grid-structured input to extract high-level feature maps and capture semantic information (Figure 3D). The encoder leads to the **bottleneck layer** at the bottom of the ‘U’-shaped architecture, which contains an abstract, compact feature representation called the **latent representation** (Figure 3D). The **decoder (expanding path)** then takes these features and uses deconvolution or transposed convolution (**upsampling**) layers to restore spatial dimension and reconstruct the output (Figure 3D). A key feature of the U-Net is its use of **skip connections**, where the input to each layer (deconvolutional) in the decoder combines the output from the previous layer with the corresponding output from an encoder layer (convolutional), allowing information to bypass intermediate layers and preserve spatial details (Figure 3D). Like 3D CNNs, 3D U-Nets are designed for segmentation of 3D grid-structured data [46]. 3D U-Nets excel at precise voxel-wise segmentation in cryo-EM density maps, accurately identifying backbone atoms, secondary structures, amino acid types, and nucleotides.

The vanishing gradient problem arises when training deep CNNs or U-Nets. **Residual neural networks (ResNets)** use residual connections to address this issue, facilitating the training of much deeper networks [47]. Residual connections are a specific type of skip connection in which the input of a layer is added to its output. Early in training, if some layers are unnecessary, ResNet can learn to ignore them via skip connections, effectively allowing the network to “skip” over these layers and mitigate the vanishing gradient issue. ResNets can serve as a backbone architecture for feature extraction from cryo-EM density maps.

**Diffusion models** are generative machine learning models that learn the underlying probability distribution of a dataset to create new data resembling the original [48]. They operate by gradually adding noise to data (diffusion) and then learning to reverse this process to generate new data (denoising). The neural network architecture underlying the denoising step in diffusion models is often based on the U-Net architecture (Figure 3D). In cryo-EM, diffusion models have shown strong performance in denoising density maps, thereby enhancing model building.

### Graph-structured data and Graph Neural Networks (GNNs)

A graph is a set of nodes (vertices) and edges (links), where each edge connects a pair of nodes (Figure 3E). An attributed graph is a graph in which each node and edge is associated with one or more features, called attributes (Figure 3E). **Graph neural networks (GNNs)** are neural network architectures specifically designed to operate on graph-structured data [42,49]. Attributed graphs are commonly used in GNNs and other applications where each component of the graph carries additional information beyond just connectivity (Figure 3E). In general, graphs do not have a canonical or fixed node ordering, and therefore graph operations are typically defined to be independent of node ordering. Accordingly, GNNs are designed to preserve the permutation symmetry of the input graph. A GNN accepts an attributed graph as input, performs learnable transformations on the node attributes while preserving permutation symmetry, and outputs a graph with updated node attributes but the same connectivity as the input graph (Figure 3E). GNNs progressively transform node representations through message passing operations, where each node iteratively updates its features by aggregating and transforming the features of its neighboring nodes using learnable functions (Figure 3E). **Graph Convolutional Networks (GCNs)** are neural networks designed for graph-structured data, performing convolution-like operations to aggregate and combine information from each node’s neighbors. Biomacromolecules such as proteins and nucleic acids are naturally represented as graph-structured data, where atoms or residues serve as nodes, and the bonds or interactions connecting them serve as edges [50]. In an attributed graph representing a biomacromolecule, nodes (atoms or residues) can carry features such as element or residue type, charge, hydrophobicity, secondary structure, or 3D coordinates [50]. Edges (bonds or interactions) may include attributes like bond type, bond order, interaction type (e.g., hydrogen bond), distance, or interaction energy [50]. These attributes encode chemical, structural, and functional information, enabling richer representations for GNN tasks such as property prediction, structural analysis, or molecular modeling [50].

### Sequential data and Recurrent Neural Networks (RNNs)

While grid and graph-structured input data are spatial in nature, many types of data operated on by machine learning models are sequential, such as text, speech, and time-series information. **Recurrent neural networks (RNNs)** are designed to process sequential data, where the order and context of information are important, and can be used to classify input sequences or predict sequence-dependent properties (Figure 3F) [51]. Their primary applications are in natural language processing (NLP), including speech recognition, language translation, and text generation [52]. Unlike feedforward neural networks, which contain no loops or cycles, neurons in RNNs have recurrent connections that form directed cycles. Recurrent connections enable RNNs to maintain an internal or hidden state that captures information about previously processed inputs, giving RNNs a notion of memory and allowing information to cycle within the network (Figure 3F). This mechanism enables RNNs to capture sequential and temporal dependencies in the input data. However, RNNs suffer from the vanishing gradient problem when trained on long sequences, which limits their ability to capture long-term dependencies. As a result, information from the early parts of a sequence may be lost by the time it reaches later steps. Gating mechanisms, implemented as special structures, were introduced in RNN architecture to address this problem. Standard RNNs update their hidden state at every step in the same way, causing early information to fade when sequences are long. Gates allow the network to decide what information to remember, forget, and output, enabling long-term memory. **Long Short-Term Memory (LSTM)** networks are gated RNNs consisting of a memory cell and three gates: input, forget, and output [53]. These gates control the flow of information into, out of, and within the memory cell, allowing it to retain important information over extended periods and enabling the network to effectively handle long-term dependencies in sequential data. LSTMs, often used in conjunction with other neural network architectures, have been applied to protein structure prediction and modeling [54].

### Attention mechanisms and Transformer Networks

The sequential nature of input processing in RNNs makes it difficult to access specific parts of the sequence and to capture long-range dependencies when generating outputs. Attention mechanisms alleviate this limitation by allowing RNNs to focus on different parts of the input sequence [55]. **Transformer networks**, which achieved state-of-the-art results in NLP tasks, introduced the self-attention mechanism, enabling them to dynamically access all parts of the input sequence simultaneously and effectively capture long-range dependencies [56]. Unlike the static filters of CNNs, self-attention dynamically computes filters based on the input, allowing Transformers to naturally process irregular and non-grid structured data. Unlike the attention used in RNNs, self-attention in Transformers analyzes all sequence elements in parallel and weighs their importance relative to each other, thus capturing both local and distant dependencies. In contrast to RNNs and LSTMs, Transformers enable parallel processing of sequential data during both training and inference, leading to significantly faster training times and making them well-suited for handling large-scale protein sequence datasets. Transformers generally adopt an encoder–decoder architecture, where the encoder applies self-attention to learn relationships within the input sequence, and the decoder uses attention mechanisms to generate the output sequence while incorporating information from the encoder. In biological applications, Transformers model long-range interactions in protein sequences essential for protein structure prediction tasks [57]. Originally designed for sequential data, Transformers have been adapted with domain-specific architectural modifications to process grid structured data (e.g., Vision Transformers [58]) and graph structured data (e.g., Graph Transformers [59]), making them highly versatile across multiple domains.

The **Swin-Conv-UNet** was developed to improve performance in semantic segmentation [60]. It is a hybrid neural network architecture combining CNNs and transformer-based attention mechanisms within a U-Net framework. Like U-Net, it uses an encoder–decoder structure with skip connections to capture multi-scale features and preserve spatial details (Figure 3D). Swin Transformer blocks are integrated to model long-range dependencies, allowing the network to capture both local and global contextual information. The convolutional components (**Conv**) learn fine-grained local features, while the transformer components (**Swin**) model complex global relationships across multiple scales (**UNet**), making Swin-Conv-UNet particularly effective for high-resolution and structured data tasks.

## 4 Deep learning-based automated model building in cryo-EM using a multi-modal approach

Deep learning methods for model building in cryo-EM density maps differ in the underlying neural network architecture, prediction task, target biomolecules, and the level of structural detail in the resulting models (Tables 2,3). However, most approaches follow a common multi-stage pipeline that transforms raw cryo-EM density maps into molecular models. Briefly, the representative steps are (**Figure 4**): (1) preprocessing the input density map; (2) applying deep neural networks to learn features from the preprocessed density map that identify different aspects of a biomolecular structure; and (3) model building from the learned features followed by refinement.

**Figure 4.**
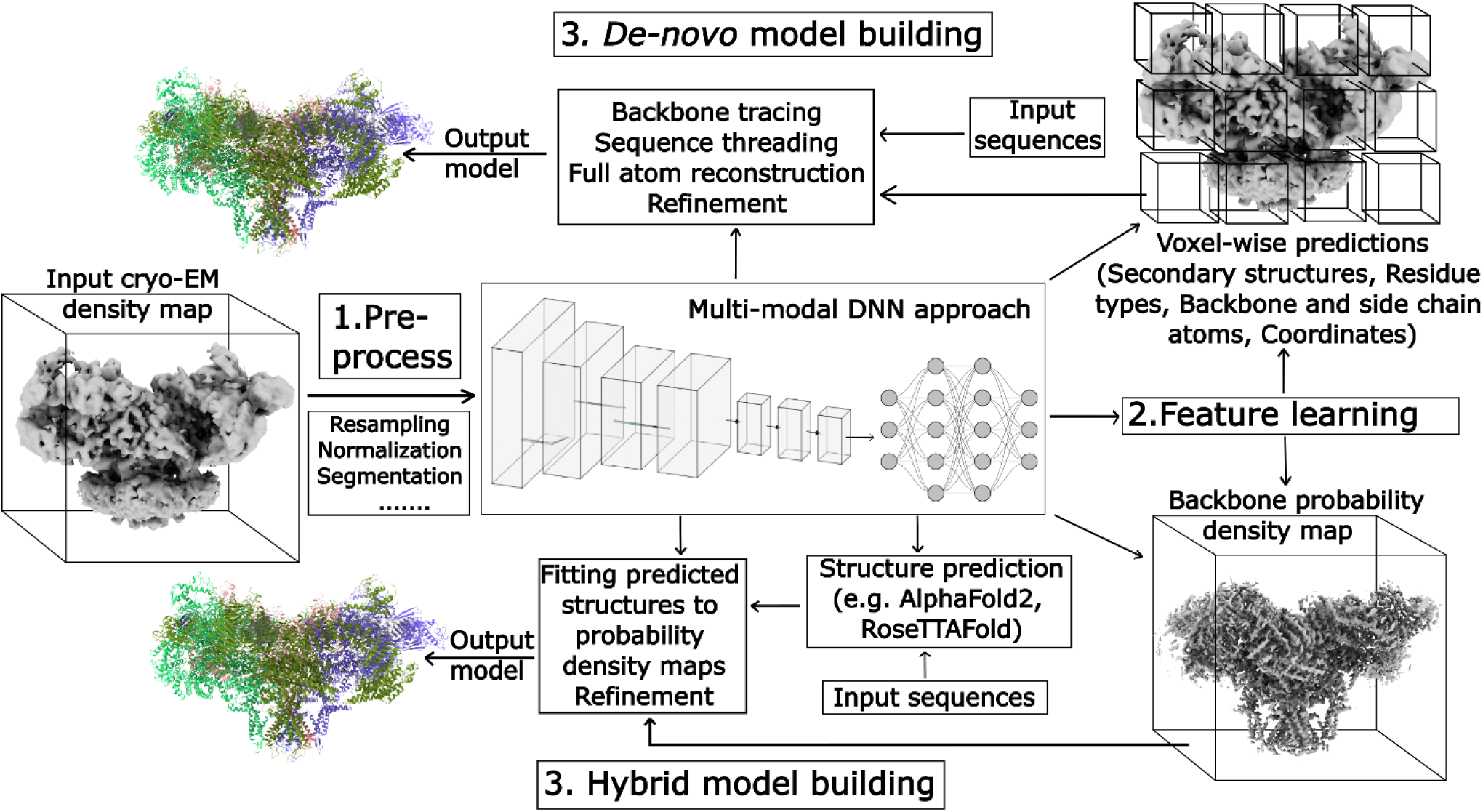
Schematic overview of the representative steps used by deep learning-based tools for automated model building in cryo-EM, using a density map and the model of ryanodine receptor 1 as input and output, respectively. Illustrations of the deep neural networks (DNNs) are generated using NN-SVG [61].

### Training datasets and preprocessing

Deep neural networks underlying automated model building methods in cryo-EM are trained and tested on large, labeled datasets comprising experimental and/or simulated density maps (Table 4). Since few experimental cryo-EM density maps were available in the EMDB [18] during the early years (Figure 1), the first deep learning-based methods for cryo-EM model building relied on simulated or synthetic maps, generated from PDB structures, for training and testing (Table 4). Utilities such as *pdb2mrc*, *pdb2vol*, or *molmap* from the EMAN2 package [62], the Situs package [63], or UCSF Chimera [64] can be used to simulate cryo-EM density maps from PDB models at different resolutions. As the cryo-EM resolution revolution increased the number of experimental cryo-EM density maps in the EMDB (Figure 1), subsequent deep learning-based methods for cryo-EM model building began to utilize these maps for training and testing (Table 4). Simulated maps are still useful in cases where experimental density maps do not provide enough data to train deep learning models for a specific model-building task. Therefore, many cryo-EM model-building methods combine simulated and experimental density maps to train their deep neural networks (Table 4). Before training, the input cryo-EM density maps are first preprocessed using resampling and normalization. Resampling involves standardizing the varying voxel sizes of raw density maps to a uniform size. The resampled maps are then normalized to make density values comparable across different maps. Alternatively, publicly available datasets such as Cryo2Struct2Data [65] can be utilized. This dataset is a significant resource comprising 7,600 preprocessed cryo-EM density maps, in which voxels are labeled based on known atomic structures, making it suitable for training and testing deep learning-based model building methods in cryo-EM.

### Feature learning

During training, deep neural networks automatically learn hierarchical, increasingly abstract representations from preprocessed cryo-EM density maps, enabling them to identify key structural features for model building. Specifically, deep neural networks learn voxel-wise representations from preprocessed cryo-EM density maps that predict multiple structural features such as backbone atom positions, residue types, and secondary structures at each voxel, offering a preliminary, coarse-grained representation of the molecular structure that can guide subsequent model construction. Deep learning-based automated model building methods employ different neural network architectures for feature learning (Table 2), with specific details provided for each method in the following sections.

**Table 2.**
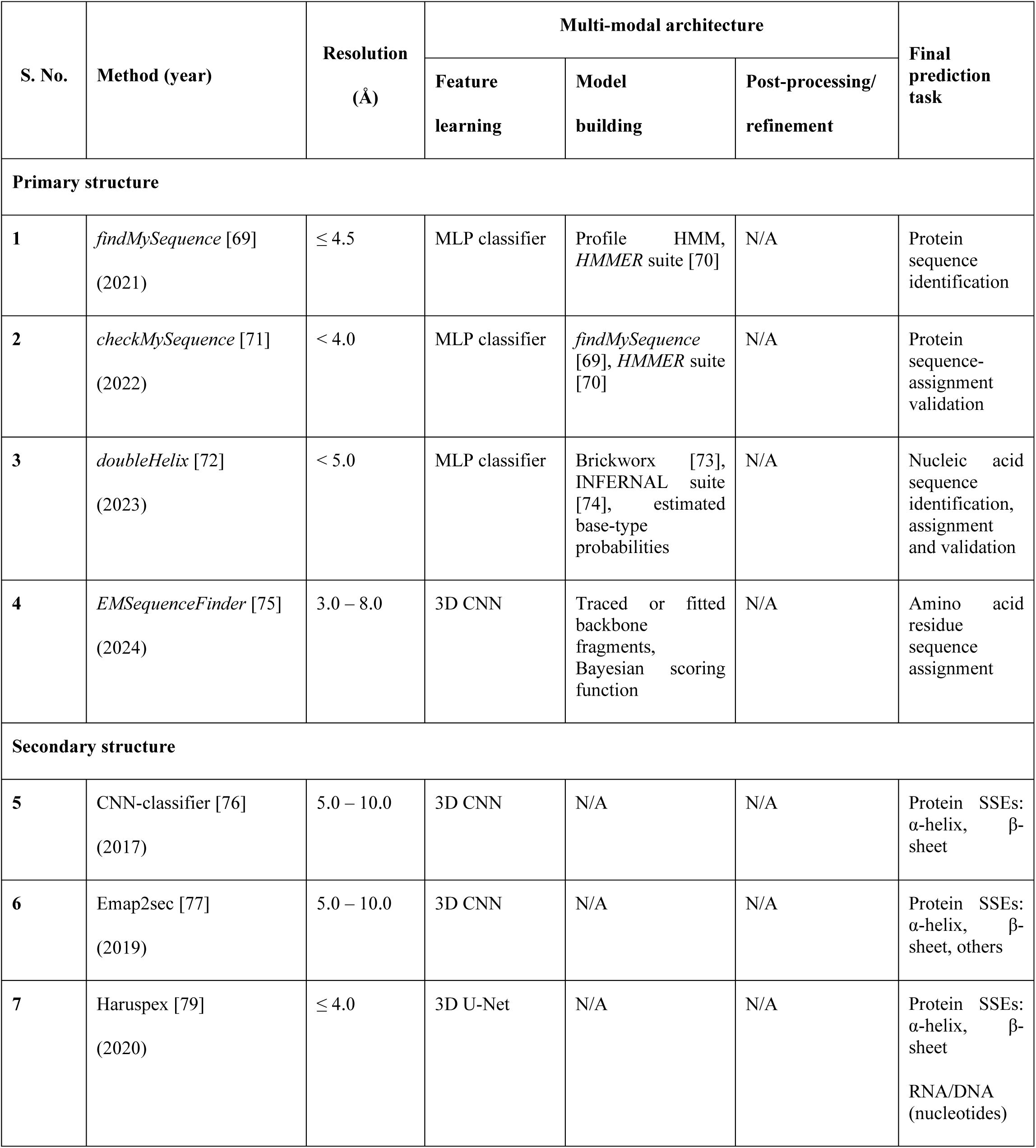
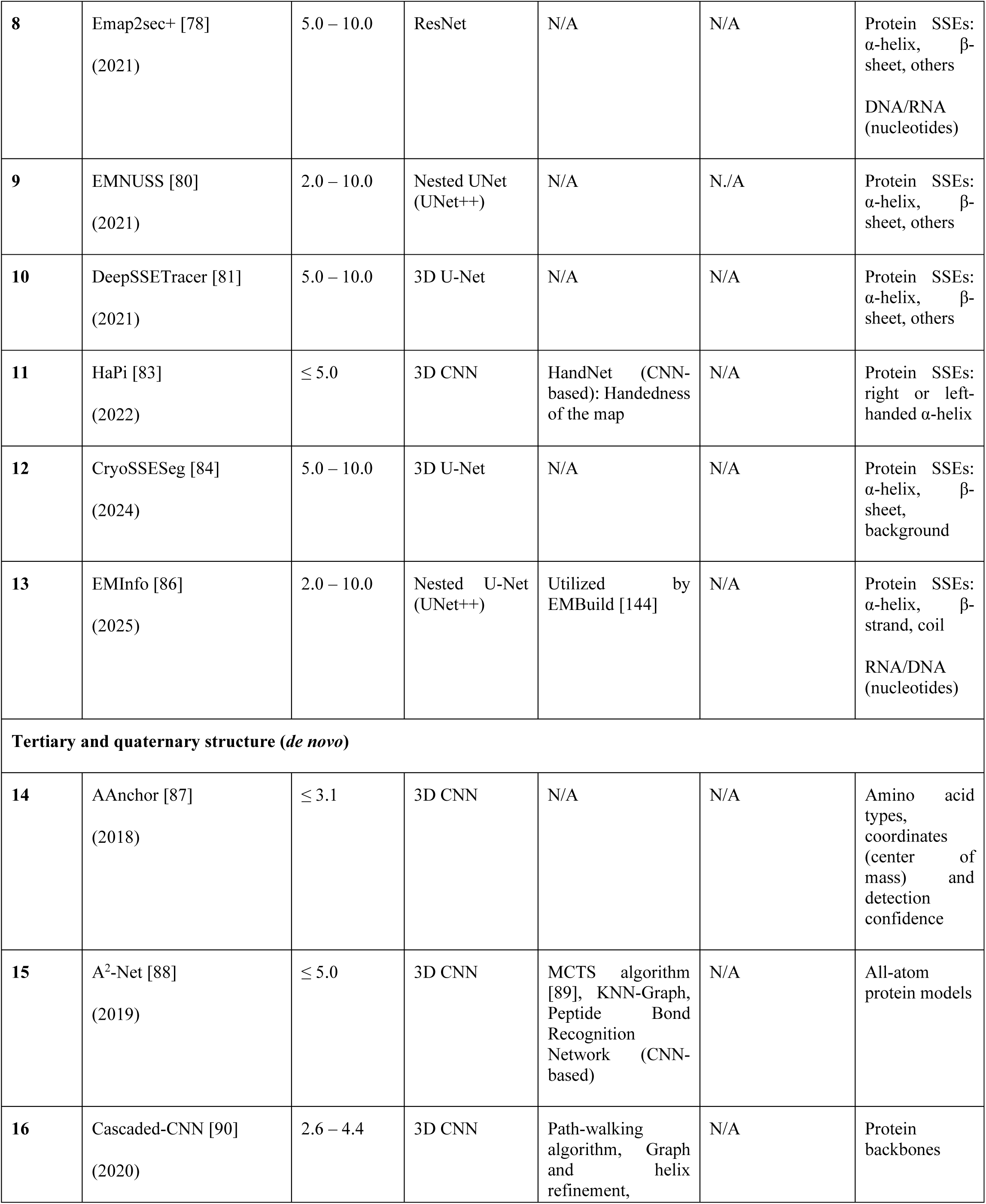
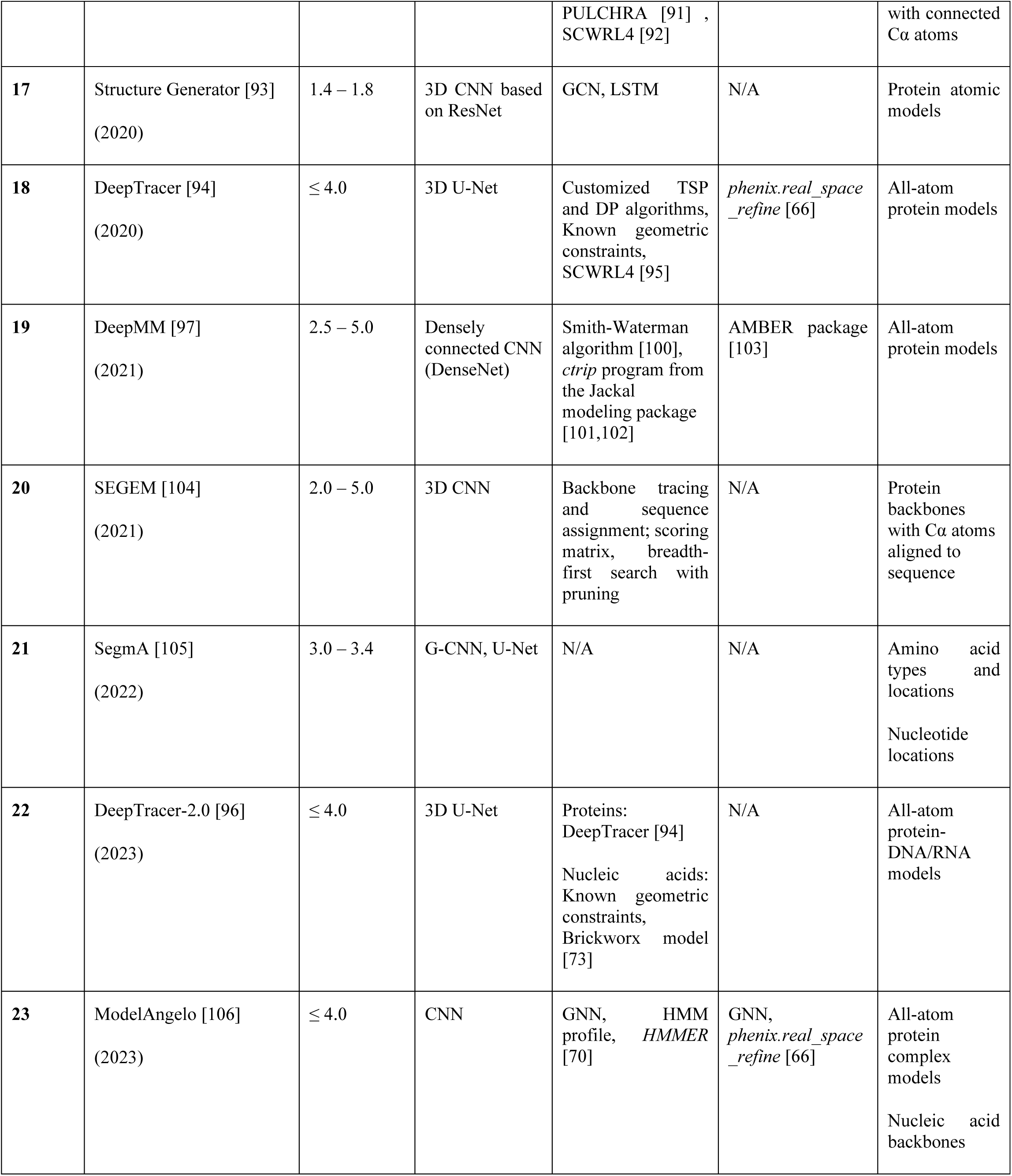
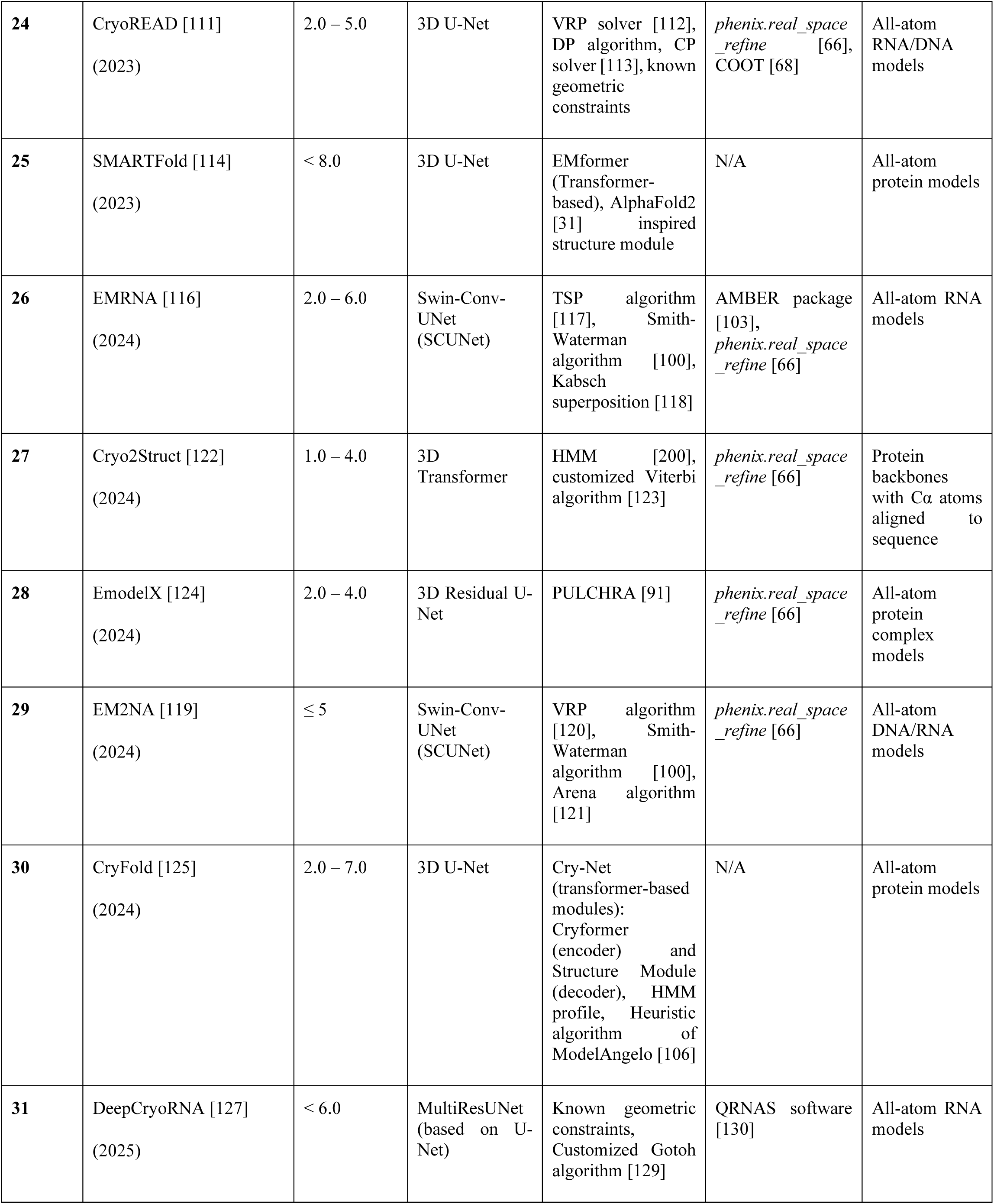
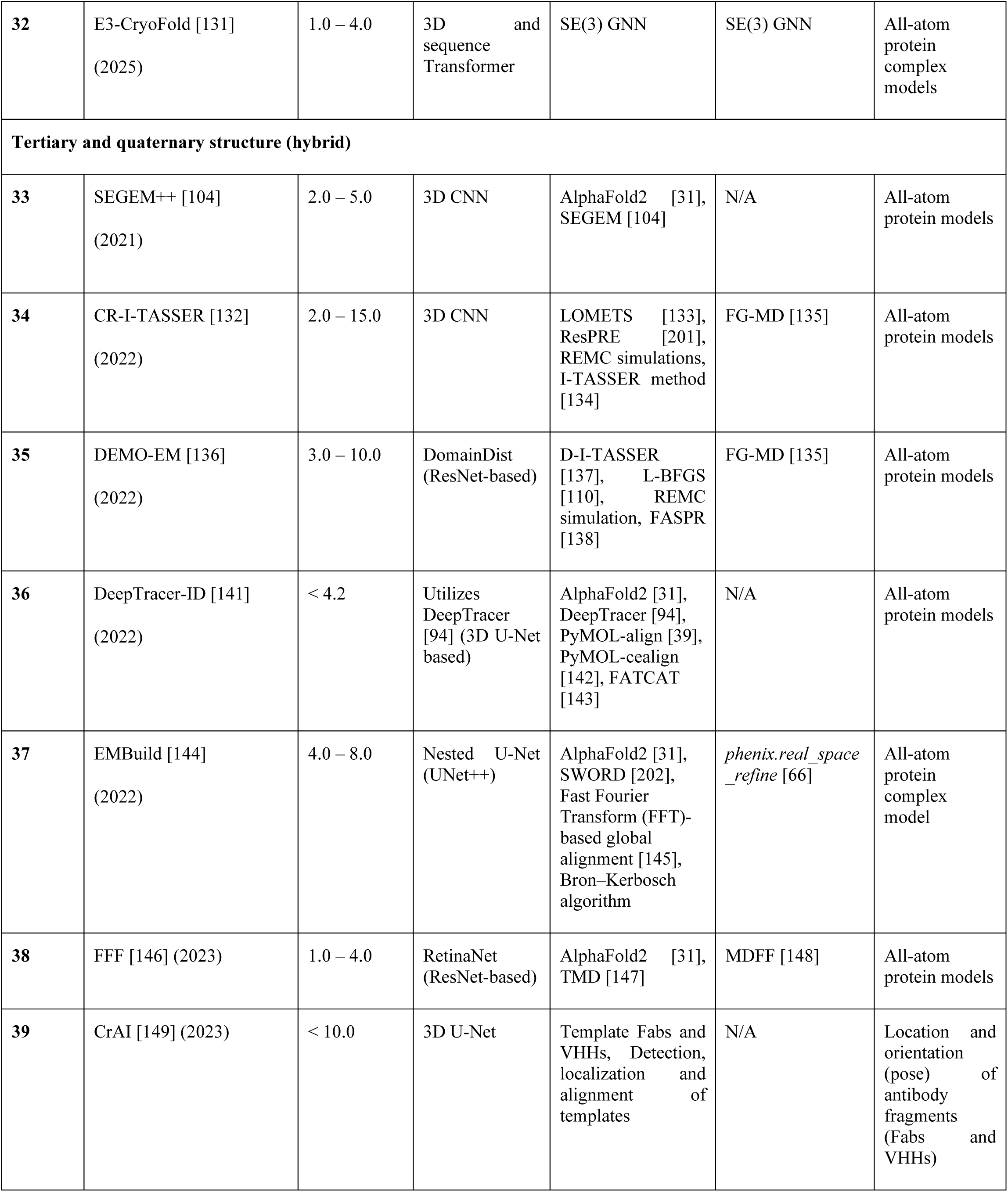
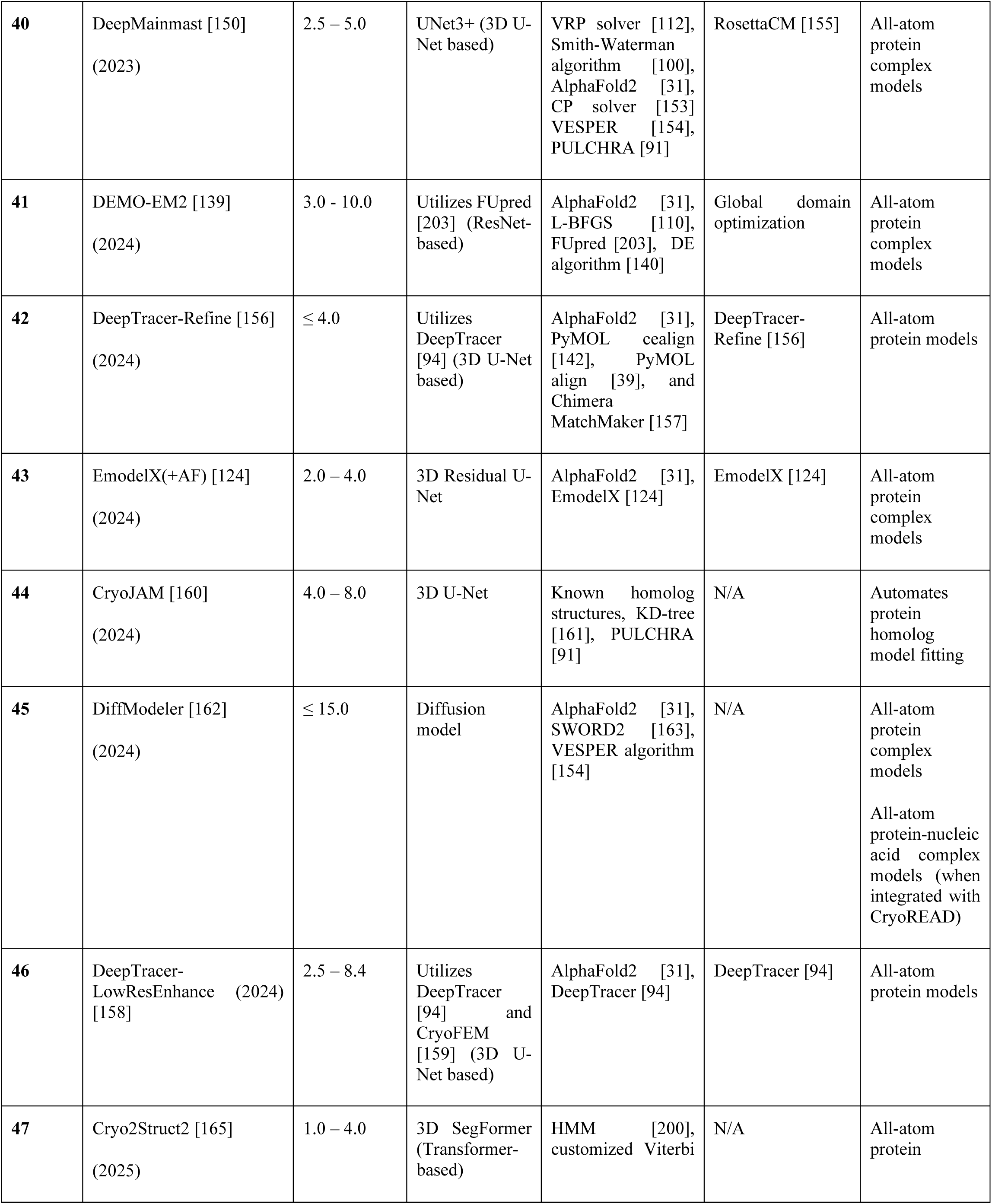
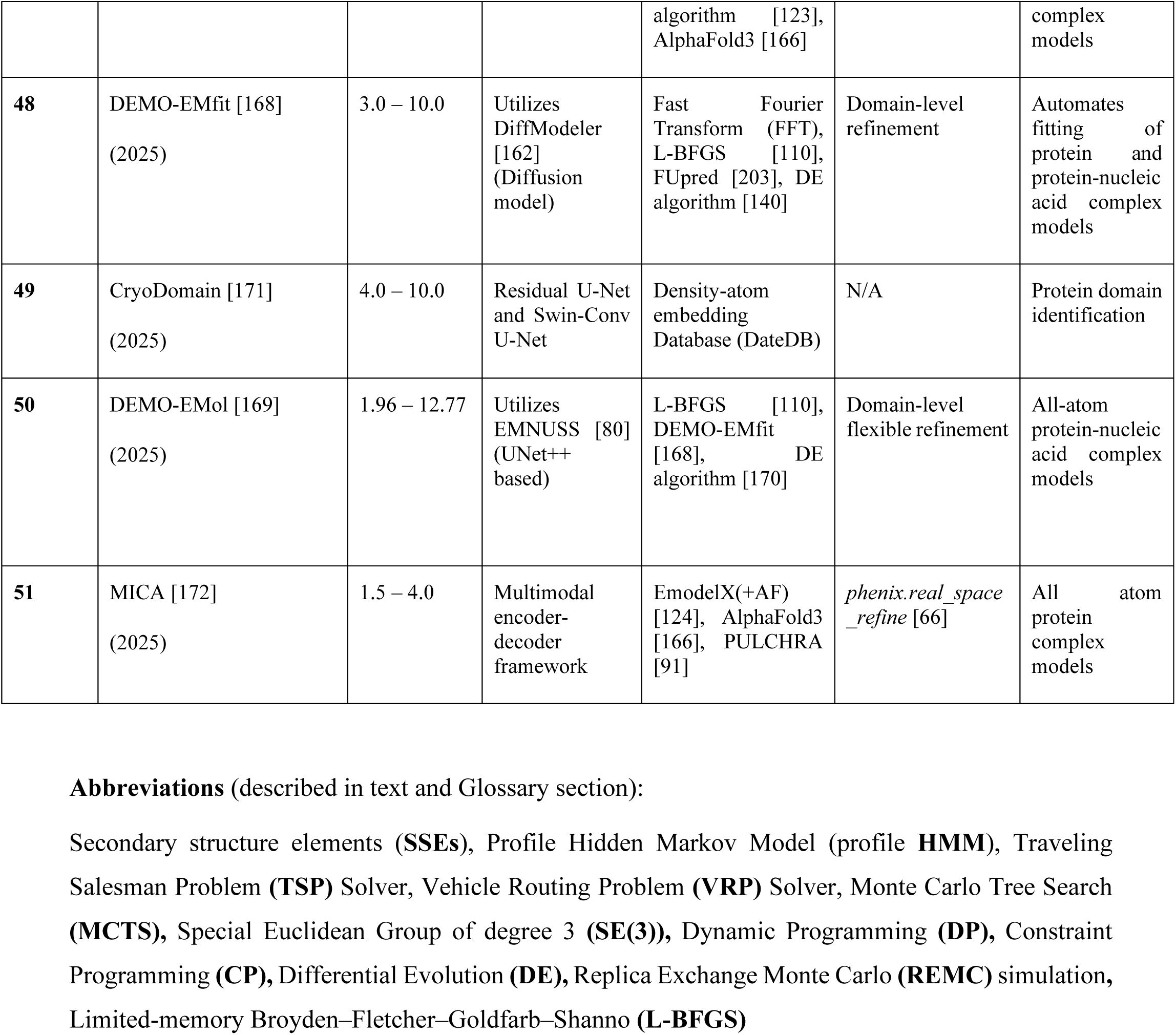
Architecture of deep learning-based automated model building methods in cryo-EM.

### Model building

Since the voxel-wise representations learned by deep neural networks from preprocessed cryo-EM density maps provide only a coarse structural representation, they must be further processed to generate a complete atomic model. This broadly involves backbone tracing, sequence assignment, and full-atom reconstruction. The learned voxel representations serve as a basis for constructing an initial structural model. Many methods represent this as a graph where nodes correspond to residues, secondary structure elements, or small structural fragments, while edges denote spatial proximity or potential chemical bonds, which is then iteratively processed and refined. Backbone tracing generates chains and fragments by connecting predicted backbone atoms while incorporating stereochemical constraints. This is a complex and challenging step due to the factorial growth in the number of ways spatially distributed atoms can be connected, requiring the use of advanced optimization algorithms. Sequence assignment involves correctly threading the biomolecular sequence onto the traced backbone structures, followed by full-atom reconstruction including side chains. Details of the model building step for each method are summarized in Table 2 and described in the following sections.

### Refinement

Deep learning–based automated model-building methods often employ established structural biology tools, including *phenix.real_space_refine* [66], molecular mechanics force fields, molecular dynamics flexible fitting (MDFF) [67] and COOT [68], to further refine reconstructed atomic models. These tools utilize energy minimization techniques, apply physical and stereochemical constraints, and incorporate empirical knowledge-based restraints to refine models, thereby ensuring that the biomacromolecule retains correct chemical, stereochemical, and geometric properties including realistic bond lengths and angles as well as minimal steric clashes. The application of *phenix.real_space_refine* [66] significantly improves side-chain conformations and map–model correlations. In some workflows, this step is followed by additional refinement using molecular dynamics (MD) simulations or energy minimization algorithms, which further optimize the atomic arrangement by simulating physical forces and interactions.

### Deep learning-based automated model building methods

Biomacromolecules exhibit hierarchical levels of structural complexity and organization, ranging from the linear sequence of ordered residues to the complex three-dimensional spatial arrangement of atoms. (**Section 2**). In this section, we group deep learning-based automated model-building methods by their ability to predict **(4.1)** primary, **(4.2)** secondary, and **(4.3)** tertiary or quaternary structural aspects of biomacromolecules. Deep learning tools for building atomic models in cryo-EM density maps are further grouped as *de novo* **(4.3.1)**, where the model is predicted directly from features learned from the cryo-EM density, or hybrid **(4.3.2)**, where it is derived by integrating structural templates with these features. It should be noted that methods grouped under tertiary or quaternary structure also extract structural features at the secondary (such as α-helices and β-sheets) or primary structure level, but their ultimate prediction task is to determine the three-dimensional arrangement of residues and atoms. Methods grouped under secondary structure, in contrast, focus only on predicting secondary structure classes from the input cryo-EM density maps.

4.1 **<u>Primary structure prediction:</u>** The primary structure of proteins and nucleic acids is essentially their linear sequence of residues covalently linked to each other along the polymer chain. Neural networks can predict residue type probabilities from cryo-EM reconstructions, enabling the assignment and validation of protein or nucleic acid sequences. Tools specifically designed for this task are described below.

***findMySequence*** is a computer program that identifies protein sequences from cryo-EM reconstructions and crystallographic data [69]. To achieve this, it uses machine-learning predicted residue-type probabilities to query sequence databases using *HMMER* suite [70]. To predict residue-type probabilities from cryo-EM map and main-chain models, *findMySequence* utilizes a neural network with two fully connected hidden layers offering improved performance over a previous support-vector machine-based classifier. To identify a sequence that matches the predicted residue-type probabilities, the probabilities are first converted into a multiple sequence alignment (MSA), then into a profile Hidden Markov Model (profile-HMM) using the *HMMER* suite. This profile-HMM is then used to query a sequence database to find plausible matches (**Box 1**: **Glossary section**). To build side chains for an input main-chain model fragment, *findMySequence* considers all possible alignments of the fragment to the target sequence and selects the most plausible one based on the predicted residue-type probabilities to assign residue types to the fragment.

***checkMySequence*** is a new, automated tool designed to quickly and accurately detect register shifts in protein models built into cryo-EM density maps including large macromolecular complexes [71]. To detect register shifts, *checkMySequence* first identifies a reference sequence for each protein chain in the input model. This step uses a protocol from the *findMySequence* program [69], employing a neural network classifier to generate residue-type probability profiles from the cryo-EM density map and input model. These probabilities are then used with the *HMMER* suite [70] to query sequence databases, scoring plausible matches. Once reference sequences are established, *checkMySequence* assigns fragments of the input model to them, identifying areas where this new assignment conflicts with the original sequence assignment in the input model. If a discrepancy is found, the method suggests a more plausible sequence assignment.

***doubleHelix*** is a computer program designed for the assignment, identification, and validation of nucleic acid sequences in structures determined using both cryo-EM and X-ray crystallography [72]. *doubleHelix* combines neural network classifiers to identify nucleobases with a sequence-independent secondary structure assignment approach for comprehensive nucleic acid analysis. The neural network classifiers estimate the likelihood of a nucleotide being a purine or a pyrimidine based on a provided backbone model and its corresponding density map. Each neural network classifier is built with two fully connected hidden layers. After estimating purine and pyrimidine probabilities using the neural network, *doubleHelix* then identifies base pairs in RNA and DNA models by aligning recurring nucleic acid structural motifs to the target model with base-pairing restraints derived from the backbone conformation. This process involves superimposing small search-fragments of known secondary structures onto the input model using an algorithm from the *Brickworx* program [73]. For sequence identification, *doubleHelix* leverages the INFERNAL suite [74] to find the most plausible sequence in a database, using the input model’s residue-type probabilities and secondary structure. Similarly, it uses these estimated base-type probabilities to assign RNA or DNA models to a target sequence.

***EMSequenceFinder*** is a deep learning method that assigns amino acid sequences to backbone fragments traced in cryo-EM density maps [75]. The method requires a cryo-EM density map with a resolution better than 8.0 Å, the corresponding backbone traces fitted or traced into the map (specifying N, Cα, C, and O atom positions), and one or more protein sequences. *EMSequenceFinder* quantifies the density map fit of traces using a 3D Convolutional Neural Network (CNN), trained on a large dataset of cryo-EM density maps. The network determines the likelihood of each residue type by extracting features from the side chain voxel intensities. These extracted features are then combined with additional input, such as map resolution and the secondary structure type of the trace containing the residue, to output the probability for each amino acid. *EMSequenceFinder* then assigns protein sequences to backbone fragments by identifying the best-scoring threading among all possible alignments of a sequence to the backbone trace. It achieves this using a Bayesian scoring function that ranks the 20 standard amino acid types at each backbone position, considering the fit to the density map (quantified above using a 3D CNN), map resolution, and secondary structure propensity. The output is the best-scoring threading for each input backbone trace.

4.2 **<u>Secondary structure prediction:</u>** Secondary structure refers to local, stable, and regular conformations of the polymer backbone, such as α-helices and β-sheets in proteins, and regular helical segments in nucleic acids. Deep learning tools specifically designed to predict voxel-wise probabilities of secondary structure types from the cryo-EM density maps are described below in an approximate chronological order.

**CNN-classifier** is a 3D convolutional neural network (3D CNN) designed to automatically detect secondary structures of proteins in cryo-EM density maps [76]. It predicts the probability of a voxel belonging to specific protein secondary structure elements (SSEs), such as α-helices, β-sheets, or background. By using 3D convolutions, the model effectively captures 3D spatial information within the protein structures, learning discriminative features automatically from cryo-EM density maps. To enhance efficiency, the CNN-classifier incorporates multiple deconvolution operations at various intermediate layers to generate feature maps of same dimension, which were then summed to create a multi-scale representation. The CNN-classifier combines inception learning and residual learning with dilated convolutions. The inception component uses multiple convolutional layers with different filter sizes to create diverse paths between network hidden layers. This increases the number of trainable parameters at each step without increasing the overall computational complexity. Residual learning employs shortcut identity mappings to simulate nonlinear relationships between input and output layers allowing the network to achieve accurate results without added computational cost. Dilated convolutions, integrated at the reception module, expand the network’s receptive field, allowing it to capture information across various scales without losing image resolution.

**Emap2sec** is a deep learning method designed to identify protein secondary structures (α-helices, β-sheets, and other structures) directly from cryo-EM density maps with resolutions between 5 and 10 Å [77]. It employs a 3D convolutional neural network (3D CNN) to assign a secondary structure to each grid point. The core of Emap2sec is a two-phase stacked neural network where phase 1 network analyzes the normalized density of a single voxel and outputs probability values for the three secondary structure classes (α-helix, β-sheet and other structure; defined as structure that is not α-helix nor β-sheet). Phase 2 network then refines these initial predictions by incorporating contextual information from neighboring voxels. Ultimately, each voxel is assigned to the secondary structure class with the highest probability among the three types.

**Emap2sec+**, successor of Emap2sec, is designed to identify DNA or RNA in addition to protein secondary structures directly from cryo-EM maps at 5-10 Å resolution [78]. It employs a deep residual convolutional neural network (ResNet), framing the task as a classification problem rather than segmentation due to prior success with Emap2sec and that classification would perform better than segmentation at intermediate resolutions. Emap2sec+ classifies individual voxels from cryo-EM density maps into one of four structural categories - DNA/RNA or a protein secondary structure (α-helices, β-sheets, other structures). This classification occurs through a two-phase neural network. In Phase 1, an input voxel undergoes five independent evaluations: four binary classifiers each determining the probability of a specific structure at the voxel (DNA/RNA, α-helix, β-sheet or other structures), and a fifth multi-class classifier providing probabilities for all four categories. The probability values from these Phase 1 classifiers are then fed into the Phase 2 network which refines the final probability assignments of the four structure classes for the central voxel by considering the structural predictions of neighboring voxels, which effectively smooths the structural assignment across the entire cryo-EM density map.

**Haruspex** is a deep learning tool that leverages convolutional neural networks to automatically identify and annotate protein secondary structure elements and RNA/DNA within cryo-EM density maps at an average map resolution of 4.0 Å or better [79]. Haruspex utilizes a U-Net-style convolutional neural network. It processes voxel segments through multiple convolutional and pooling layers and extracts features to identify different structural elements, which are later combined to restore spatial detail. The final output provides a probability for each voxel, indicating whether it belongs to an α-helix, β-strand, nucleotide, or is unassigned. This essentially annotates the cryo-EM density map with the locations of these key biomolecular structures.

**EMNUSS** utilizes a nested 3D U-Net architecture (UNet++) to annotate secondary structures in cryo-EM density maps, effective at both intermediate and high resolutions [80]. This design, which starts with an encoder subnetwork followed by a decoder subnetwork, concatenates the outputs of these subnetworks, after which a final convolution layer assigns secondary structure classifications. For each voxel in the annotated region, EMNUSS provides three channels containing probabilities that indicate whether the voxel is close to an α-helix residue, a β-strand residue, or a coil residue. Compared to the U-Net architecture, UNet++ incorporates dense skip connections on its skip pathways, which significantly enhances gradient flow.

**DeepSSETracer** employs a neural network architecture of end-to-end convolution operations, adapted from the 3D U-Net architecture [46], to detect secondary structures within cryo-EM component maps for individual chains at medium resolution, rather than analyzing an entire cryo-EM density map, which may contain multiple chains. [81]. This architecture allows DeepSSETracer to process density maps of varying sizes, predicting the probability of each voxel belonging to an α-helix, β-sheet, or other structure. The tool is integrated with ChimeraX [82] in a software bundle that integrates the secondary structure prediction with visualization capabilities of ChimeraX.

Cryo-EM reconstruction can yield two equally consistent, but mirror-image, 3D density maps. Since proteins have a specific handedness, only one reconstruction is correct. Currently, biologists must manually determine this by inspecting the rotations of α-helices within the map, a task that becomes challenging at lower resolutions as helices lose their distinct handedness. **HaPi** (Handedness Pipeline), uses two 3D convolutional neural networks (CNNs) to automatically determine handedness of the cryo-EM density map for resolutions up to 5.0 Å [83]. The first network, *AlphaVolNet*, identifies the location of α-helices throughout the entire map by processing each non-background voxel. The second network, *HandNet*, then evaluates the overall handedness of the map using a consensus strategy that averages the individual handedness predictions.

**CryoSSESeg** is a convolutional neural network (CNN) framework designed to identify the organization of protein secondary structure elements within medium-resolution cryo-EM density maps [84]. CryoSSESeg works by first isolating individual protein chains from an atomic model and then using their coordinates to extract and mask the corresponding regions in the cryo-EM density map. This chain-based cropping ensures that entire secondary structures are present within the image, which helps the model learn more effectively. The STRIDE secondary structure annotation tool [85] is then used to label individual voxels within the density map. The network is an adaptation of the 3D U-Net model, featuring a downsampling [86] path (contracting path) that condenses spatial information, a bottleneck layer that further compresses data and enhances feature representation, and an upsampling path (expansive path) that recovers spatial detail. The final layer outputs three channels, each corresponding to one of three classes: α-helix, β-sheet, or background.

**EMInfo** is a deep learning method that uses a 3D UNet++ architecture to automatically detect protein secondary structures and nucleic acid locations in cryo-EM density maps [86]. UNet++ is a supervised encoder-decoder network comprising of downsampling, upsampling, and skip connections. This design allows UNet++ to implement multi-scale feature extraction from input maps without significantly increasing computational cost. EMInfo outputs a four-channel probability for each voxel representing the likelihood of it belonging to different structural categories. For each voxel, the channel with the highest probability is selected as its predicted category, ultimately generating a structure annotation map that directly corresponds to the input density map. EMInfo achieves accurate structural annotation by ignoring background voxels, those with a density value below a specified contour level, ensuring that structural regions are not misclassified as background.

4.3 **<u>Tertiary and quaternary structure prediction:</u>** Tertiary and quaternary structures refer to the three-dimensional arrangement of atoms in a single polymer chain or multiple chains (subunits), respectively. This structural level captures interactions between residues that may be far apart in the sequence. Deep learning tools specifically designed to build 3D atomic models in the cryo-EM density map are discussed below. They are further grouped as *de novo* **(4.3.1)**, where the model is predicted directly from features learned from the cryo-EM density, or hybrid **(4.3.2)**, where it is derived by integrating structural templates with these features.

**4.3.1 *De novo* model building:** In this approach, deep learning tools perform model building directly from voxel-wise backbone (and sometimes sidechain) atom, secondary structure and residue type probabilities learned from the cryo-EM density map to generate 3D atomic models.

**AAnchor** (amino-acid anchor) is a deep learning method designed to identify and precisely locate high-confidence anchor amino acid residues in high-resolution cryo-EM density maps (at resolutions 3.1 Å or better) [87]. AAnchor uses a detection algorithm to first generate candidate locations for amino acids, filtering out those likely to be inaccurate and a classification convolutional neural network (CNN) to classify these candidate volume cubes into one of 21 labels (20 amino acids plus none). The detected anchor residues can be crucial for various local *de novo* modeling tasks, including accurately positioning secondary structures, modeling loops, and facilitating general fragment-based modeling.

*<u>Feature learning</u>***: A^2^-Net** takes a cryo-EM density map and a protein sequence as input and determines the 3D structure of the protein [88]. The method employs deep convolutional neural networks (CNNs) to detect amino acids by learning the conformational densities of individual amino acids within the density volume. To further enhance detection, prior knowledge of protein sequences is incorporated by designing a sequence-guided neighbor loss during training. A^2^-Net processes 3D density volumes to generate 3D feature volumes and uses localization network (locNet) and recognition network (recNet) to detect and classify amino acids in 3D space and estimate its pose, which determines the 3D coordinates of atoms in each amino acid. In locNet, a 3D Region Proposal Network (RPN) uses 3D convolutional layers to propose amino acid locations. The RPN classifies valid proposals and estimates their coordinates. In recNet, amino acid proposals are then fed to a new Aspect-ratio Preserved Region of Interest (APRoI) layer (a specialized type of pooling layer), which extracts Regions of Interest (RoI) into fixed cubic which are further processed by 3D convolutional layers to predict its amino acid category. For atomic-level detail, PoseNet, a 3D stacked hourglass network (a type of fully convolutional neural network designed for volumetric data, structured in an encoder–decoder style with skip connections), regresses the 3D coordinates of each atom in amino acids, completing the pose estimation. *<u>Model building</u>*: To construct protein main chains and thus the full molecular structure, **A^2^-Net** uses Monte Carlo Tree Search (MCTS) algorithm [89] (**Box 1**: **Glossary section**) which iteratively builds a search tree to effectively search and thread candidate amino acid proposals. To make MCTS search computationally feasible with many proposals, a K-nearest neighbors (KNN) graph is created which connects spatially close amino acid proposals, narrowing the search space for the MCTS. This search process is further optimized by implementing tree pruning which incorporates a convolutional neural network (CNN)-based Peptide Bond Recognition Network (PBNet) to predict peptide bonds between proposals which reduces the edges in the KNN-graph.

*<u>Feature learning</u>***: Cascaded-CNN** (C-CNN) is a model building tool that comprises of a series of convolutional neural networks (CNNs), each predicting a specific component of protein structure from the input cryo-EM density maps [90]. The overall goal of C-CNN is to accurately predict Cα atom locations using information from the intermediate secondary structure elements (SSEs) and backbone predictions. C-CNN leverages the fully connected network design and dilated convolutions to classify a full 3D image in a single pass. Dilated convolutional layers increase the receptive field while preserving the size of the input image. C-CNN comprises of three feedforward neural networks, taking cryo-EM density maps as input. SSE CNN predicts α-helices, β-sheets, or loops/turns for each voxel and output a confidence map for each SSE. These three SSE maps along with input density maps serve as input to the Backbone CNN to predict whether each voxel is part of the backbone structure of the protein or not and outputs two backbone confidence maps. Cα-Atom CNN then predicts Cα atom locations by taking all the previous maps - input density map, SSE and backbone confidence maps - and classifies if a voxel is part of a Cα atom or not producing two Cα atom confidence maps. Each voxel receives a confidence value, with its final classification determined by the highest confidence among the output maps. *<u>Model building</u>:* The confidence maps from the **C-CNN** are post-processed using a path-walking technique to generate protein backbone structure with precise Cα atom locations. The path-walking algorithm navigates high-confidence backbone regions, connecting Cα atoms using a novel tabu-search algorithm that scores movements based on the location’s local density prediction confidence and distance, and incorporates backbone torsion angles and known geometrical parameters of secondary structures as weights. The algorithm continues tracing until no suitable Cα atoms can be found or previously processed areas are encountered. The output is a PDB file of disconnected Cα atom traces. The disconnected traces, representing partial protein backbones, are further refined through path combination and backbone refinement. First, the traces are converted into a graph where Cα atoms are nodes and connections are edges. The path combination step then merges these disjoint graphs into a single, fully connected representation of the protein’s backbone. After this, a backbone refinement step removes false-positive connections, leaving only accurate Cα node and edge connections. Further improvements to the α-helix SSEs of the backbone trace are done using a helix-refinement algorithm. Finally, a novel quality assessment-based combinatorial algorithm is used to map protein sequences onto the reconstructed Cα traces, generating full-atom protein structures. This algorithm also reconstructs side-chain atoms using PULCHRA [91] and SCWRL4 [92], based on the Cα coordinates of each segment.

*<u>Feature learning</u>***: Structure Generator** is a fully automated, template-free deep learning method for protein model building in cryo-EM density maps [93]. It uses RotamerNet, a 3D convolutional neural network (CNN) built on the ResNet architecture, to output the predicted amino acid and rotamer identity, along with the proposed coordinates for its Cα atom. RotamerNet analyzes the density profiles to propose a set of candidate amino acids and their 3D locations, unconstrained by the known protein sequence. *<u>Model building</u>:* **Structure Generator** uses a graph convolutional network (GCN) to create an embedding from initial rotamer-based amino acid identities and predicted candidate 3D Cα locations. Following this, a bidirectional long short-term memory (LSTM) module processes this embedding to order and label the candidate identities and atomic positions, ensuring consistency with the input protein sequence to ultimately generate a structural model. The graph convolutional network (GCN) efficiently encodes the output from RotamerNet as a graph, while a bidirectional Long Short-Term Memory (LSTM) module accurately decodes this information to generate a directed amino acid chain.

*<u>Feature learning</u>***: DeepTracer** is a fully automated deep learning method designed for the rapid *de novo* structure determination of multi-chain proteins directly from high-resolution cryo-EM density maps [94]. Its core is a specialized convolutional neural network (CNN) made up of four connected U-Nets. From the preprocessed cryo-EM density maps, each U-Net predicts a distinct aspect of the protein structure (atoms, backbone, secondary structure elements, and amino acid types). The Atoms U-Net predicts if a voxel contains either a Cα atom, a nitrogen (N) atom, a carbon (C) atom, or no atom (four output channels). The Backbone U-Net determines if each voxel belongs to the protein backbone, side chains, or non-protein regions (three output channels). The Secondary Structure U-Net recognizes loops, α-helices, β-sheets and no structure (four output channels). The Amino Acid Type U-Net determines the specific type of amino acid at each voxel (21 output channels, 20 standard amino acids plus no amino acid). *<u>Model building</u>:* To create an initial model structure, **DeepTracer** first determines disconnected chains using the output of the Backbone U-Net by identifying connected areas of backbone voxels, with each disconnected area being designated as a separate chain. It then calculates the precise 3D coordinates of each Cα atom using output Cα channels of the Atoms U-Net. For connecting the Cα atoms into continuous chains, DeepTracer uses a modified traveling salesman problem (TSP) algorithm (**Box 1**: **Glossary section**). Instead of simply minimizing distance (as in a traditional TSP), DeepTracer uses a custom confidence function to determine the likelihood of two atoms being connected and the overall goal of the TSP algorithm becomes maximizing the sum of these confidence scores. DeepTracer then uses a custom dynamic programming (DP) algorithm (**Box 1**: **Glossary section**) for protein sequence alignment, aligning segments of the predicted amino acid sequence with the known amino acid sequence of the target protein. Based on the alignment, the initially predicted amino acid types are updated for greater accuracy. DeepTracer then builds the complete protein backbone by adding carbon (C) and nitrogen (N) atoms to the previously placed Cα atoms. For this task, it uses U-Net provided confidence maps for C and N atoms and applies molecular mechanics principles specific to peptide chains including planar peptide geometry, to ensure chemically accurate placement. At the final step, DeepTracer predicts sidechains, aiming to accurately position the side-chain atoms for each amino acid. This is achieved using SCWRL4 [95], an automated tool that takes the complete protein backbone and amino acid types as input and outputs a sterically plausible protein structure.

*<u>Feature learning</u>***: DeepTracer-2.0** enhances the capabilities of DeepTracer by incorporating the identification of nucleic acids alongside amino acids from cryo-EM density maps [96]. DeepTracer-2.0 achieves this through an initial segmentation step that separates the cryo-EM map into distinct macromolecular densities. Following this, the pipeline employs two specialized U-Net architectures: an amino acid U-Net for protein backbone and Cα atom determination, and a newly integrated nucleotide U-Net for identifying phosphate (P) and sugar carbon atoms in nucleic acid regions. This nucleotide U-Net, distinct from the amino acid U-Net due to the differing molecular structures of proteins and nucleic acids, predicts the structural aspects of nucleic acids. The Atoms U-Net, with four output channels, identifies whether each input voxel contains a phosphate (P), sugar carbon atoms - C1’, C4’ or no atoms. Concurrently, its Backbone U-Net, with three output channels, determines if a voxel belongs to the sugar-phosphate backbone, the nitrogenous base, or neither. Both U-Nets are optimized for defining the DNA/RNA phosphate backbone. *<u>Model building</u>***: DeepTracer-2.0** post-processing phase refines the predictions from nucleotide U-Net, the predicted phosphate (P) and carbon atom (C1’, C4’) positions, to build an accurate sugar-phosphate backbone consistent with DNA/RNA biological principles. This involves reducing spurious phosphate predictions and connecting the remaining ones based on characteristic DNA/RNA geometry, considering the influence of sugar puckers, and utilizing pseudotorsion angles to simplify backbone construction. Finally, the refined phosphate atoms and cryo-EM density map data are fed to Brickworx model [73], which completes the nucleotide modeling by identifying matching double-stranded helical motifs for DNA or extending to recurrent RNA motifs, including single-stranded segments, ensuring the final structure adheres to known nucleic acid conformations. The nucleotide post-processing step allows DeepTracer-2.0 to model the complete nucleotide structure from the cryo-EM density map and sequence data. After independent post-processing to complete each structure, the predicted protein and DNA/RNA models are combined, ultimately yielding a comprehensive model of the entire macromolecular complex.

*<u>Feature learning</u>***: DeepMM** uses a multi-task Densely Connected Convolutional Network (DenseNet) architecture to construct all-atom models from near-atomic resolution cryo-EM density maps [97]. Compared to CNNs, DenseNets, which connect each layer to all subsequent layers in a feed-forward fashion within each dense block, alleviate the vanishing-gradient problem, and encourage feature reuse while reducing the number of parameters. DeepMM features two embedded DenseNets. DenseNet A simultaneously predicts the probability of main-chain atoms (N, C and Cα) and Cα positions for each voxel, creating a 3D probability map. From this map, local dense points (LDPs) are then identified using mean-shift algorithm [98] and are used by a main-chain tracing algorithm, MAINMAST [24], to generate possible main-chain paths. MAINMAST connects LDPs to form a minimum spanning tree (MST), in which total distance of connected points is minimized, and iteratively refines this tree structure using a tabu search method [99] and the longest path within the refined tree is ultimately traced as the main-chain path. DenseNet B then predicts the amino acid and secondary structure types for each main-chain local dense point (LDP) on these main-chain paths. *<u>Model building</u>*: After **DeepMM** determines the Cα probability, amino acid type, and secondary structure for each main-chain point on the main-chain path, the protein’s target sequence is aligned to these main-chain paths using Smith-Waterman dynamic programming (DP) algorithm [100], which evaluates the match between the sequence and the main-chain path using scoring matrices for both amino acid and secondary structure types. The resulting Cα models are then ranked based on their alignment scores. Finally, the top-ranked Cα models are used to construct the complete all-atom protein structures with the *ctrip* program from the Jackal modeling package [101,102] and refined using the AMBER package [103].

*<u>Feature learning:</u>* **SEGEM** is an automated method that quickly and accurately builds protein backbone structures from cryo-EM density maps [104]. SEGEM employs 3D convolutional neural networks to predict both Cα locations and their amino acid types simultaneously from cryo-EM density maps. The CNN model employed for this task involves an initial image preprocessing step. The sampled sub-images are fed into three separate CNN models. Each model has a specific prediction task: one for Cα identification which generates a predicted Cα probability density map, another for amino acid type prediction, and a third for secondary structure prediction. Non-Maximum Suppression (NMS) algorithm is applied to pick out local maximum Cα probability density voxels as predicted Cα sites. These sites then have their amino acid types predicted, a vital step for accurately assigning them to the overall protein sequence. *<u>Model building</u>*: At the model construction step, **SEGEM** uses a highly parallel pipeline to efficiently match CNN predicted sites to the native protein sequence using a score matrix. Specifically, Cα local tracing connects predicted Cαs with its neighbors to form continuous traces. Next, these traces are assigned to protein sequence segments by calculating a matching score, which leverages predicted amino acid types to create an amino acid scoring matrix. For any unassigned segments, protein threading using a breadth-first search with pruning strategy is employed for faster processing ultimately yielding a complete model of Cα coordinates aligned to the protein sequence.

*<u>Feature learning:</u>* **SegmA** is a novel deep neural method for cryo-EM density map visualization and protein modeling [105]. It works by performing residue type segmentation, labeling and color-coding voxels in a cryo-EM density map based on whether they represent specific amino acids or nucleic acids. This color-coded visualization helps with both manual and automated modeling. Beyond visualization, SegmA can also predict amino acid centers of mass, score how well a protein template fits a map, and assist in *de novo* modeling of protein complexes. SegmA consists of a cascade of convolutional neural networks (CNNs) and group rotational equivariant CNNs (G-CNNs) to label voxels in a cryo-EM density map to one of the following categories: 20 amino acids, nucleotide, background or unconfident (uncertain). A G-CNN is rotation equivariant, unlike traditional CNNs which are only translation equivariant, meaning the feature maps transform accordingly when the input is rotated or translated. The Classification Net (CLF-NET), a G-CNN, performs an initial labeling of the processed volume. Its output is then passed to the Segmentation Net (SEG-NET), a U-Net CNN with contraction path (encoder) and an expansion path (decoder), which performs the final labeling of the voxels. Lastly, the Confidence Net (CNF-NET), another G-CNN, evaluates results from the SEG-NET assigning a binary confidence label to each voxel and only reporting the correct ones.

*<u>Feature learning:</u>* **ModelAngelo** is a deep-learning tool that automates atomic model building in cryo-EM density maps at resolutions better than 4.0 Å [106,107]. It generates protein models comparable to those built by human experts and creates highly accurate backbones for nucleic acid models. Additionally, ModelAngelo also identifies protein chains in cryo-EM density maps facilitating visual exploration of proteomes. ModelAngelo employs a modified feature-pyramid network [108] which is a convolutional neural network (CNN) to predict the approximate positions of protein and nucleic acid residues within the cryo-EM map. Specifically, it determines whether each voxel in the map contains an Cα atom of an amino acid, a phosphorus (P) atom of a nucleic acid residue, or neither. This process effectively initializes the graph representation, in which each residue is a node, and edges are formed between each residue and its nearest neighbors, by identifying potential residue positions. *<u>Model</u> <u>building</u>*: **ModelAngelo** employs a graph neural network (GNN) to optimize residue positions and orientations, predicts amino/nucleic acid identity, and determines side chain/base torsion angles. This GNN comprises of three modules - a cryo-EM module, a sequence module, and an invariant point attention (IPA) module - where each module refines a node associated residue feature vector by integrating new information, progressively extracting more detail from the various inputs. The cryo-EM module within the GNN extracts and integrates information from the cryo-EM density map to refine residue representations. It allows the GNN, using convolutional neural networks (CNNs), to analyze the cryo-EM density around each Cα atom and the density connecting it to neighboring nodes, thereby updating its internal representation. It uses a cross-attention mechanism to allow each residue to exchange information with its 20 closest neighbors and is driven by how connected the cryo-EM density appears between them. Unlike self-attention, which dynamically accesses all parts of the same input sequence and effectively captures long-range dependencies, cross-attention enables information flow between different data modalities. Concurrently, the module extracts a cubic section of the cryo-EM density map around current residue position which is processed by another CNN. The features extracted from this CNN are then combined with the output from the cross-attention. This combined information is used to predict amino and nucleic acid identities and outputs the updated residue feature vector. The sequence module, implemented as a Transformer module within ModelAngelo, integrates sequence information into the GNN. It performs cross-attention for each residue with the user-provided amino acid sequences, which are embedded using the pre-trained ESM-1b protein language model [109]. The output from this cross-attention is then used in two ways: a dedicated MLP (multi-layer perceptron) generates predictions for amino and nucleic acid identities, and a second MLP generates the updated residue feature vector of the sequence module. The IPA module in the GNN integrates geometric information from the nodes in the graph. It allows the model to learn the topology of neighboring residues, such as secondary structure, by assessing their spatial relationships. ModelAngelo generates the complete atomic model by post-processing residue feature vectors. These vectors serve as input to two separate MLPs that predict position and orientation for each residue, along with torsion angles for amino acid side chains and nucleic acid bases. Predictions for amino/nucleic acid identities, derived from the cryo-EM and sequence modules, are averaged to create probability distributions. These probabilities form a Hidden Markov Model (HMM) profile (**Box 1**: **Glossary section**), which is then used with *HMMER* [70] to search against input sequences. The parameters of the profile HMM are estimated from ModelAngelo predictions. Residues matching the sequence are updated, and separate chains are connected based on sequence alignment and proximity, chains shorter than four residues are removed. Finally, a complete atomic model is generated by integrating the predicted positions and orientations of each residue with their corresponding amino acid or nucleic base torsion angles, utilizing idealized geometries to ensure structural accuracy. The refined coordinates are then fed back into the GNN for three additional recycling iterations to further improve the accuracy of the model. ModelAngelo optimizes atomic positions using an L-BFGS optimizer [110]. This final relaxation step removes unnatural side-chain distances and steric clashes

*<u>Feature learning:</u>* **CryoREAD** employs a two-stage deep neural network for reconstructing nucleic acid structures from cryo-EM density maps at resolutions from 2.0 to 5.0 Å [111]. The Stage 1 network uses a cascaded, two-stage U-Net architecture, concatenating two 3D U-shape-based convolutional network (UNet) models with full-scale skip connections. The first of these U-Nets focuses on detection of sugar, phosphate, base, and protein, while the second U-Net specifically predicts individual base types (A, C, G, and T/U), leveraging information passed from the first U-Net’s encoder. The Stage 2 network then refines these initial probabilities (protein, phosphate, sugar, base, and the four base types) and generates more accurate outputs. CryoREAD applies the mean-shift algorithm [98] to cluster grid points that exceed specific probability thresholds to identify representative sugar nodes which are then connected into a graph. Edges are established between sugar nodes based on their probability and inter-node distance. *<u>Model building</u>*: **CryoREAD** uses vehicle routing problem (VRP) solver [112] (**Box 1: Glossary section**) to trace the nucleic acid backbone(s) from the predicted sugar graph. Unlike the traveling salesman problem (TSP) solver, VRP employs multiple “vehicles” to visit nodes, allowing for the identification of multiple, non-overlapping paths within the graph. This approach is well-suited for cryo-EM density maps that may contain multiple nucleic acid chains, as the VRP solver aims to maximize visited nodes while minimizing total route costs. CryoREAD assigns nucleic acid sequences to the sugar nodes within the traced paths using two sub-steps: assigning base sequence fragments to paths and assembling them. Initially, sugar backbone paths are segmented and these segments are then aligned with the nucleic acid sequence using a dynamic programming (DP) algorithm (**Box 1**: **Glossary section**) to identify the top candidate sequence fragments. Subsequently, a constraint programming (CP) solver [113] assembles these assigned sequence fragments. This solver aims to maximize the combined probability score of sugar nodes with their assigned bases while ensuring consistency across overlapping path segments and the nucleic acid sequences. At this point, the model comprises the sugar backbone and bases linked to representative sugar nodes. The final step involves incorporating nearby phosphate and base nodes that meet specific distance criteria into the sugar nodes. The output models are then refined using a two-step process: an initial refinement of predicted RNA/DNA regions using *phenix.real_space_refine* [66], followed by all-atom refinement in COOT [68].

*<u>Feature learning:</u>* **SMARTFold** is a deep learning protein structure prediction model that integrates cryo-EM density map features with sequence alignment features to accurately predict protein folds and outputs full atomic structure requiring no additional post processing steps [114]. SMARTFold first uses the protein sequence into the AlphaFold-Multimer [115] data pipeline to generate initial MSA and residue pair representations. Simultaneously, from the raw cryo-EM density map, a 3D U-Net extracts a representative point cloud, which captures backbone confidence map from the sparsely populated 3D EM density map. From this map, support points are sampled along predicted high-confidence backbone areas. The geometric features of these support points are then extracted and embedded into a point-pair representation. To maintain the relationship between these support points and the protein residues, a point-residue pair representation is introduced. Finally, these geometric features are integrated with the sequence alignment features as input for the protein folding prediction. *<u>Model building</u>*: **SMARTFold** introduced EMformer, a novel module that integrates geometric features with sequence alignment features to predict protein structure. Within EMformer, MSA, residue pair, point-residue pair, and point pair representations all exchange and update their information. Finally, an AlphaFold2-inspired [31] structure module predicts the atomic structure. Like AlphaFold2 [31], these learned representations can be recycled to further enhance model performance.

*<u>Feature learning:</u>* **EMRNA** is a deep learning-based method designed to automatically and accurately determine full-length, all-atom RNA structures directly from cryo-EM density maps (resolutions ranging from 2.0 to 6.0 Å) and RNA sequence as inputs [116]. EMRNA utilizes a Swin-Conv-UNet (SCUNet) deep learning architecture [60] to predict the probability of RNA phosphate (P), C4’, and N1/N9 atom positions, along with their corresponding nucleotide types, for every voxel in the RNA cryo-EM density map. The SCUNet architecture is built with three encoder, one transition, and three decoder Swin-Conv (SC) blocks, linked by skip connections. The “Swin” component - a shifted window transformer — excels at nonlocal modeling while the “Conv” part, a convolutional network, provides efficient local modeling. This combination gives the SC block a significant advantage over traditional convolutional neural networks, as it can effectively capture both local and long-range structural information from the cryo-EM density maps. The local maximums identified using mean-shift algorithm, within the predicted P and C4’s probability maps are then used to identify main-chain points (MCPs) which represent potential atom locations. *<u>Model building</u>*: **EMRNA** constructs the RNA backbone by threading SCUNet derived MCPs into multiple backbone traces by solving a traveling salesman problem (TSP) [117] (**Box 1**: **Glossary section**), with diverse trace types sampled from P, C4’, or combined positions. These traces are then scored by aligning them with the RNA sequence using a Smith–Waterman dynamic programming algorithm [100] and incorporating predicted secondary structure information, C4’ probabilities and nucleotide type assignments to remove incorrect paths. The most probable C4’ trace is selected via sequence alignment, after which P atoms are placed along it and N1/N9 positions are located from their respective probability maps. Once the coarse-grained backbones are established, EMRNA constructs the full-atom RNA structure. It does this by rigidly aligning A, U, G, and C nucleotide coordinates, extracted from ideal A–U and G–C pairings, onto the backbone using Kabsch superposition [118], based on their P, C4’, and N1/N9 atoms. The process then detects possible base pairings using inter-C4’ distances, followed by detecting helices and further refinement of the base-pair conformations. The output model is energy minimized using the AMBER package [103] and further refined using *phenix.real_space_refine* [66].

*<u>Feature learning:</u>* Unlike EMRNA which is specific to RNA, **EM2NA** works with cryo-EM density maps of protein-DNA/RNA or multi-chain DNA/RNA complexes at < 5.0 Å resolutions and uses deep learning to automatically build all-atom nucleic acid structures including DNA [119]. EM2NA is built on a two-stage Swin-Conv-UNet (SCUNet) network architecture [60]. The SCUNet uniquely combines Swin Transformer blocks, which excel at non-local modeling, with Convolutional Network blocks, known for their efficient local modeling. This hybrid approach allows EM2NA to leverage both local and non-local learning capabilities, outperforming traditional CNNs. In stage-1 SCUNet, EM2NA processes raw cryo-EM maps to segment and detect DNA/RNA regions, distinguishing them from protein and background. The identified nucleic acid density is then fed into the stage-2 SCUNet. Here, the network predicts nucleotide information, including the precise positions of P, C4’, and N1 or N9 atoms, as well as their corresponding nucleotide types at each voxel. Finally, the backbone atom probabilities generated by the stage-2 SCUNet are converted into 3D points by detecting the local maxima using a mean-shift algorithm.*<u>Model building</u>*: Unlike EMRNA which uses a TSP solver, **EM2NA** employs a Vehicle Routing Problem (VRP) algorithm [120] to trace SCUNet generated initial P and C4’ points into multiple backbone paths. Since VRP allows multiple paths for traveling, it is ideal for constructing multi-chain DNA/RNA structures. Determining the correct direction for each path is straightforward, leveraging the known nucleotide geometries. Once the backbone paths are established, a Smith-Waterman algorithm [100] is used to assign sequences to each backbone. This assignment is further refined by considering base pairing in double helices and helical geometry. Finally, with the built DNA/RNA backbone paths and assigned nucleotide types, the full-atom DNA/RNA structure is built by aligning template nucleotide conformations onto the P-C4’-N1/N9 backbone using the Arena algorithm [121].

*<u>Feature learning:</u>* **Cryo2Struct** automatically generates atomic protein structures from medium and high-resolution cryo-EM density maps and corresponding amino acid sequences [122]. It initiates this process by using two distinct 3D transformer-based deep learning models to classify each voxel in the cryo-EM density map. One model identifies backbone atom types (Cα, C, N, or no atom), while the other predicts the amino acid type (20 standard amino acids and the absence of an amino acid or unknown amino acid). These models are trained as sequence-to-sequence predictors, leveraging a transformer-encoder to capture long-range voxel-voxel dependencies and a skip-connected decoder (like a U-Net) for feature integration and classification. The training was conducted on the extensive Cryo2StructData dataset [65], which contains 6,652 cryo-EM maps for training and 740 for validation, followed by blind testing on two separate datasets. After prediction, a clustering strategy is applied to group spatially close Cα voxel predictions within a 2.0 Å radius, selecting the centrally located voxel to represent the final Cα atom and eliminate redundancy. *<u>Model building</u>*: To connect the predicted Cα atoms into protein chains and accurately assign their amino acid types, **Cryo2Struct** employs an innovative Hidden Markov Model (HMM) (**Box 1**: **Glossary section**) where each predicted Cα atom represents a hidden state. These hidden states are fully connected, with transition probabilities determined by the spatial distance between corresponding Cα atoms. The likelihood of each hidden state emitting a specific amino acid is based on the predicted probabilities for that Cα atom. Next, a customized Viterbi algorithm [123] aligns sequence of the target protein (or sequence for each chain for multi-chain proteins) to this HMM. This generates the most probable path of hidden states (Cα atoms), and the path for the aligned chain represents the connected Cα atoms and thus the backbone structure of the protein. For multi-chain proteins, these individual chain paths, combined with their aligned sequences, create the complete atomic backbone. The customized Viterbi algorithm ensures that each Cα position is used only once in the aligned path. This is crucial because each Cα corresponds to a single amino acid in the protein sequence. This HMM-based approach excels at assigning every amino acid of the protein to a Cα position, assuming enough predicted Cα atoms are present enabling Cryo2Struct to build very complete structural models from density maps.

*<u>Feature learning:</u>* **EModelX** is a method that constructs protein complex structure models from cryo-EM density maps and protein sequences using cross modal alignment [124]. It normalizes the cryo-EM density map and feeds it into a multi-task 3D residual U-Net. This U-Net incorporates skip-connections, which help maintain resolution despite max-pooling and address the vanishing gradient issue common in deep networks. The network predicts the distributions of Cα atoms, backbone atoms, and amino acid types. Cα candidates are identified from the predicted Cα distribution using point-cloud clustering and non-maximum suppression (NMS). *<u>Model building</u>*: **EModelX** uses predicted distributions of Cα atoms and amino acid types to sample Cα traces and generate sequence profiles from cryo-EM density maps. A Cα-sequence aligning score matrix is created, and high-confidence alignments are used to build an initial model, incorporating connectivity and sequence registration. Unmodeled gaps are then filled using a sequence-guiding Cα threading algorithm to build the Cα backbone model of protein complex, followed by full atom construction using PULCHRA tool [91] which is further refined using *phenix.real_space_refine* [66]. **EModelX(+AF),** which combines EModelX with AlphaFold2 [31] can perform template-based modeling and refine AlphaFold’s incorrectly folded structures. Cα traces are sampled from both the EModelX predicted Cα atoms and the AlphaFold2 predicted structure. The comparison of the structural similarity of these traces further improves the Cα-sequence alignment score and enhance sequence-guiding Cα threading.

*<u>Feature learning</u>***: CryFold** [125] introduces a novel approach for *de novo* model building for cryo-EM density maps, leveraging key advancements from AlphaFold2 [31] and ModelAngelo [106]. CryFold accelerates automated protein model building and produces more complete models while reducing the requirement for map resolution using two-step processes. First, a 3D convolution-based network U-Net takes the cryo-EM density map as input and output a probability map, with each voxel representing the likelihood of containing a Cα atom. The U-Net architecture features a bottleneck structure in its encoder to preserve spatial information during downsampling. Its decoder utilizes the Res2Net architecture to extract rich semantic information during upsampling. These two types of information are then combined to generate the final Cα atom probability map. The initial Cα atom coordinates are then refined using the mean-shift algorithm to obtain a precise set of Cα atom coordinates. *<u>Model building</u>*: **CryFold** then constructs the complete all-atom structure from the input density map, amino acid sequences, and predicted Cα atoms using an enhanced transformer network called Cry-Net. Cry-Net comprises of two transformer-based modules. Cryformer (encoder) transforms the density map into initial node and edge representations guided by the predicted Cα atom positions. The Structure Module (decoder) generates all-atom positions from these refined representations. Cry-Net iteratively updates these key protein structure representations through local attention mechanism leveraging spatial restraints from the density map. Cryformer processes initial node and edge representations along with ESM-2 sequence embeddings [126]. Its core components include sequence attention, which integrates protein sequence information into node features via a cross-attention mechanism using sequence embeddings from ESM-2 [126]. Node and edge attention update their respective representations through self-attention. A 3D Rotary Position Embedding (3D-RoPE) effectively encodes each node’s positional information into all attention calculations, leveraging the inherent spatial constraints from the density map. Cryformer then assigns each node to one of the 20 amino acids by using information from the original node representation (density map of the node), updated node representation (incorporating neighboring node information), and sequence data from the sequence attention layer. A separate multi-layer perceptron (MLP) generates amino acid probability vectors for all nodes. The Structure Module decodes the updated representations from Cryformer into an all-atom structure. It predicts backbone frames and torsion angles using Cryformer updated node and edge representations. The module employs a self-attention mechanism on node features, restricted to a node and its nearest neighbors using constraints from the density map, with 3D-RoPE integrating positional information into attention scores. Finally, all-atom positions for each node are generated from the predicted backbone frame, backbone and side-chain torsion angles, and amino acid type. These positions undergo a post-processing step, similar to ModelAngelo [106].

*<u>Feature learning</u>***: DeepCryoRNA**, is a novel deep learning-based method designed for automated reconstruction of RNA 3D structures from protein-free cryo-EM density maps at resolutions 6 Å or better [127]. DeepCryoRNA employs a MultiResUNet neural network [128], a variant of U-Net architecture, to predict 18 types of RNA atoms (12 backbone, 6 base) from preprocessed cryo-EM density maps. The encoder-decoder structure of MultiResUNet employs a multi-resolution design allowing it to integrate both local and global information for superior image segmentation. After prediction, atoms are clustered to remove redundancy from neighboring voxel predictions. *<u>Model</u> <u>building</u>*: **DeepCryoRNA** constructs nucleotides based on the clustered atoms predicted from MultiResUNet by factoring in atom classes and pairwise atomic distances. It identifies nucleotide types by analyzing base atom classes and their quantities. The process then links neighboring nucleotides into short chains, and these short chains are further connected to form complete long chains. Multiple complete chains can be derived, representing various connection pathways. The model uses a modified Gotoh algorithm [129] for global sequence alignments to match these complete long chains with native RNA sequences. DeepCryoRNA then selects the top 10 alignment results to assign native chain information to the corresponding complete long chains, yielding 10 all-atom RNA structures. These structures undergo post-processing, including energy minimization ultimately generating refined RNA 3D structures. The all-atom RNA structures are then refined using QRNAS software [130], to fix broken bonds and resolve steric clashes between atoms.

*<u>Feature learning</u>***: E3-CryoFold** is an efficient, end-to-end deep learning method that takes a cryo-EM density map and corresponding protein sequence as input and provides a one-shot inference to output the complete atomic structure [131]. E3-CryoFold concurrently uses 3D and sequence transformers to extract features from density maps and protein sequences, respectively. While self-attention captures long-range dependencies within each modality, cross-attention modules integrate information between them. In E3-CryoFold, cross-attention is used to integrate spatially contextualized information from density maps into the sequence representation facilitating the integration of information from both modalities. To ensure this integration, E3-CryoFold embeds both modalities into a shared hidden space using 3D and sequence encoders. *<u>Model building</u>*: **E3-CryoFold** constructs the final 3D atomic models using an SE(3)-equivariant (**Box 1**: **Glossary section**) graph neural network, SE(3) GNN, which is conditioned on the extracted combined spatial-sequential features. SE(3)-equivariant GNN is a graph neural network for 3D data that ensures predictions transform consistently when the input is rotated or translated in 3D space. E3-CryoFold reconstructs the protein backbone by first initializing random coordinates and building a k-nearest neighbors (kNN) graph to define local spatial relationships between residues. Node embeddings are derived from integrated spatial and sequence features which are generated by combining spatial information from the cryo-EM density map with sequence information. These embeddings capture both local and global protein features, allowing the model to utilize the inherent relationship between a protein’s sequence and its 3D structure. Each residue’s local frame (orientation and position) is iteratively updated by an SE(3) GNN, which aggregates relative rotation and translation information from neighbors. This process ensures the backbone reconstruction respects geometric relationships and spatial transformations, ultimately allowing the recovery of 3D coordinates for all backbone atoms.

**4.3.2 Hybrid model building:** In this approach, deep learning tools generally integrate voxel-wise backbone (and sometimes sidechain) atom, secondary structure and residue type probabilities learned from the cryo-EM density map with template structures or fragments (such as those predicted by AlphaFold2 [31] or derived from the PDB [13]) to accomplish model building.

*<u>Feature learning</u>***: SEGEM++**, an enhanced version of the SEGEM method integrates AlphaFold2 (AF2) [31] protein structure prediction algorithm with data from cryo-EM density maps [104]. This allows SEGEM++ to not only identify accurately folded regions within AF2 structures by utilizing SEGEM predicted Cα probability densities from cryo-EM density maps, but also to correct incorrectly folded areas through protein threading on the cryo-EM map itself. *<u>Model building</u>*: **SEGEM++** calculates a confidence score for each Cα in the AF2 structure, based on its alignment with the SEGEM predicted Cα probability density map. This allows SEGEM++ to identify high-confidence, correctly folded AF2 structure fragments. These reliable fragments then serve as an improved base model, guiding subsequent protein threading on the predicted Cα probability map to build a more accurate final protein structure. In essence, SEGEM++ leverages strong AF2 predictions to refine its cryo-EM model, while simultaneously using cryo-EM data to validate and correct any inaccuracies in the AF2 predictions.

*<u>Feature learning</u>***: CR-I-TASSER** (cryo-EM iterative threading assembly refinement) is a hybrid method that combines deep neural-network learning with I-TASSER assembly simulations to automate cryo-EM structure determination [132]. It uses multithreading algorithms to identify templates from the Protein Data Bank (PDB) [13], aiding structural assembly. CR-I-TASSER employs a 3D convolutional neural network (CNN) with a residual architecture to create sequence-order-independent Cα atom trace models from cryo-EM density maps to improve threading template quality. *<u>Model</u> <u>building</u>*: **CR-I-TASSER** employs deep learning-based template refinement and regeneration, and density map-guided structural reassembly simulations. Using local meta-threading server (LOMETS) [133], CR-I-TASSER derives threading templates from the PDB. The 3D CNN predicted Cα conformation then refines threading templates through multiple heuristic iterative algorithms that align query and template sequences with the Cα conformation for template reselection and Cα trace regeneration. Finally, guided by cryo-EM density map correlations and deep-learning derived template restraints, the iterative threading assembly refinement method (I-TASSER) method [134] assembles full atomic structures which are then refined using fragment-guided molecular dynamics [135].

*<u>Model prediction</u>***: DEMO-EM** (domain enhanced modeling using cryo-electron microscopy) is an automated method designed to assemble accurate full-length structural models of multi-domain proteins from cryo-EM density maps by integrating single-domain modeling and deep residual network learning techniques with progressive domain assembly and refinement procedure [136]. DEMO-EM uses D-I-TASSER [137], which incorporates deep learning-based spatial restraints (including inter-residue contact and hydrogen-bonding potentials) into its iterative threading assembly simulations to generate an initial structural model for each domain. Meanwhile, inter-domain distances are predicted by DomainDist, a deep convolutional neural-network architecture with ResNet basic blocks. DomainDist guides the assembly of domain orientations by providing inter-domain distance maps. Each individual domain model generated by D-I-TASSER is independently fitted to the density map using a quasi-Newton optimization algorithm, Limited-memory Broyden–Fletcher–Goldfarb–Shanno (L-BFGS) [110]. Since L-BFGS is a local optimization method, simulations are initiated from multiple starting positions to identify the best location and orientation of the domain with the highest correlation with the density map. The initial full-length models are then optimized through a two-step assembly and refinement process. Model-density correlations primarily guide the domain assembly and refinement simulations. Following a rigid-body Replica Exchange Monte Carlo (REMC) simulation, the top scoring model from this stage is then refined further by flexible assembly, which incorporates atom, segment, and domain-level refinements using REMC simulation guided by the density correlation and inter-domain distance profiles. Finally, the lowest-energy model undergoes side-chain repacking with FASPR [138] to create the final model which is refined by fragment-guided molecule dynamics (FG-MD) simulations [135]. DEMO-EM can also assemble domain structures generated by any method other than D-I-TASSER. **DEMO-EM2**, an improved version of DEMO-EM, is an automated method for constructing protein complex models from cryo-EM density maps [139]. Unlike DEMO-EM, which is designed for multi-domain proteins, DEMO-EM2 focuses specifically on assembling protein complexes through an iterative assembly procedure. Instead of using D-I-TASSER, DEMO-EM2 employs AlphaFold2 [31], due to its outstanding performance in protein structure prediction, to derive model of each individual chain. DEMO-EM2 incorporates several advancements over DEMO-EM including preprocessing the density map to reduce interference from noise during chain or domain fitting and using a differential evolution (DE) algorithm [140] in addition to the quasi-Newton optimization, preventing it from getting trapped in local optima. Further, it also masks out density map regions that have already been matched with chain models, ensuring different chains do not align the same areas.

*<u>Feature learning</u>***: DeepTracer-ID** is a server-based, *de novo* protein identification method that uses high-resolution cryo-EM density maps (better than 4.2 Å resolution) to identify candidate proteins within a user-specified organism without requiring additional information [141]. It achieves this by using DeepTracer [94], a deep learning method, to automatically generate a protein backbone model from the input cryo-EM density map. *<u>Model building</u>*: DeepTracer generated protein backbone model is used by **DeepTracer-ID** to search against the library of AlphaFold2 [31] predictions for all proteins in the given organism using three different alignment algorithms. PyMOL-align [39] considers both sequence and structural similarities and is the default option. PyMOL-cealign [142] is ideal for proteins with low or no sequence similarity, or when side-chain densities in the cryo-EM map are not well-resolved. FATCAT [143] specializes in flexible protein structure comparison, simultaneously optimizing alignment and minimizing rigid-body movements. It can be particularly useful for mitigating errors in AlphaFold2 predictions, and for smaller proteins or those where local environment dictates their 3D structure.

*<u>Feature learning</u>***: EMBuild** is an automated, deep learning-based method designed to construct multi-chain protein complex models directly from intermediate-resolution cryo-EM density maps [144]. EMBuild employs a nested U-Net (UNet++) architecture with dense skip connections to predict a precise main-chain probability map from the input cryo-EM density map, and AlphaFold2 [31] to predict 3D structures of the input protein sequences. The main-chain probability map assigns a probability to each grid point, indicating the likelihood of a main-chain atom being present in that vicinity. Instead of directly fitting protein chains to the raw cryo-EM density, EMBuild uses the accurate main-chain probability map with more precise location information for main-chain atoms, which significantly improves the precision of subsequent chain fitting. *<u>Model building</u>***: EMBuild** aligns each AlphaFold2 [31] predicted structures of individual protein chains to the main-chain probability map using a Fast Fourier Transform (FFT)-based global alignment [145]. To account for potential deviations between the input protein chain model and ground truth structure, a semi-flexible domain refinement strategy is then employed: each chain is first rigidly fitted, and then its individual structural domains are locally refined. For each fitted protein chain, EMBuild calculates a main-chain match score to quantify its fit to the probability map. With all individual chain fitting results, the final protein complex structure is assembled by identifying the optimal combination of fitted chains. This is achieved through an iterative Bron–Kerbosch maximum clique algorithm, which selects combinations with the highest total main-chain match score while preventing severe atomic clashes between chains. Unassembled chains are then iteratively integrated into the complex through further fitting and the complex is refined using *phenix.real_space_refine* [66]. Structural category annotations from EMInfo [86] have been shown to improve the modeling accuracy of EMBuild. EMBuild treats all density voxels equally during fitting, which can lead to inaccuracies in fitting at protein regions containing a mixture of α-helices, β-sheets, and coils. By incorporating secondary structure details from EMInfo, **EMBuild+EMInfo** can accurately fit protein fragments by matching them to density voxels of the corresponding secondary structure type.

*<u>Feature learning:</u>* **FFF** (”Fragment-guided Flexible Fitting”) uses a deep-learning-based multi-level recognition network to capture diverse structural features from cryo-EM density maps [146]. Inspired by RetinaNet (ResNet-based), but adapted with 3D convolutions, this network predicts not only a voxel-wise backbone probability map but also unifies four distinct coarse-grained tasks: Cα atom detection, Cα location prediction, pseudo-peptide vector (PPV) estimation, and amino acid (AA) classification. The network’s backbone (BB) component identifies whether each voxel is part of the protein backbone. The Cα detection module then estimates the likelihood of a grid cell containing a Cα atom. For cells likely to have a Cα atom, the amino acid classification module predicts the specific type of amino acid. Finally, the pseudo-peptide vector (PPV) estimation module determines vectors connecting a Cα atom to its consequent Cα atoms. *<u>Model building</u>*: **FFF** uses the extracted pseudo-peptide vectors (PPVs) to generate and recognize protein structural fragments. This involves connecting the Cα atoms of residues into fragments, a process guided by selecting neighboring atoms based on the estimated PPVs and the known protein sequence. Once the protein fragments and backbone maps are identified, targeted molecular dynamics (TMD) [147] is used to refine and update an initial structure, aligning it with the recognized fragments. Next, molecular dynamics flexible fitting (MDFF) [148] updates the entire backbone conformation of initial structure to match the predicted backbone map yielding the complete protein structure of the target protein. During this MDFF step, positional restraints are added to the atoms initially selected in the TMD phase, preventing significant deviations.

*<u>Feature learning</u>***: CrAI** is an automatic deep learning method that detects and aligns antibodies (Fabs and VHHs) within cryo-EM density maps at resolutions up to 10.0 Å [149]. The core of CrAI is a customized 3D U-Net architecture, trained on a curated dataset. It uses a unique representation of antibody structures to facilitate the learning process, framing the task as a special instance of 3D object detection. To prevent redundant predictions of overlapping objects, CrAI employs a Non-Maximal Suppression (NMS) algorithm, a crucial post-processing technique to refine the output of object detection models. *<u>Model prediction</u>*: After detection, **CrAI** fits pre-classified Fab or VHH templates to the predicted locations and poses, providing a structural model of antibodies (Fabs and VHHs) within cryo-EM density maps.

*<u>Feature learning</u>***: DeepMainmast** is a method for protein structure modeling from cryo-EM density maps at resolutions between 2.5 and 5.0 Å [150]. It achieves this by combining protein main-chain tracing using deep learning with structure modeling from AlphaFold2 [31]. DeepMainmast utilizes Emap2sf (Emap to structural features), a deep-learning method having a U-shaped network (UNet) architecture with skip connections [151]. The network, consisting of three encoder blocks and two decoder blocks built upon a three-dimensional convolutional layer (Conv3d), outputs probability values for 20 amino acid types and backbone atom (N, Cα, C) at each grid point in the density map which is required for subsequent Cα-tracing. Local dense points (LDPs) are created by clustering grid points with a high probability for Cα using the mean shift algorithm [98]. *<u>Model building</u>*: **DeepMainmast** reconstructs protein structures from cryo-EM density maps through a multi-stage process. It starts by connecting LDPs, identified from high Cα probability regions, into Cα paths using a VRP (Vehicle Routing Problem) solver which efficiently finds optimal routes by minimizing costs based on distance and main-chain atom probabilities. Once Cα paths are established, they are aligned with the target protein sequence using the Smith-Waterman algorithm [100], with matching scores determined by DAQ(AA) scores [152] calculated from Emap2sf output. This generates numerous Cα fragments, with the entire process repeated across various parameter combinations (Cα probability cutoff, number of VRP vehicles, and cost function parameters). A key innovation in DeepMainmast is the integration of AlphaFold2 (AF2) models, specifically the top-ranked one based on pLDDT scores. AF2 models contribute by providing additional Cα fragments to fill gaps in low-density regions and also serve as global structure for fitting to the density map. Next, Cα protein models are assembled from these combined fragment libraries using a constraint programming (CP) solver [153]. This solver optimizes fragment combinations to maximize the total DAQ score while preventing steric clashes, ensuring consistent amino acid positioning, and maintaining consistent Cα-Cα distances. In parallel, AF2 models are also directly superimposed onto the density map using structure fitting program, VESPER [154]. Finally, these refined Cα models are converted into full-atom structures using PULCHRA [91], with any missing regions subsequently filled and refined by RosettaCM [155].

*<u>Model prediction</u>*: **DeepTracer-Refine** [156] improves protein structure prediction by combining DeepTracer [94] (a map-to-model method) with AlphaFold2 [31] (a sequence-to-model method). It splits AlphaFold structures into compact domains, identifying optimal separation points based on AlphaFold’s predicted Local Distance Difference Test (pLDDT) metric, which estimates the confidence level for each residue in its prediction. Each of these smaller domains then undergoes rigid body alignment using a selection of algorithms, including PyMOL cealign [142], PyMOL align [39], and Chimera MatchMaker [157], with the best fit chosen for maximum residue coverage. This iterative alignment process continuously updates AlphaFold’s residue locations as each domain is aligned to DeepTracer’s prediction, ultimately providing a more refined and accurate protein structure. *<u>Model</u> <u>prediction</u>*: **DeepTracer-LowResEnhance** [158] is a computational method that enhances low-resolution cryo-EM density maps by integrating structural predictions from AlphaFold2 [31] with a deep learning-based map refinement strategy. The input sequence is first processed by AlphaFold2 to generate an initial 3D structure, which is then used to generate a simulated map using ChimeraX [82]. Both the simulated map and the original cryo-EM map are then fed into the CryoFEM [159] module which averages these maps, splits them into chunks, and uses a UNet-based deep neural network to reconstruct a refined map. This integration leverages AlphaFold’s accurate sequence-based structural predictions with cryo-EM data, significantly enhancing model quality in low-resolution cases. Finally, DeepTracer [94] generates a high-accuracy 3D protein structure model from the refined cryo-EM map.

*<u>Feature learning</u>***: CryoJAM** is a deep learning-based tool designed to automate and enhance the challenging process of fitting large protein complexes into medium-resolution cryo-EM density maps, thereby accelerating their structural modeling [160]. The 3D convolutional neural network (CNN) of CryoJAM leverages a U-Net architecture and a novel composite loss function that incorporates both Fourier-shell correlation (FSC) and Root Mean Squared Error (RMSE). FSC serves as a proxy for the quality of fit in Fourier space, while RMSE directly optimizes atomic accuracy in real space. The UNet-based architecture of CryoJAM handles both 3D volumetric cryo-EM densities and homolog structures and its outputs represent the adjusted homolog coordinates. *<u>Model building</u>*: **CryoJAM** generates a volume representing predicted Cα atom locations within the cryo-EM density highlighting backbone density. Since this output is continuous volume, CryoJAM employs a post-processing workflow to derive discrete all-atom coordinates. This involves using a KD-tree [161] to select the top Cα voxel activations, binarizing the volume, and outputting 3D coordinates. A greedy matching algorithm aligns these selected Cα atoms to their closest counterparts in the input structure for adjustment. Finally, these Cα traces are processed by PULCHRA [91] which can construct physically realistic all-atom structures from only Cα coordinates.

*<u>Feature learning</u>***: DiffModeler** is a fully automated method designed to model large protein complex structures, effectively fitting them into cryo-EM density maps with resolutions up to approximately 15.0 Å [162]. It employs a diffusion model to trace protein backbones by capturing local density patterns representing protein backbones in low resolution cryo-EM density maps. It then integrates this diffusion model enhanced map with AlphaFold2 [31] predicted structures for accurate structure fitting, thereby enhancing the extraction of structural information from intermediate-resolution cryo-EM density maps. A diffusion model is a generative model within a probabilistic framework, trained to generate data samples that closely resemble the underlying data distribution. The conditional diffusion model of DiffModeler starts with random Gaussian noise and a cryo-EM density map as inputs. The model employs encoder-decoder network architecture (based on U-Net). The encoder first processes a cryo-EM density box, computing and embedding its hidden features. The decoder then begins with random Gaussian noise and iteratively refines its density estimates, moving closer to the ground-truth traced backbone. The entire encoder-decoder network is optimized by comparing the predicted and ground-truth traced backbones. Once trained, the conditional diffusion model generates a refined, traced backbone based on the input cryo-EM density. *<u>Model building</u>*: **DiffModeler** fits AlphaFold2 [31] predicted single-chain protein structures into the diffusion model enhanced map using VESPER algorithm [154]. VESPER aligns each subunit with the diffused map and generates the top 100 candidate poses. A subsequent assembly phase then uses a greedy algorithm to combine suitable poses from these subunits, thereby constructing the complete protein complex structure. Additionally, DiffModeler splits the multi-domain AlphaFold2 structures into individual domains using SWORD2 [163] and uses these domains in the fitting process to mitigate inaccuracies, particularly in AlphaFold2 models where domain orientations are incorrect despite accurate individual domains. **DMcloud** is local structure fitting tool for medium to low resolution cryo-EM density maps [164]. It fits structures by converting molecular models and cryo-EM density maps into point clouds for precise local alignments and iterative refinements to address erroneous AlphaFold2 models that have accurate local structural details, but their global conformation is inaccurate.

*<u>Feature learning</u>***: Cryo2struct2** [165] is a deep learning model that combines sequence-based features from a Protein Language Model (ESM) [126] with cryo-EM density maps to derive templates for AlphaFold3 [166] structure predictions. This integration allows Cryo2Struct2 to generate more accurate atomic models, especially for large proteins with flexible or complex conformations and those with regions of low-resolution and missing density. Unlike Cryo2Struct [122] which uses two separate 3D transformer models to predict atom types and amino acid types respectively, its successor Cryo2Struct2 uses a unified deep learning model with a shared transformer encoder to extract features from cryo-EM density maps. This encoder feeds into two specialized decoders for atom-type and amino acid-type predictions. The architecture of the model is based on 3D SegFormer [167] and is designed to integrate ESM protein language model embeddings with map features. This is done by transforming the ESM embeddings via a multi-layer perceptron (MLP) and adding them to the multi-scale feature representations from the map, ensuring sequence-level information is incorporated. The transformer encoder, utilizing an efficient self-attention mechanism, captures hierarchical features from the density map, while each decoder predicts its specific labels (atom types or amino acid types) for every voxel. The atom-type decoder directly predicts labels (Cα, N, C, or no atom), while the amino acid-type decoder, benefiting from the atom-type features as an auxiliary input, predicts 21 different amino acid classes (20 standard amino acids plus the absence of an amino acid or unknown amino acid). Cryo2Struct2 is also trained on Cryo2StructData dataset [65] and uses two clustering thresholds: 2 Å and 3 Å for clustering predicted Cα voxels. *<u>Model building</u>*: **Cryo2Struct2** uses a two-step process to generate accurate protein models. First, predicted atom and amino acid type probabilities are used to build a Hidden Markov Model (HMM) (**Box 1**: **Glossary section**). This HMM, processed by a modified Viterbi algorithm [123] (similar to Cryo2Struct [122]), aligns the protein amino acid sequence to the predicted and clustered Cα voxel coordinates to construct an initial 3D atomic protein backbone. These initial structures then serve as templates for advanced structure prediction capabilities of AlphaFold3 [166] and guide AlphaFold3 to generate structures that are consistent with the cryo-EM density. To obtain structurally meaningful templates, the query protein sequence is aligned to the template sequences derived from Cryo2Struct2 generated structural predictions. The templates allow AlphaFold3 to refine the structure, incorporate prior structural information, and ultimately improve the accuracy of the final atomic model while maintaining consistency with experimental cryo-EM density data.

*<u>Feature learning</u>***: DEMO-EMfit** is a method for fitting atomic structures of protein and protein-nucleic acid complexes into cryo-EM and cryo-ET maps [168]. It integrates deep learning-based backbone map extraction from cryo-EM density map with a global-local structural pose search and optimization. Since DEMO-EMfit utilizes the correlation between the cryo-EM density map and the backbone atoms of the structure during fitting procedure, it first extracts key structural features from input density maps. For this, it leverages DiffModeler [162], a deep learning method based on diffusion model, to generate a backbone density map from input cryo-EM density map that exclusively contains backbone atom information. *<u>Model building</u>*: **DEMO-EMfit** employs Fast Fourier Transform (FFT) and Limited-memory Broyden–Fletcher–Goldfarb–Shanno (L-BFGS) [110] algorithms for global and local searches to determine the optimal structure pose. Initially, an FFT-based global search generates raw poses by exhaustively exploring possible structure orientations in Fourier space, evaluating them using the density correlation coefficient between the structure and the map. The top-scoring poses from this global search then undergoes a local search using the L-BFGS algorithm for refinement. Finally, domain-level optimization is applied to further refine the fitted model, addressing potential domain-level biases.

*<u>Feature learning</u>***: DEMO-EMol** is an improved server for accurately assembling protein-nucleic acid complex structures from cryo-EM density maps [169]. It integrates deep learning-based map segmentation of protein and nucleic acid regions with an iterative structure fitting and assembly process guided by map constraints. DEMO-EMol begins by segmenting protein and nucleic acid regions from the input density map using the U-Net++ architecture from EMNUSS [80] with the training dataset obtained from the first stage dataset of EM2NA [119], another deep learning tool specific for modeling nucleic acids in cryo-EM maps. These separate protein and nucleic acid density maps are then used to iteratively fit and assemble their respective chain models. *<u>Model building</u>*: **DEMO-EMol** independently fits protein and nucleic acid chain models into their respective segmented maps by sequentially optimizing their poses using the L-BFGS algorithm [110]. Since L-BFGS is a local optimization method, multiple initial poses are explored for each chain. To enhance accuracy and reduce the search space, a map masking strategy is employed, masking map regions already matched. The L-BFGS optimization uses a composite scoring function that integrates global and local model-to-map correlation coefficients [168] along with Fourier Shell Correlation (FSC). Once all chain models are fitted, DEMO-EMol constructs the final complex model by identifying the optimal combination of all chain poses using a differential evolution algorithm [170], followed by a domain-level flexible refinement where the positions and orientations of all protein domains are simultaneously optimized.

*<u>Feature learning</u>***: CryoDomain** is a deep neural network that identifies protein domains from low-resolution cryo-EM density maps by leveraging a dual-tower network architecture: the DensityTower and the AtomTower [171]. Each tower undergoes self-supervised pre-training on its respective modality - raw cryo-EM density maps for the DensityTower and atomic structures for the AtomTower - to extract modality-specific features. CryoDomain then simultaneously learns embeddings from both protein domain density maps and atomic structures within a shared, low-dimensional space. The network integrates these two modalities into a unified representation through an alignment process. The DensityTower (U-Net-like architecture) network comprises of a Residual U-Net, a Swin-Conv U-Net, Conv module, and a Compress Module. The Residual U-Net and a Swin-Conv U-Net progressively learn both local and non-local spatial features from cryo-EM density maps. The Conv module reconstructs the map, and the Compress Module projects it into a density map embedding. In the AtomTower network, an AtomEncoder, Structure Module, and Fusion Module collectively extract an atomic structure embedding from the input atomic structure. During training, density map embedding is aligned with its corresponding atomic structure embedding. *<u>Model building</u>*: **CryoDomain** effectively transfers knowledge from rich atomic structure datasets to sparse density map datasets by integrating these two modalities through cross-modal alignment. This alignment maximizes similarity between embeddings of the same domain types while minimizing similarity between different ones, creating a unified low-dimensional representation. After alignment, CryoDomain constructs a Density-atom embedding Database (DateDB) of atomic structure embeddings, enabling protein domain identification from density maps through embedding retrieval.

*<u>Feature learning</u>***: MICA** [172] is a deep learning method that combines cryo-EM density maps with AlphaFold3 [166] predicted structures to create more accurate protein models in cryo-EM density maps of resolution 1.0 – 4.0 Å. Unlike other methods that use predicted structures at the end in the post-processing step, MICA integrates AlphaFold3 predicted structures with cryo-EM density maps at both input and output levels. This allows MICA to integrate the strengths of both data types, the experimental accuracy of cryo-EM maps and the completeness of AlphaFold3 predictions and compensate for low-resolution areas in maps or inaccuracies in AlphaFold3 predictions of large protein complexes. At the input stage, MICA combines cryo-EM density maps and AlphaFold3 predicted structures through representation learning before passing them to a deep learning network to build protein structures. Its deep learning architecture processes the fused representation of cryo-EM density maps and AlphaFold3 structures using a multi-task encoder-decoder system with a feature pyramid network (FPN) [108] to predict the locations of backbone atoms, Cα atoms, and amino acid types. The predicted Cα candidates are refined using DBSCAN clustering strategy [173] and non-maximum suppression algorithm to output Cα atoms which along with their amino acid type predictions are used as input for the backbone tracing to build an initial protein backbone model. *<u>Model building</u>*: **MICA** uses the backbone tracing procedure of EModelX(+AF) [124] to build an initial backbone model from the predicted backbone atoms, Cα atoms and amino acid types. It starts by identifying high-confidence Cα atoms, linking them as protein chains and assigning their amino acid types followed by filling in any gaps in the initial backbone model by using information from AlphaFold3 [166] structures. This Cα backbone model is then converted into a full-atom model using PULCHRA [91] and refined using *phenix.real_space_refine* [66].

## 5. Assessment and Validation

Assessing the accuracy of the predicted models is the critical final step in the model building pipeline [174]. The assessment methods can be broadly classified into three categories: predicted-target structure assessment, map-model assessment and model quality assessment. Evaluation metrics for assessing the accuracy of **predicted protein models against target structures** fundamentally rely on the alignment of Cα atoms, as these atoms represent the backbone of individual residues and thus their spatial arrangement. Evaluation metrics commonly used to quantify the percentage of correctly paired Cα atoms are recall, sequence recall, precision, F1-score, Cα matching score, Cα sequence match score, Cα quality score and TM-score. **Recall** measures the percentage of residues where the predicted Cα atom is within 3.0 Å of the deposited model [107]. **Sequence recall** measures the percentage of residues where the predicted Cα atom is within 3 Å of the deposited model and the predicted amino acid type is correct [107]. **Precision** is the percentage of predicted Cα atoms that fall within 3 Å of a Cα atom in the deposited map[107]. The **F1 score** is the harmonic mean of precision and recall for Cα atoms[107]. This metric offers a balanced assessment, considering both the specificity and sensitivity of predicted Cα atom positions. The **Cα match score** is the percentage of Cα atoms (residues) in a predicted model that are within 3.0 Å of their corresponding residues in the true structure [20]. The **sequence match score** indicates the percentage of aligned residues that possess the identical amino acid type as their corresponding counterparts in the true structure [20]. The **Cα quality score** is calculated by multiplying the Cα match score by the ratio of the total predicted residues to the total residues in the experimental structure [122]. A standard **TM-score** [175] quantifies the structural similarity between a predicted model and its corresponding known structure. The **normalized TM-score** [122] is the TM-score of the atomic models, adjusted by the length of the known structure. For the deep learning-based model building methods discussed, their performance metrics are available in their respective publications. However, note that the test datasets used for evaluation often differ between these methods. The existing evaluation metrics have certain limitations because they often ignore chain-level correspondence and, in the case of TM-score, do not account for residue identity. Improved evaluation metrics such as **TMRR-score** [37] have been introduced that combines TM-score with residue-recall, to measure both structural and residue type similarities. A recent study [37] comprehensively benchmarked state-of-the-art model-building approaches using the TMRR-score and 50 cryo-EM density maps across various resolutions. This assessment evaluated how well predicted models aligned from atomic to intermediate resolutions, their runtime efficiency, and the benefits of integrating structure prediction techniques.

**Model validation methods** use validation criteria such Ramachandran plot outliers, all-atom clash scores, deviations in bonding geometry, and rotamer preferences to assess the quality and accuracy of the macromolecular structures. MolProbity [176], frequently used and incorporated into the PDB validation reports, is a comprehensive web server for validating the quality of 3D structures, including proteins, nucleic acids, and their complexes. It provides detailed all-atom contact analysis to identify steric clashes and offers updated diagnostics for dihedral angles, hydrogen bonds, and van der Waals contacts at molecular interfaces. MolProbity combines multiple geometric parameters into a single, overall MolProbity score, where lower values indicate higher model quality. Specifically, this score is a log-weighted combination of the clashscore, the percentage of Ramachandran plot outliers, and the percentage of side chain rotamer outliers. MolProbity is unique in offering all-atom contact analysis and its use of highly accurate, up-to-date Ramachandran and rotamer distributions. Another metric, CaBLAM [177] (Calpha-Based Low-resolution Annotation Method) assesses backbone geometry of proteins by detecting Cα -geometry outliers to identify areas of probable secondary structure. CaBLAM is designed for low-resolution structures, where complex errors or ambiguities can render highly sensitive conformational analyses, like Ramachandran analysis, difficult or impossible to interpret. Among the deep-learning based model validation methods, the DAQ [152] (Deep-learning-based Amino-acid-wise Quality) score has been computed for all the PDB entries from cryo-EM maps in the resolution range 2.5 Å - 5.0 Å. DAQ score assesses the local model quality at the residue level. A key advantage of DAQ is its ability to identify regions with incorrect amino acid assignments (e.g., sequence shifts), even when the backbone is accurately modeled. The predicted local distance difference test (pLDDT) score is a valuable tool for assessing AlphaFold [31] predicted models, especially since AlphaFold predicted models are now commonly used in cryo-EM model building. pLDDT is a per-residue measure of local confidence for predicted protein structures, scaled from 0 to 100. Higher scores indicate greater confidence and typically a more accurate prediction.

**Map-model scores** quantitatively assess how well a structure model fits the experimental cryo-EM density map. Several software packages such as TEMPy [178], CCP-EM [179] and Phenix [180] offer tools to calculate cross-correlation scores to evaluate how well a structural model fits its cryo-EM map. Recent advancements have introduced local map-model fitting scores like Strudel score [181], EMRinger [182], Q-scores [183], DeepQs [184], FSC-Q [185], and MEDIC [15], which provide more granular insights into model quality. 3D-Strudel [181] is a model-dependent tool for validating map features in cryo-EM structures ranging from 2 to 4 Å resolution. It calculates a Strudel score, which is the linear correlation coefficient between the experimental map values around a target residue and the values from a rotamer-specific map-motif obtained from the 3D-Strudel motif library. EMRinger [182] evaluates how well side-chain conformation of a residue in a model aligns with the cryo-EM map compared to other common rotamers. Q-score [183], now included in the PDB validation reports, quantitatively assesses the resolvability of individual atoms by comparing their density profiles to an ideal Gaussian reference profile of the atom. DeepQs [184] uses a 3D Vision Transformer (ViT) to estimate local Q-scores in cryo-EM maps up to 5.0 Å resolution. FSC-Q [185] quantifies local resolution differences between a cryo-EM map and an atomic model through a localized Fourier Shell Correlation (FSC) analysis. It compares the local resolutions derived from small sub-maps of both the experimental map and a model-derived map. An FSC-Q value near zero indicates strong support for atoms by the map. MEDIC (Model Error Detection in Cryo-EM) [15] is a robust statistical model that identifies local backbone errors in cryo-EM protein structures. It combines local fit-to-density with deep-learning-derived structural information, including energy metrics and a predicted error score from a machine learning model trained to distinguish native from decoy structures.

## 6 Availability and Applications

Most of the tools we have discussed are open source, with their source code publicly accessible on repositories like GitHub and Zenodo [186] (links provided in **Table 5**). These platforms provide detailed information on everything, from software and hardware dependencies to specific local installation instructions for these tools. This ensures users can successfully set up and run these programs on their own computing systems, provided they meet the necessary computing software and hardware requirements. Beyond local installations, many of these tools are also readily available on web-based or cloud-based computational platforms. This includes services like Cosmic Cryo-EM [187], Code Ocean [188], and Google Colab, as well as dedicated servers (links provided in **Table 5**). These platforms are particularly beneficial for users who may not possess the robust computational infrastructure required for local execution, such as high-performance GPUs. On these cloud-based environments, users can simply upload their necessary inputs, such as a density map or a sequence, and receive the processed output, often a built model, without needing to manage the underlying computational resources themselves. Deep learning–based automated model-building methods have become an integral part of the structural biologist’s toolkit, enabling applications that span from contaminant identification in heterogeneous cryo-EM datasets to the structure determination of physiologically important macromolecular complexes, as well as method development across diverse areas of structural biology. Cryo-EM studies where these tools have had notable contributions are discussed in the **Supporting Information.**

## 7 Limitations and Future directions

In general, the performance of deep learning-based automated model-building methods declines as the resolution of cryo-EM density maps decreases, with accurate models typically obtained only within the resolution ranges represented in their training datasets. Cryo-EM density maps often exhibit heterogeneous resolution distribution within a single reconstruction. Thus, even when the global resolution of the map falls within the preferred resolution range of a specific method, regions of lower local resolutions in the map may lie outside this range, resulting in inaccurate or incomplete structural models in those areas. Moreover, modeling efforts focus on interpretable, high-resolution regions, while lower-resolution and uninterpretable areas remain poorly characterized and lack structural labels. As the map resolution declines, backbone tracing and sequence registration become challenging for *de novo* approaches and may require assistance from external structure prediction methods. For hybrid methods, the accuracy of the output models may be affected by the quality of the structural templates integrated with voxel-wise structural predictions during model generation. Cryo-EM density maps contain information about the conformational heterogeneity and dynamic regions of biomolecules. However, most deep learning–based automated model-building methods are limited by their ability to handle conformational heterogeneity and produce static models from inherently dynamic cryo-EM density map inputs. CryoBoltz [189] uses global–local constraints derived from input density maps to guide the sampling process of Boltz-1 [190], an open-source diffusion-based protein structure prediction model. This enables CryoBoltz to generate structural ensembles that capture the underlying conformational heterogeneity of the maps. Since simulated maps lack the complexity present in experimental cryo-EM maps, such as heterogeneous resolution, conformational heterogeneity, and complex noise, deep learning-based methods trained on these simulated maps may not perform well on experimental cryo-EM maps even when the input map resolution is within the resolution range of such methods. The strengths and limitations of different deep learning architectures (**Section 3**) directly influence the performance of automated model building tools that utilize them. This means that if a particular deep learning architecture has inherent limitations for a given task, those same limitations will be evident in the tools built upon it. Conversely, neural network architecture with specific advantages for certain prediction tasks will lead to more effective tools.

***findMySequence*** [69], a tool to identify protein sequences in crystallography and cryo-EM maps, depends on the accuracy of the traced backbone, fragmented or mistraced models often found in low-resolution maps may affect sequence identification. While the approach scales to multimeric assemblies, it requires manual selection of intermediate fragments in an iterative modeling process.

Nevertheless, the application is very fast for majority of cases. The performance of ***checkMySequence*** [71], which automatically detects register-shift errors in protein models built into cryo-EM maps, depends on the quality of input map and model, specifically on map preprocessing and local map resolution. Further, it uses relatively long test fragments at lower local map resolutions which may result in missing short, local register-shift errors. Nevertheless, checkMySequence yields useful results where manual residue-by-residue validation is difficult. The accuracy of ***doubleHelix*** [72] for nucleic-acid sequence identification, assignment, and validation in cryo-EM maps depends on the accuracy of input nucleic-acid models. Further, performance of its neural-network classifier is influenced by map quality and since nucleic acid regions in cryo-EM maps are typically poorly resolved, it may affect accurate sequence assignment. Moreover, the signal in density maps is often limited to distinguishing only two nucleobase types - purines and pyrimidines - complicating sequence assignment. Nevertheless, doubleHelix can successfully assign sequences to models built in cryo-EM maps at local resolution as low as 5.0 Å. The performance of ***EMsequenceFinder*** [75] to accurately assign an amino acid sequence to backbone fragments is dependent on the accuracy of the backbone traced in the input cryo-EM density map. Reliable backbone traces can be generated by using methods listed in Table 2. Further, the program at present only considers fragments with α-helical and β-strand backbones and other secondary structure elements, such as coils and loops will be considered in future. Like other methods, its prediction accuracy declines as the map resolution deteriorates. Map preprocessing such as denoising and density modification could improve prediction accuracy by reducing noise.

**CNN-classifier** [76], one of the early deep learning methods developed to detect protein secondary structures in medium resolution cryo-EM density maps, is trained on a limited number of simulated cryo-EM maps (Table 4) which may limit its performance on experimental cryo-EM density maps for secondary structure detection. **Emap2sec** [77] can only analyze protein maps and, while it detects density regions of secondary structure, it does not actually place the α-helices and β-strands within these detected density regions. Its successor, **Emap2sec+** [78] can detect both protein secondary structure elements and nucleic acids in cryo-EM density maps and also improves the accuracy of protein secondary structure detection. **Haruspex** [79], a method for identifying protein secondary structures and nucleic acids in cryo-EM density maps, may incorrectly label semi-helical structures, β-hairpin turns, and polyproline type II (PII) helices as α-helices and loosely parallel structures that lack the typical hydrogen-bond pattern as β-strands. In future versions of Haruspex, it will predict additional classes like β-turns, polyproline helices, and membrane detergent regions to reduce the number of misidentified secondary structure elements and improve the overall accuracy of the method. The performance of **EMNUSS** [80] is sensitive to map resolution, especially at middle-resolution (5.0 to 9.5 Å) as EMNUSS was trained on small number of maps (120 maps) at that resolution range. Further, incorrect predictions may occur on unusual density volumes as the network of EMNUSS is not trained on lower-resolution maps. Therefore, more maps are needed for training to improve the performance and robustness of EMNUSS. **DeepSSETracer** [81], a tool to identify protein secondary structures in medium resolution maps, is designed to operate on component maps with a maximum size of 100 voxels in any dimension and requires prior segmentation of the cryo-EM density map containing multiple chains. As such its detection performance depends on the quality of the segmented maps. **HaPi** [83] struggles to determine the handedness of structures lacking clear or sufficient α-helices. This difficulty increases with lower resolutions, particularly as the resolution approaches 5.0 Å, where less information regarding handedness is encoded. The minimum resolution for determining the hand of an α-helix is its pitch, which on average is 5.4 Å. Above this resolution, an α-helix appears as a cylinder, which possesses no apparent handedness. **CryoSSESeg** [84], a tool to detect protein secondary structures in medium-resolution cryo-EM maps, segments the entire density of a protein chain so that the model can learn from complete secondary structures and avoiding artifacts of cutting secondary structures as seen in patch-based segmentation. However, it is important to note that this approach may require huge amount of memory when working with large protein chains. **EMInfo** [86], an automated tool for predicting protein secondary structure elements and nucleic acids in cryo-EM density maps, may struggle to accurately predict structural categories in several cases such as at the terminal ends of nucleic acids with weak density signals, in lower-resolution density regions, at the interfaces between different macromolecule categories and may identify coils with strands.

The main factor limiting the detection accuracy of **AAnchor** [87] at high resolution maps (2.2 Å) is the limited experimental cryo-EM data used (3 maps, 1.8 - 2.3 Å range) for the training of the AAnchor algorithm. Given that the number of high resolution experimental cryo-EM maps have increased since then (Figure 1), the current version of AAnchor may provide improved results. The large-scale dataset of **A^2^-Net** [88], a tool for amino acid determination in a cryo-EM density map, is derived from simulated cryo-EM densities and could potentially benefit from including experimental maps in its training. Trained on a larger set of simulated cryo-EM maps, **Cascaded-CNN** [90], a tool for predicting protein backbone structures in high-resolution maps, could potentially improve its performance by including experimental cryo-EM maps in its training data. Further, threshold value selected to normalize input maps before processing for model building, a common step in many methods, may affect final structure prediction. The authors of Cascaded-CNN have developed a method that automatically estimates the correct threshold value for density maps to alleviate this issue [191]. **Structure Generator** [93], a template-free method to build protein structures in cryo-EM maps, is also trained on simulated cryo-EM density profiles of proteins (Table 4), and its performance may vary on experimental cryo-EM maps. Due to the noise in experimental cryo-EM density maps, models created by **DeepTracer** [94] may sometimes show poor fit to the density map including misplaced side-chains, geometric and connectivity errors thus requiring downstream model rebuilding and refinement using tools such as molecular dynamics flexible fitting (MDFF) [148] and *phenix_real_space_refine* [66]. Furthermore, DeepTracer does not build structures for other macromolecules such as nucleic acids in cryo-EM density maps. **DeepTracer-2.0** [96] extends DeepTracer’s capability to model protein-DNA/RNA macromolecular complexes from the cryo-EM density maps. The performance of DeepTracer-2.0 to accurately model protein-DNA/RNA complexes depends upon accurate segmentation to extract the density maps for separate macromolecules. As such its performance decreases on low resolution cryo-EM density maps, often observed for nucleic acids, due to the challenges in accurately segmenting such maps. As increasingly high-resolution density maps become available for protein-DNA/RNA complexes, its performance will improve. DeepTracer-2.0 also depends on Brickworx model [73] for postprocessing and building models of DNA/RNA from the predicted voxels which could be time consuming. Future work will make DeepTracer-2.0 less reliant on third-party software for post-processing. **DeepMM** [97] can introduce errors or uncertainties into the built models in cryo-EM density maps that have low overall resolution or low-resolution regions. Because DeepMM is designed for single chains, it relies on Segger [192] to segment the original map into separate regions to model multi-chain complexes. As such its performance on multi-chain complexes is dependent on the quality of the map segmentation. **SEGEM** [104] showed lower amino acid prediction accuracy for the experimental test set than the simulated set due to the varying local resolution and noise in experimental maps, which makes it difficult to normalize them for convolutional neural network (CNN) training. **SEGEM++** [104], which combine SEGEM and AlphaFold2 also struggles to build complete models in cryo-EM density maps with varying local resolution and noise impacting its ability to identify native Cα sites. Nevertheless, performance of SEGEM and SEGEM++ is expected to improve with advancements in 3D image semantic segmentation and cryo-EM image processing. **SegmA** [105], a tool designed for residue type segmentation of a cryo-EM density map, may mislabel amino acid voxels as nucleotides at protein-nucleic acid interfaces and regions with low resolutions. Further, it may also struggle to accurately label amino acids with similar properties as they appear similar in cryo-EM maps. Like other methods, its performance is sensitive to map resolution and degrades as the resolution lowers for amino acids.

Nevertheless, SegmA is a powerful tool to distinguish between protein residues and nucleotides. The resolution of input cryo-EM maps affects the model building performance of **ModelAngelo** [106]. The quality of the initial graph generated by the convolutional neural network (CNN) and the accuracy of amino acid classification, both crucial for mapping sequences to the main chain, are significantly affected from low-resolution cryo-EM data. Poor amino acid classifications can lead to errors in sequence assignments and subsequent incorrect chain assignments, especially in complexes with many similar sequences. Protein structure prediction methods, such as AlphaFold2, can be integrated with the features derived from low-resolution cryo-EM maps to automate model building as demonstrated by some of the methods listed in Table 3. For nucleic acids, assigning the correct sequence of nucleobases (the equivalent of side chains in RNA or DNA) to the predicted backbone becomes challenging especially at resolutions around 3.5 Å as it becomes difficult to differentiate between individual purines (adenine from guanine) or pyrimidines (cytosine from thymine/uracil). *doubleHelix*, a neural network classifier to identify nucleobases, can be combined with ModelAngelo predicted nucleic acid backbone to automate the model building of complete nucleic acid structures. While **CryoREAD** [111]accurately identifies nucleobase positions, here also base-type detection remains less precise, especially at lower resolutions, affecting accuracy of sequence matching. Nevertheless, CryoREAD provides accurate nucleic acid structures even from lower-resolution maps for manual sequence assignment. For large structures, where the final model from CryoREAD might have backbone gaps or incorrect base-pairing, manual refinement with tools like COOT is required. CryoREAD aims to improve nucleic acid structure modeling by integrating secondary structure information for nucleic acids predicted with high accuracy. While **SMARTFold** [114] creates more complete and accurate atomic structures compared with other state-of-the-art methods, its memory intensive architecture limits its use to model protein sequences with a maximum of 2,500 residues. In addition, the multiple sequence alignment (MSA) search is time-consuming. Future versions of SMARTFold will address these issues with a smaller model that uses smaller feature channels to reduce memory use and a protein language model to replace the time-consuming MSA search. **EMRNA** [116] performance can be inconsistent in low-resolution maps (4.0 – 6.0 Å) as it is challenging to build RNA models at these resolutions. For large RNA molecules, EMRNA accurately places RNA backbone fragments but struggles to correctly order these fragments sequentially, often leading to models with a high root-mean-square deviation (RMSD). Since many of these models still retain the correct overall fold, minor manual adjustments can often fix them. However, third party software is often required to build base atoms to achieve high accuracy. Unlike **EM2NA** [119], which can automatically segment and identify DNA/RNA regions in raw cryo-EM maps, EMRNA requires users to properly mask the map around the RNA being modeled for optimal performance. It is challenging for EM2NA to automatically assign the correct nucleotide type for the built model, often requiring prior knowledge and expert intervention to complete the modeling. EM2NA may struggle to recognize non-standard helical geometry of nucleic acids including structures like bulges or flipped-out nucleotides that can sometimes occur in protein-nucleic acid complexes. Nevertheless, EM2NA accelerates model building significantly compared to manually starting from scratch. While **Cryo2Struct** [122] can correctly identify most Cα atoms and build highly accurate atomic models, creating comprehensive and accurate models of large protein structures from density maps alone remains a significant challenge. This is primarily because resolution heterogeneity in cryo-EM maps limits the resolvability of every residue of a protein. Even prediction errors for a few Cα atoms, in noisy cryo-EM maps, can limit the accurate prediction of long, continuous stretches of polypeptide chains. Supplementing cryo-EM density maps with additional inputs, such as AlphaFold2 protein structures and symmetry of the multi-chain protein complexes can produce more accurate and complete predictions. **Cryo2Struct2** [165] improves Cryo2Struct by integrating cryo-EM data with advance structure prediction capabilities of AlphaFold3. **EmodelX** [124], a method for building protein complex structures in cryo-EM maps, could be extended to include modeling of DNA/RNA-protein assemblies and small molecules. **EModelX(+AF)** [124], which combines EModelX and AlphaFold, effectively handles both low-resolution cryo-EM density maps and inaccurate AlphaFold predictions while modeling protein complexes in such density maps. Though **CryFold** [125] is less reliant on map resolution for model building, modeling at low resolutions remains challenging due to the difficulty in accurately identifying protein side chains in these regions. **DeepCryoRNA** [127], which only models RNA in the cryo-EM density maps, may have variable performance for maps with resolutions below 4.5 Å. Future iterations of DeepCryoRNA are anticipated to extend to modeling DNA or DNA/RNA-protein complexes in cryo-EM maps like CryoREAD, EM2NA and DeepTracer. One of the main limitations of **E3-CryoFold** [131] is that it currently only models the residue backbone without considering side chains. Side chain modeling of E3-CryoFold derived protein backbone can potentially be achieved using tools like SCWRL4 [92], similar to the approach taken by DeepTracer. Further, E3-CryoFold can result in inconsistent root-mean-square deviation (RMSD) between predicted and target structures due to lack of constraints during generation of atom coordinates. Incorporating atom coordinate information, derived directly from density maps, with the E3-CryoFold predictions can help with this issue.

**Table 3.**
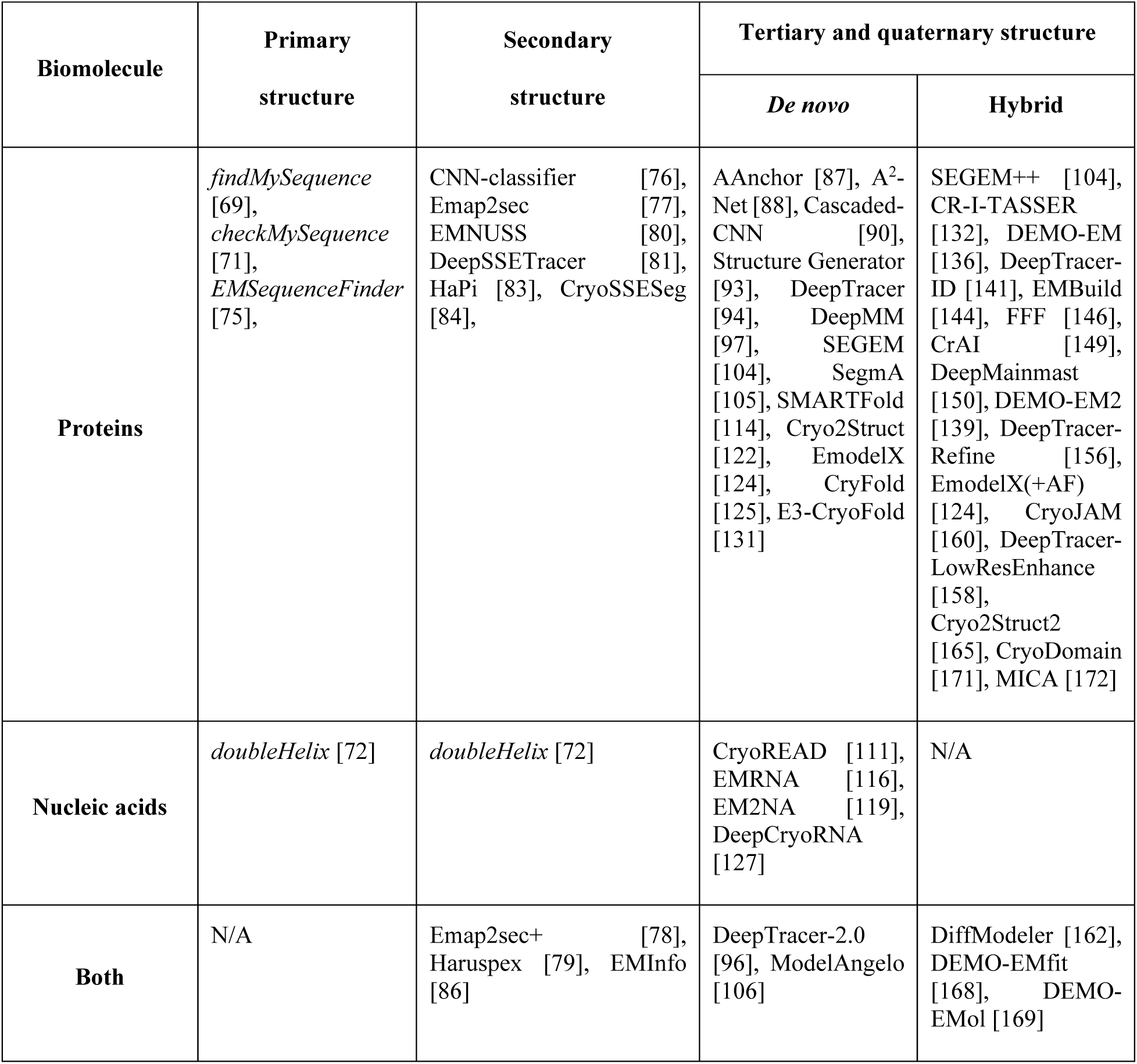
Available cryo-EM structure modeling tools across resolutions and molecules.

**Table 4.**
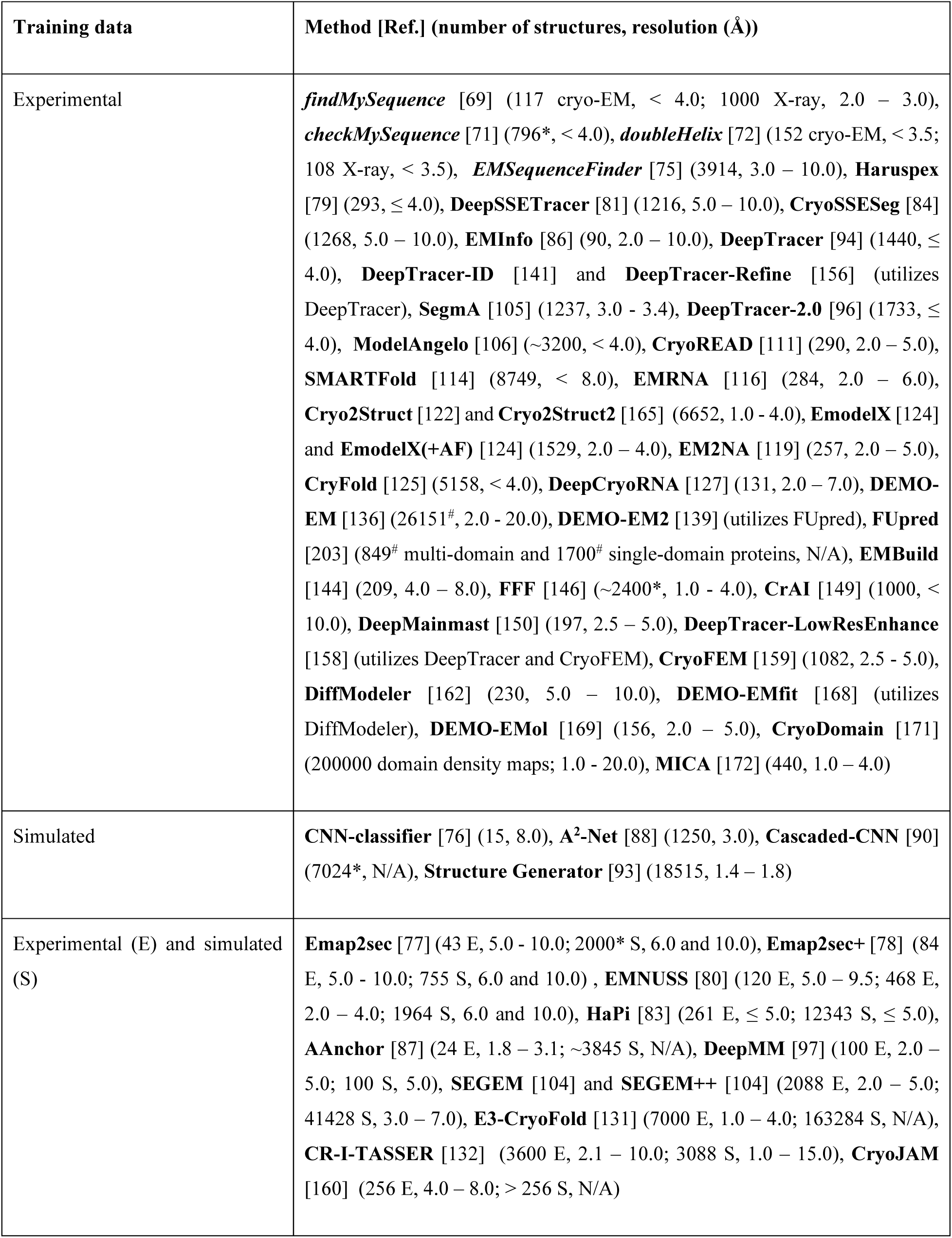
Characteristics of training datasets used by deep learning-based automated model building methods in cryo-EM. (*size of train and test sets from the dataset size is not known; ^#^number of cryo-EM structures is not known)

**Table 5:**
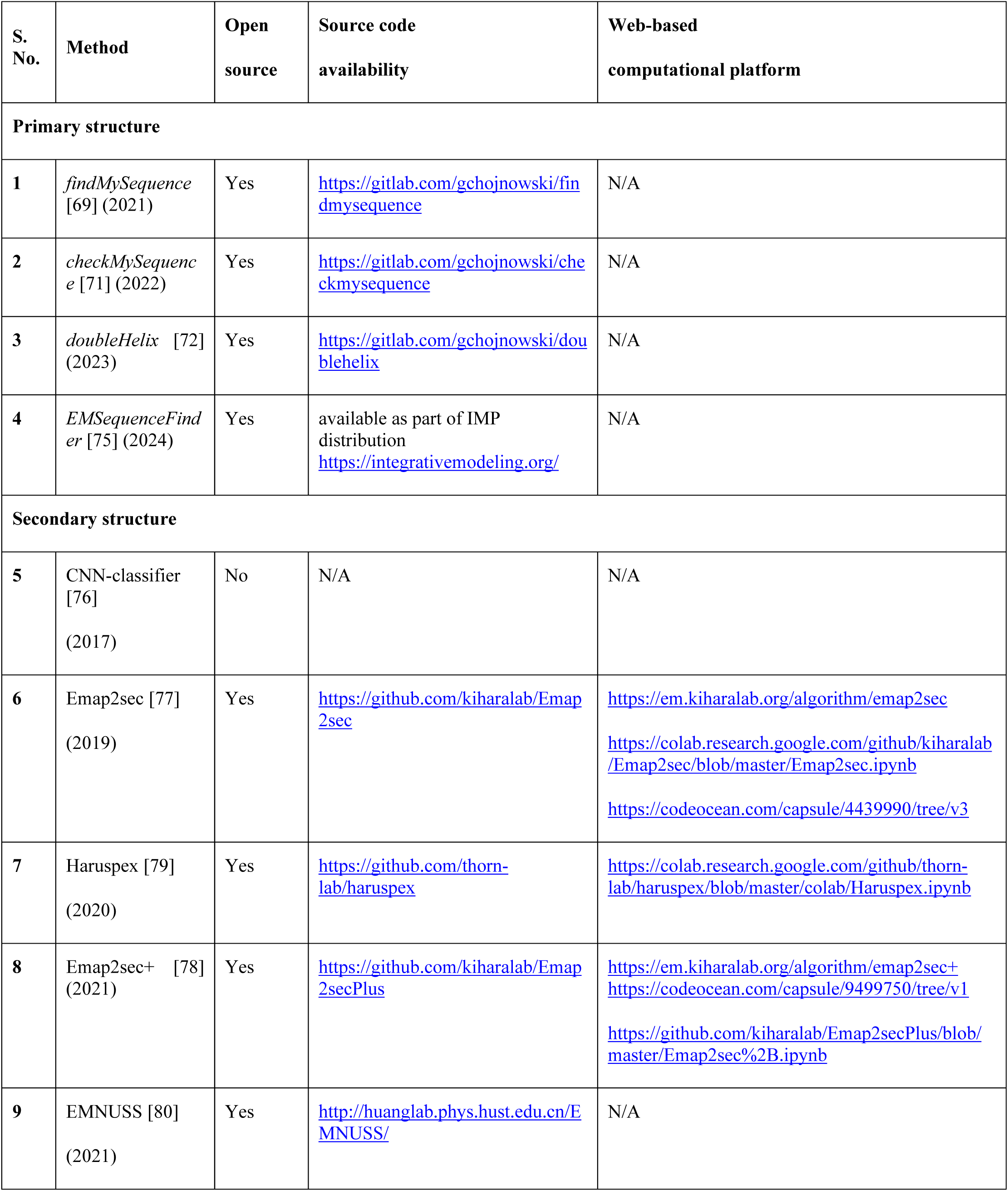
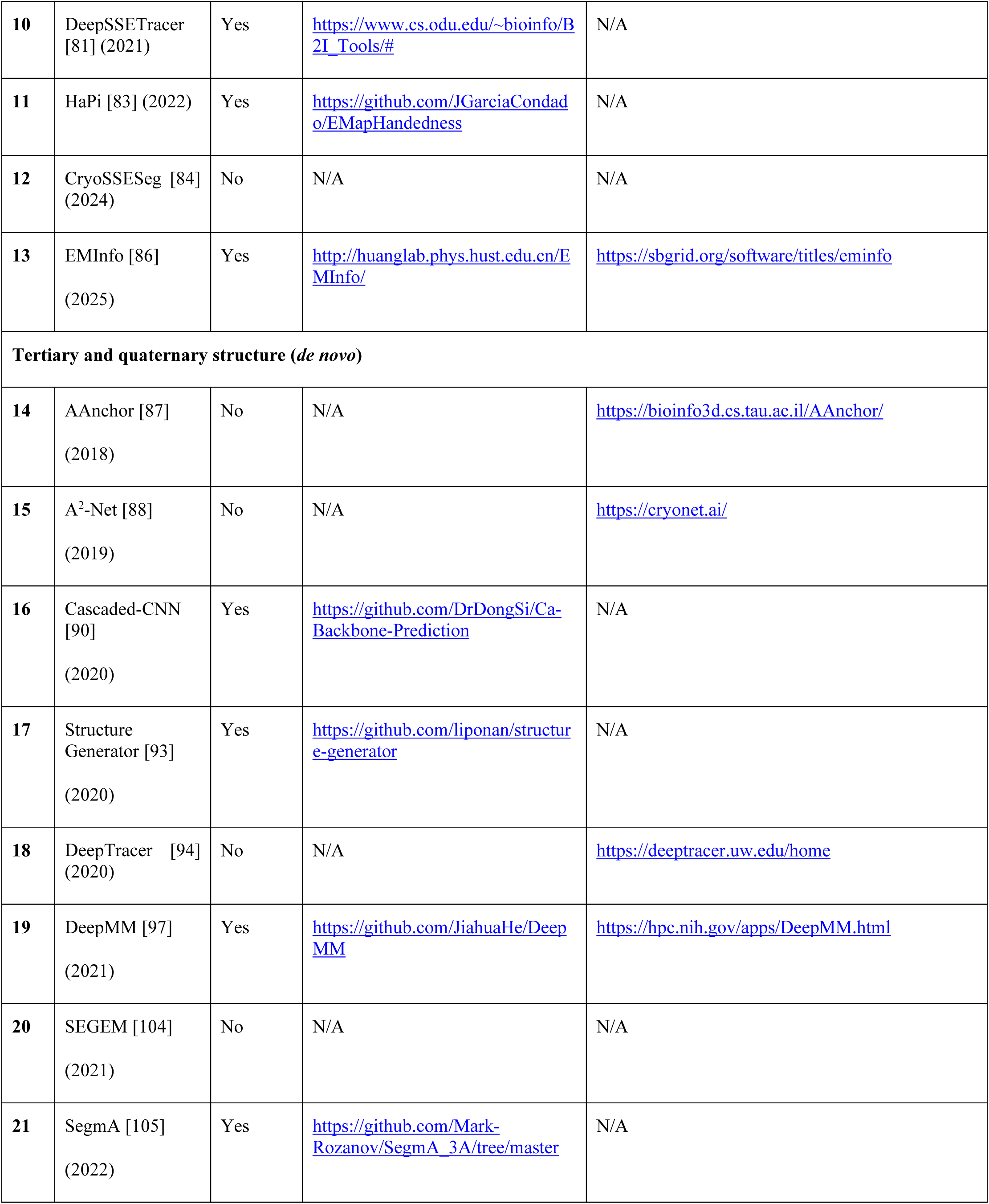
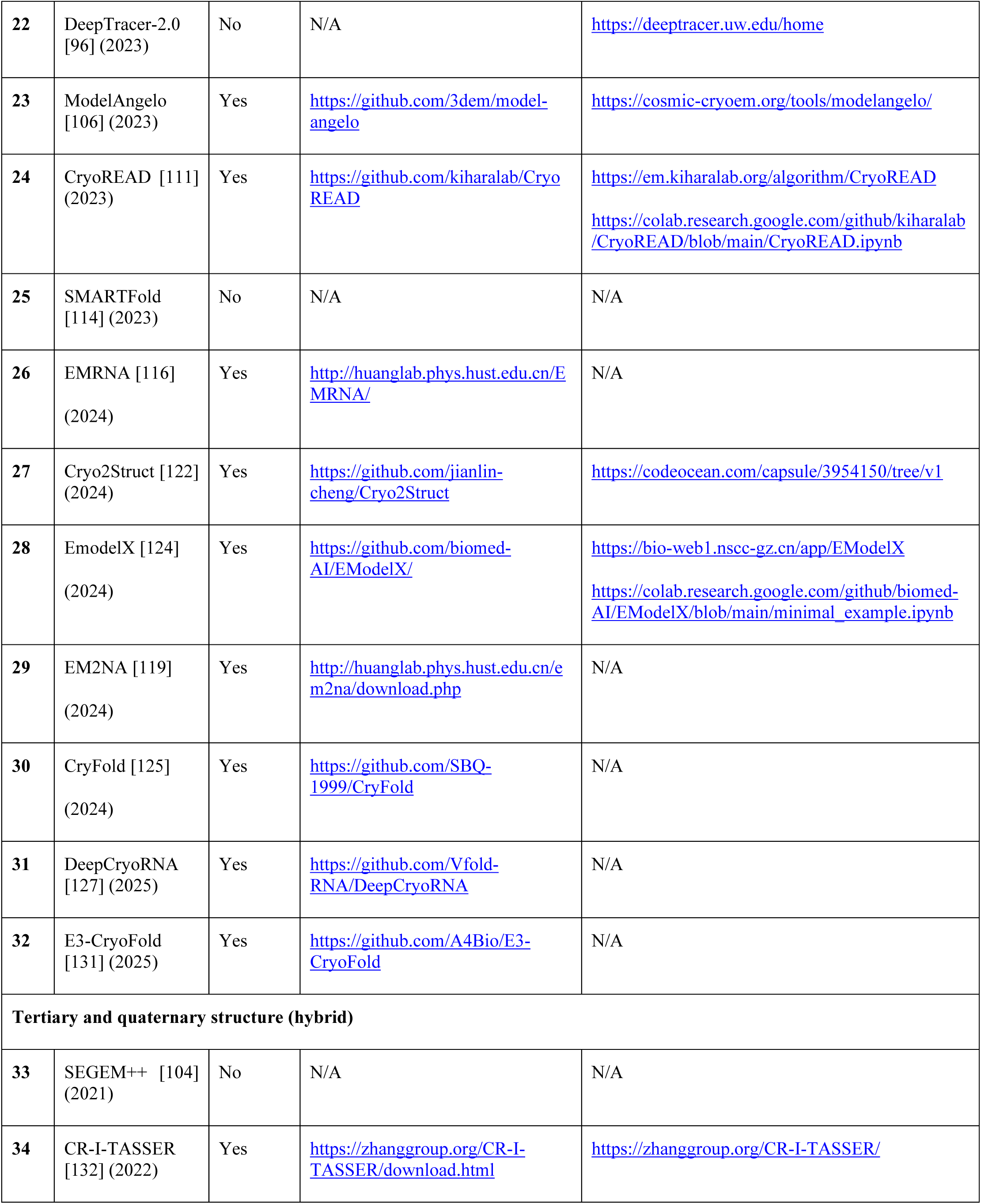
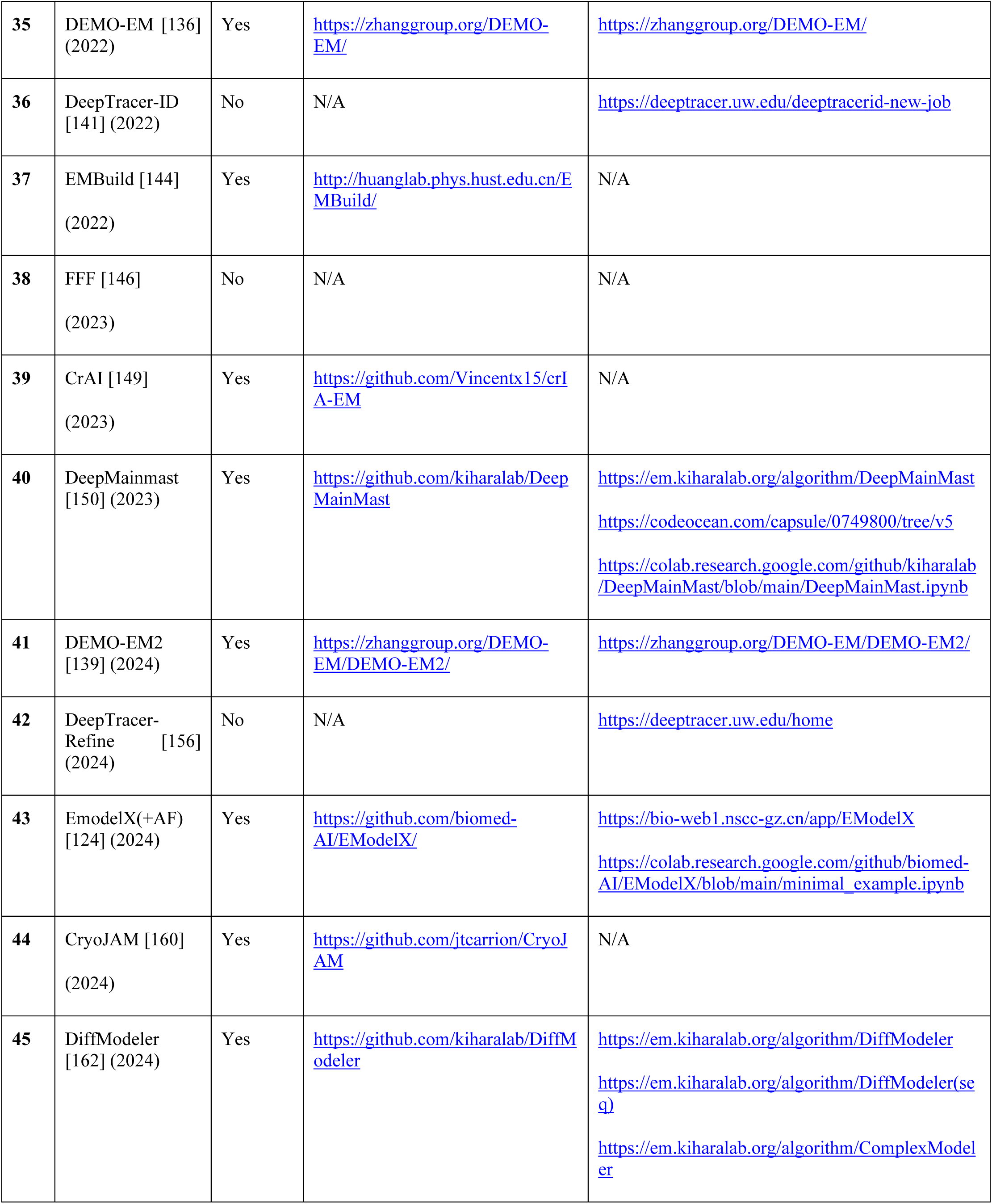
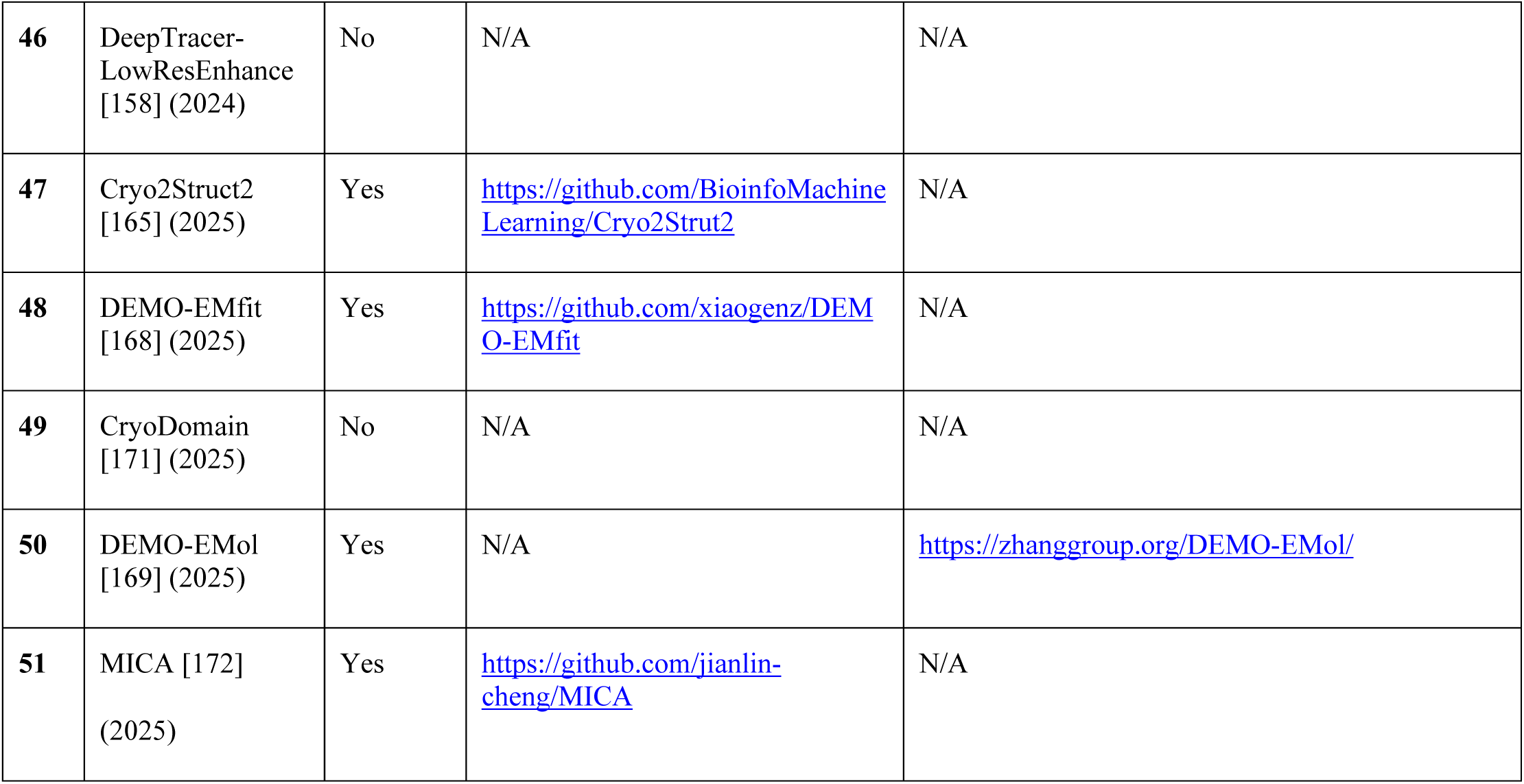
Availability of deep learning-based automated model building methods in cryo-EM.

The accuracy of 3D convolutional neural network (3D-CNN) and therefore **CR-I-TASSER** [132] to predict Cα trace decreases as resolution deteriorates. CR-I-TASSER also struggles to model target structures containing loops and disordered regions. Future iterations of CR-I-TASSER will combine 3D-CNN with multiple sequence alignment (MSA) to improve accuracy of its Cα trace and model prediction. Although CR-I-TASSER is currently limited to model monomer proteins that require manual segmentation of cryo-EM maps, the CR-I-TASSER pipeline will be extended to model larger protein–protein/protein–nucleic acid complexes. **DEMO-EM** [136] pipeline, which assembles multi-domain proteins, requires manual segmentation of maps and could be improved by including automatic segmentation techniques to reduce computational time. Another limitation of DEMO-EM is that the initial domain models, which are created by D-I-TASSER independent of the cryo-EM density data, can be inaccurate and can negatively affect the quality of the final models. Future versions of DEMO-EM will improve the accuracy of the final models by using cryo-EM density data restraints to guide the initial domain model generation. **DEMO-EM2** [139], which constructs protein complex models, improves many aspects of DEMO-EM such as preprocessing density maps to suppress noise effects on fitting protein models and using advanced strategies to prevent the algorithm from getting stuck in a local optima. However, DEMO-EM2 can be further improved by using deep learning techniques, to identify distinct domain regions and predict distances and orientations between protein chains and domains directly from density maps, and use them as constraints to guide the assembly of protein complexes. **DEMO-EMol** [169] server, which assembles protein-nucleic acid complex structures from cryo-EM maps, aims to create an end-to-end model-to-map fitting method by using deep learning, and improve fitting efficiency by matching local and global geometric features. Additionally, it will add explicit energy terms that retain base pairing in output nucleic acid models thus improving their accuracy. **DEMO-EMfit** [168], which fits atomic structures into cryo-EM density maps, outlines several points for potential improvement in its current version. To improve both the accuracy and efficiency of fitting, end-to-end deep learning approaches can be used to directly learn structural poses from point clouds extracted from cryo-EM density maps. For better evaluation of fitting quality, a combination of physics-based Gaussian mixture models and DOT scores from VESPER may be more reliable than traditional correlation coefficient metrics. **DeepTracer-ID** [141], which can identify component proteins directly in cryo-EM maps, may struggle with modeling small proteins (<100 amino acids) at a resolution of 4.2 Å or better. Small proteins often exist within larger complexes, yet it is rare for them to achieve near-atomic resolution in cryo-EM studies. This lack of high-quality data means there is a scarcity of detailed experimental structural information for these proteins requiring input from protein prediction methods such as AlphaFold. **DeepTracer-Refine** [156], which uses DeepTracer predicted structures to refine AlphaFold predictions, is limited to docking domains from AlphaFold structures in the cryo-EM density maps and at present cannot refine the structure at residue level. Further, DeepTracer-Refine struggles with AlphaFold predictions where domains are incorrectly folded as it is designed to only correct inaccurate domain arrangements in AlphaFold predictions. Future development of DeepTracer-Refine will explore residue-level refinement of the backbone. **DeepTracer-LowResEnhance** [158], which extends DeepTracer’s ability to build models in low-resolution cryo-EM maps, is currently limited to proteins and future investigation will explore its application to model DNA/RNA structures from low-resolution cryo-EM maps. **EMBuild** [144] accurately builds protein complex models into cryo-EM maps when provided with the accurate predictions of individual chains at the fragment or domain level. Its performance suffers when regions in the input structures, such as those from AlphaFold2, are inaccurate requiring removal of inaccurate or disordered parts of the predicted structures based on their pLDDT values. Like other methods, **FFF** [146] is sensitive to map resolution. Its performance degrades as resolution decreases and can lead to errors in the protein sequence alignment during the fragment recognition stage ultimately affecting subsequent structure-fitting steps. Although **CrAI** [149] is a state-of-the-art method for detecting antibodies in cryo-EM maps, it performs better on Fabs than on VHHs. This is because VHHs are smaller, have non-canonical binding modes, and less training data is available for them. Despite these challenges, CrAI still outperforms other methods at detecting VHHs. Despite successful prediction, CrAI may have less precision as it relies on a template instead of using a ground truth structure. The performance of **DeepMainmast** [150] decreases as the local map resolution becomes lower requiring AlphaFold2 models in low resolution regions of map to correct inaccurate backbone tracing in the density. DeepMainmast also tends to model residues in helices more accurately than β-strands and loops. **CryoJAM** [160], an automated tool for fitting protein homologs into cryo-EM maps, may struggle to identify small differences between very similar homolog structures. It also faces significant challenges with low-resolution maps, or those containing membrane or detergent density. Diffusion model of **DiffModeler** [162] struggles to generate accurate backbone traces for low resolution regions limiting subsequent structure fitting to regions with higher local resolution. Further, DiffModeler’s pipeline is currently limited to model protein complexes but will extend its capabilities to model protein/DNA/RNA complexes in future. Also, accuracy of DiffModeler is influenced by the quality of the initial AlphaFold2 models it uses, fitting individual protein domains rather than entire proteins could help overcome the inaccuracies of the initial models. DiffModeler can potentially be used for cryo-electron tomography (cryo-ET) with maps at a resolution of 15 Å or better. The performance of **CryoDomain** [171], a tool to identify protein domains in low resolution cryo-EM maps, may be influenced by the quality and accuracy of the initial segmentation of the map into its individual components. Although CryoDomain may sometimes retrieve incorrect domain types, resulting in false positives, it is more accurate and robust across a wide range of resolutions compared to other methods. **MICA** [172] struggles with some large protein complexes when cryo-EM data is noisy or has missing density, and when AlphaFold3 predictions do not align well with the experimental maps. Future efforts will focus on enhancing sequence and chain registration, considering symmetry of protein complex during sequence registration, and integrating advanced side-chain prediction algorithms directly into the deep learning framework of MICA.

## 8 Conclusions

Deep learning has undeniably accelerated and automated every aspect of cryo-EM structure determination pipeline. This review presented a comprehensive and up to date survey of the current landscape of deep learning-based methods (∼50) for automated model building into cryo-EM density maps. We outlined common conceptual strategies across diverse methods and summarized key aspects of these tools, including their training datasets, neural network architectures, prediction tasks, the types of biomolecules they build, and their availability as servers or publicly accessible code. By discussing the capabilities and limitations of available methods, we hope this synthesis will serve as a valuable resource and stimulate future improvements in the field.

The resolution revolution in cryo-EM now increasingly enables high-resolution structure determination of biomolecular complexes with interpretable density for bound small molecules and drugs. However, most of the current deep learning-based model building methods focus primarily on building the biomacromolecule itself. This highlights a pressing need for developing robust deep learning tools for automatically identifying and building the bound small molecules [193], thereby significantly accelerating structure-based drug-discovery efforts. Addressing this requires overcoming challenges related to training data. While deep learning-based methods train on large datasets, their application is often limited by the availability of map-model pairs that perfectly capture experimental reality. Real cryo-EM maps exhibit conformational heterogeneity, which the associated model does not capture. This discrepancy hinders the development of deep learning-based methods capable of modeling true conformational ensembles [194–196] or accurately representing ligand binding states [197]. Therefore, efforts are required to curate experimental datasets of conformationally-resolved experimental map-model pairs for training deep-learning-based methods [198].

Looking ahead, as neural network architectures continue to evolve rapidly, multi-modal approaches that integrate different architectures, diverse data types, and geometric deep learning [42,50] to learn from multi-dimensional data show significant promise for capturing diverse structural features of biomolecules. Deep learning-based automated model building for cryo-electron tomography (cryo-ET) datasets represent a crucial next step, holding the potential to identify and build macromolecules directly within their native cellular environment and ushering in a new era of *in-situ* structural biology [199].

## 10 Box 1: Glossary section

**SE(3):** Special Euclidean Group of degree 3 is a group of all rigid motions in 3D space and combines rotations and translations. It describes transformations that preserve shape and size of objects.

**Monte Carlo Tree Search (MCTS):** MCTS is a powerful algorithmic framework used for decision making in sequential decision processes. MCTS builds a search tree by simulating many possible moves and selecting the most promising path.

**Profile Hidden Markov Model (profile HMM):** A profile hidden Markov model (profile HMM) is a probabilistic model constructed from a multiple sequence alignment (MSA) of related protein or nucleic acid sequences. It captures both conservation and variability in the alignment by defining match states for conserved positions, insert and delete states for regions where extra residues or gaps are likely, and probabilities that describe which residues are observed at each aligned column. The model is parameterized by two types of probabilities: transition probabilities, which specify the likelihood of moving between match, insert, and delete states as the sequence progresses along the profile, and emission probabilities, which define the likelihood of observing a specific residue from a match or insert state. To assess whether a new sequence belongs to the family, the model computes the probability that the sequence could be generated by traversing the profile’s states, emitting the observed residues along the way. The higher this probability, the more likely the new sequence is a member of that family.

**Dynamic Programming:** Algorithms designed for solving optimization problems by breaking them into simpler, smaller and overlapping subproblems.

**Constraint Programming:** Techniques developed for solving combinatorial problems where the solution must satisfy a set of constraints.

**Mean-Shift Algorithm:** Non-parametric clustering technique to locate the highest density points of a distribution or simply identify dense regions in a dataset.

**Traveling Salesman Problem (TSP) Solver:** The goal of the solver is to find the shortest possible route that visits a set of points exactly once and returns to the starting point as a way of solving the well-known combinatorial optimization TSP problem.

**Vehicle Routing Problem (VRP) Problem:** The goal of the solver is to optimize routes for multiple vehicles visiting a set of locations or points while satisfying a set of constraints. VRP is a generalization of TSP.

## Notes

### Competing Interest Statement

The authors have declared no competing interest.

### Funding

- National Institute of General Medical Sciences: R35GM133598
- New York University: Research and Professional Development grant, Startup grant

