## Supporting Information for "Connecting the dots: deep learning-based automated model building methods in cryo-EM"

Since their introduction, deep learning–based methods have automated and accelerated the atomic model–building step in cryogenic electron microscopy (cryo-EM), enabling molecular-level interpretation of density maps. These advances have yielded mechanistic and functional insights into protein–protein and protein–nucleic acid assemblies, thereby informing drug discovery and therapeutic development. When integrated with mass spectrometry, cryo-EM, and cryogenic electron tomography (cryo-ET), deep learning approaches are advancing efforts toward *in situ* structural characterization, capturing the organization and interactions of macromolecules within their native cellular contexts. More broadly, deep learning–based tools have revolutionized model building in structural biology, transforming computational methods into indispensable resources for the community. These approaches now tackle diverse challenges, from detecting contaminants within heterogeneous cryo-EM datasets to resolving three-dimensional structures of macromolecules at near-atomic resolution. As these technologies mature, they are catalysing methodological innovation across the discipline and enabling analyses that were previously unattainable. Examples of cryo-EM studies where these tools have made notable contributions are described below.

While homo-oligomerization is vital for many cellular processes, RNA quaternary structures have rarely been reported. Cryo-EM structures of four RNA families, including ARRPOF, OLE, ROOL and GOLLD at 2.6 to 4.6 Å resolutions revealed their diverse homo-oligomeric assemblies and offered crucial insights into the molecular basis of intermolecular interactions driving RNA multivalency and its potential functional relevance [1]. The atomic models of ARRPOF, OLE and ROOL with cryo-EM maps at better than 4 Å resolution were built *de novo* with the help of RNA-specific model building tools such as EMRNA, EM2NA and CryoREAD which are designed to accurately recognize the RNA backbone at these resolutions.

Cryo-EM structures of arrestins in complex with phosphorylated atypical chemokine receptor 3 (ACKR3) have revealed how arrestins differentially recognize unique phosphorylation barcodes on G-protein-coupled receptors (GPCRs), which are believed to regulate distinct cellular responses [2]. This study utilized DiffModeler to calculate a diffusion model, a class of generative models in deep learning, to enhance structural features including backbone positions in the map, facilitating downstream structure fitting and modelling tasks.

While previous cryo-EM structures showed overall architecture of coagulation factor V (fV), mechanism that keeps fV in its inactive state was unresolved because of an intrinsic disorder in its B domain. A 3.2 Å cryo-EM structure of fV short, a splice variant with a truncated B domain that exhibits constitutive activity, further advances understanding of the mechanism that keeps fV in its inactive state [3]. Models generated using the CR-I-TASSER web server were used to partially guide the manual placement of residues in the B domain during model building in the cryo-EM maps.

Cryo-EM structure of human antiviral protein APOBEC3G (A3G) bound to HIV-1 Vif reveals RNA as a molecular glue that enables Vif to suppress A3G through both ubiquitin-dependent and independent pathways [4]. This study used Haruspex, a convolutional neural network trained to detect DNA/RNA versus protein in density maps, to annotate unaccounted-for density sandwiched between A3G and Vif in the cryo-EM maps as oligonucleotides.

The anti-cancer drug bortezomib exhibits potent activity against *Mycobacterium tuberculosis (Mtb)* by binding to its caseinolytic protease (Clp) protease system. A cryo-EM structural study, which utilized CryoNet (improved version of $A^2$-Net) tool to predict initial models for model building and refinement into the cryo-EM maps, showed that bortezomib disrupts the Mtb Clp system's intricate regulation by binding to its protease sites [5].

Pre-mRNA splicing is essential for gene expression, removing non-coding introns and joining coding exons. While the major spliceosome handles most introns (U2-type), the minor spliceosome processes a rare subset (U12-type). Together with mass spectrometry data, cryo-EM was used to structurally characterize minor spliceosome and its unique protein components [6]. Protein sequences and the density map were used as inputs to CryoNet to generate atomic models of protein components PPIL2, SCNM1, CRIPT, and RBM48.

In photosynthesis, light energy is harvested by antenna pigments and delivered to reaction centers (RCs) to drive the electron transfer (ET) processes. During cryo-EM structure determination, CryoNet was used to *de novo* build atomic models of the *Chloracidobacterium thermophilum* reaction centers (CabRC) enabling identification of novel subunits and providing crucial insights into the ET process within the CabRC [7].

Upon infection, podophages typically inject their viral genome into host cells through a transmembrane ejectosome. Cryo-EM structure of *Anabaena* cyanopodophage A4 revealed a pentameric pre-ejectosome at the core of its capsid. Importantly, the initial models for the protein components of A4 were generated by ModelAngelo and CryoNet for fitting into the cryo-EM density maps, enabling a better understanding of podophage assembly and DNA ejection and highlighting potential applications of A4 in synthetic biology [8].

The 2-oxoglutarate dehydrogenase complex (OGDHc) plays a crucial role in regulating the TCA cycle. A combination of cryo-EM, cryo-ET and subtomogram averaging (STA) revealed the intact structure of native OGDHc and each of its subunit from *Sus scrofa* heart tissue highlighting the unique factors driving the architecture of the intact, native OGDHc [9]. Here, the initial atomic modelling was performed using CryoNet.

Photosynthetic organisms use various light-harvesting systems to capture light and transfer energy to reaction centers (RCs). *Chloroflexus aurantiacus*, with its integral membrane antenna (LH complex) and extrinsic chlorosomes, is an excellent model for studying these processes. A 3.05 Å cryo-EM structure of a minimal RC–LH photocomplex from *Cfx. aurantiacus* (CaRC-LH) provides understanding of the possible assembly patterns of RC-LH and the existence of chlorosomes in *Cfx. Aurantiacus* [10]. Notably, the atomic model of the CaRC–LH complex was initially auto-built using CryoNet thus accelerating the structure determination process.

Cryo-EM structure of human Microfibril-associated glycoprotein 4 (MFAP4), a key protein in various diseases like organ fibrosis, chronic obstructive pulmonary disease, and cardiovascular disorders reveals multiple protein-binding surfaces on MFAP4, providing a clearer understanding of its crucial biological functions [11]. CryoNet's trained neural network was used to accurately estimate the local resolution of the MFAP4 density maps thus assisting structure determination process.

Coronaviruses enter host cells when their spike proteins recognize specific receptors. TMPRSS2 and sialoglycan have been recently identified as receptors for HCoV-HKU1. A cryo-EM study, utilizing CryoNet to *de novo* construct the HCoV-HKU1C spike structure based solely on its amino acid sequence and the cryo-EM density map, provides structural basis of complex viral-host interactions during coronavirus entry, laying the groundwork for developing new therapeutics against coronavirus-associated diseases [12].

Understanding how the SARS-CoV-2 spike protein moves bidirectionally within host cells (between the endoplasmic reticulum, Golgi, and plasma membrane) is crucial for comprehending virus assembly and creating better genetic vaccines. By using sequence analysis and AlphaFold modeling, researchers devised a strategy for the internal tagging of the spike protein. Cryo-EM structure determination and model building of the spike ecto-domain aided by DeepMainMast, revealed conformations compatible with ACE2 receptor binding demonstrating the feasibility of the internal tagging strategy [13].

Circularly permuted (CP) introns generate circular RNA (circRNA) through a process called back-splicing. Cryo-EM studies, leveraging auto-DRRAFTER and CryoREAD for *de novo* model building into the cryo-EM maps, reveal the structures of a natural CP intron in various back-splicing states at 2.5–2.9 Å [14]. These structures show how circularly permuted (CP) introns undergo back-splicing, which could lead to advancements in circRNA research and the development of new therapeutics.

BREX (BacteRiophage EXclusion) anti-phage systems differentiate host from invading DNA through site-specific epigenetic DNA methylation. In Type I BREX, the BrxX (PglX) methyltransferase plays a crucial role. A 2.2-Å cryo-EM structure of *E. coli* BrxX bound to target dsDNA, where the initial model of the DNA was built using cryoREAD server facilitating model building and analysis, reveals molecular details of the BREX DNA recognition [15].

Type II topoisomerase DNA gyrase uses ATP hydrolysis to negatively supercoil DNA, a process thought to involve a chiral DNA loop. High-resolution cryo-EM structures of *E. coli* gyrase holoenzyme in the chirally wrapped state bound to a 217 bp linear DNA fragment provides molecular basis of how gyrase constrains a positively supercoiled loop [16]. A cryoREAD-generated DNA model was used for guidance and to verify DNA positioning in the model. These structures reveal an updated mechanism of enzyme catalysis.

Short prokaryotic argonaute (pAgo) proteins, particularly in conjunction with TIR or Sir2 (SPARTA or SPARSA) proteins, act as antiviral systems to combat phage infections. The cryo-EM structures of

SPARSA with and without nucleic acids, at resolutions of 3.1 Å and 3.6 Å, respectively, provides mechanistic insight into the antiphage role of the SPARSA system [17]. DeepTracer was used to trace the backbone structures in the cryo-EM maps, guiding the fitting of the AlphaFold2-predicted structures into the density map.

Sperm motility, driven by the axoneme - a microtubule-based structure, is essential for successful reproduction. A recent study reveals high-resolution cryo-EM structures of axonemal doublet microtubules (DMTs) from sea urchin and bovine sperm, representing both external and internal fertilizers. Notably, deep learning-based tools were used to pinpoint many of the approximately 60 proteins that decorate sperm DMTs [18]. One method involved automatically building models into cryo-EM maps using either ModelAngelo or DeepTracer, then querying the predicted protein sequences against sequence databases with protein-BLAST. Another approach used a structure-based strategy where backbone traces from density maps were fed into DeepTracer-ID to search PDB or AlphaFold2 libraries for identification. These results provide structural foundations for understanding sperm evolution, motility, and dysfunction at a molecular level.

Type IV pili (T4P) are common, multi-functional surface appendages in archaea, essential for adhesion, motility, and communication. A recent cryo-EM study on *Saccharolobus islandicus* revealed the atomic structures of two distinct adhesive T4P, assembled from the same pilin polypeptide under different growth conditions, demonstrating that a single secretion system can produce vastly different filament structures [19]. For protein identification, DeepTracer-ID was used, which uses a combination of AlphaFold predictions and cryo-EM density maps. Further, deep learning-based model building methods - ModelAngelo or DeepTracer – were used to determine the amino acid sequence directly from the cryo-EM map followed by a BLAST search against the *S. islandicus* REY15A proteome.

Using cryo-EM, cryo-ET, and proteomics, detailed architecture of axonemal doublet microtubules (DMTs) from mammalian sperm has been resolved revealing a comprehensive model of the mammalian sperm DMT, pinpointing 181 proteins and identifying integrated chemical and mechanical regulatory elements within the axoneme [20]. For identifying proteins at intermediate resolution (around 5 Å), manually traced helices were used to query AlphaFold2 databases using DeepTracer-ID.

Conjugation, a key driver of antibiotic resistance spread, involves horizontal gene transfer between cells via a mating pilus. A recent study used cryo-EM to reveal the atomic structures of three conjugative pili, two from archaea and one from bacteria demonstrating that archaeal pili are homologous to their bacterial counterparts [21]. Notably, DeepTracer-ID was used to determine the pilin identity of one of the archaea directly from cryo-EM density maps, which was then named as TedC.

While bacterial biofilms are extensively studied due to their role in disease, archaeal biofilms have received less attention. Cryo-EM structure of a novel class of archaeal surface filaments from *Pyrobaculum calidifontis*, named archaeal bundling pili (ABP) suggests shared evolutionary and mechanistic links between bacterial and archaeal biofilms [22]. Notably, deep learning-based tools such as AlphaFold, DeepTracer and DeepTracer-ID were used for protein identification and model building into the cryo-EM maps,

The contractile tail of *Agrobacterium tumefaciens* bacteriophage Milano is uniquely flexible and bent, unlike typical rigid counterparts used by other bacteriophages and bacterial secretion systems. Cryo-EM structures of the Milano tail, including its unique sheath-tube complex, baseplate, and receptor-binding proteins reveals the electrostatic properties of its tail tube and sheath for its flexible-to-rigid transformation during contraction [23]. While AlphaFold was used to predict all 127 Milano proteins, DeepTracer-ID along with manual fitting was used to place these predicted structures into the cryo-EM density maps thus accelerating structure determination.

White spot syndrome virus (WSSV), one of the largest DNA viruses and a major crustacean pathogen, employs a capsid crucial for genome handling, transitioning between rod and oval shapes. Cryo-EM structural investigation along with DeepTracer-assisted protein identification and model building in the cryo-EM density maps, revealed a unique ring-stacked assembly mechanism of the rod-shaped capsid offering structural insights into the pressure-driven genome release [24].

Cryo-EM structures of archaeal type IV pili (AT4P), where initial models were built into the cryo-EM density maps using DeepTracer-ID, revealed how the structural diversity of AT4Ps likely allowed for an AT4P to evolve into a supercoiling archaeal flagellar filament [25].

Deep learning-based model building methods can assist in identifying contaminant proteins from heterogeneous cryo-EM datasets without prior knowledge of its protein sequence. Cryo-EM data processing combined with DeepTracer server for *de novo* model construction was used to identify a contaminant, *Escherichia* phage YDC107 [26]. Here, the sequence extracted from DeepTracer generated model was searched against the NCBI database using BALSTp to successfully identify the phage YDC107 tail protein.

RNA-dependent RNA Polymerase 2 (RDR2) is vital for RNA-mediated gene silencing, and its activity is tightly coupled with the activity of DNA-dependent Pol IV. Recent full length cryo-EM structure of RDR2 at 3.1 A, explains how Pol IV arrest facilitates RDR2's role in gene silencing [27]. Notably, Emap2sec was used to assess secondary structure propensity in the cryo-EM density maps thus assisting in the model building.

Cryo-EM and cryo-ET in combination with mass spectrometry and machine learning are facilitating visual proteomics. Leveraging this approach, distinct oligomeric protein complexes from *Azotobacter vinelandii* extracts were identified. This involved using deep-learning-based tools like CryoID [28], DeepTracer, and ModelAngelo to identify previously unknown proteins from the cryo-EM density maps, including phosphoglucoisomerase (Pgi1) in a novel decameric state, glutamine synthetase (GlnA), and bacterioferritin (Bfr) [29].

Misfolded protein aggregates, forming filamentous cellular inclusions, are a hallmark of neurodegenerative diseases. A recent study reveals that amyloid fibrils of a 135-amino acid fragment of TMEM106B are unexpectedly common across diverse human neurodegenerative disorders, including those characterized by TDP-43, tau, or α-synuclein pathology [30]. Deep learning tools such as DeepTracer, cryo-ID [28], and *findMySequence* [31] were instrumental in identifying the fibril's constituent protein and building the atomic model in the cryo-EM density maps of TMEM106B fibrils derived from postmortem human brain tissues.

Intermediate species in amyloid filament assembly are thought to be key in neurodegenerative diseases, making them important therapeutic targets. Using time-resolved cryo-EM, a study investigated the *in vitro* assembly of truncated tau (residues 297–391) into filaments linked to Alzheimer's disease and chronic traumatic encephalopathy [32]. Here, atomic models were built automatically using ModelAngelo, offering crucial structural insights into amyloid nucleation and opening the way to new therapeutic strategies.

The Integrator complex terminates RNA polymerase II (Pol II) at gene promoter-proximal regions. Cryo-EM structures, aided by ModelAngelo for peptide identification and its *de novo* model building into the 2.7 A cryo-EM density map, reveal three functional states of the complete Integrator–PP2A complex and a three-step Pol II termination model involving significant rearrangements [33].

A new hallucination-based approach for efficient, high-quality protein backbone design across various scales and applications without retraining was validated through the experimental production and characterization of over 100 proteins, and their structures confirmed through high-resolution structure determination [34]. Specifically, for two large ~100 kDa proteins, cryo-EM yielded high-resolution maps (2.7 Å and 3.3 Å), enabling out-of-the box *de novo* construction of accurate atomic models using ModelAngelo.

The Cav3.2 T-type calcium channel is a target for pain and epilepsy medications. Cryo-EM structures of Cav3.2, alone and with four different antagonists, at 2.8-3.2 Å resolution identified key residues for drug selectivity [35]. Notably, Cav3.2Apo initial model was automatically generated with ModelAngelo from the cryo-EM maps.

BamA, the core of the β-barrel assembly machine (BAM), is crucial for inserting and folding proteins into Gram-negative bacterial outer membranes. Peptide macrocycles, Peptide Targeting BamA-1 (PTB1) locks BamA into a closed state by binding an extracellular site, while PTB2 traps it in an open state by binding a luminal site offering a new template for antibiotic discovery targeting BamA and other dynamic integral membrane proteins [36]. The initial models for membrane-embedded macrocycle, PTB2-lg-1 and PTB2-lg-2, were first predicted by ModelAngelo for subsequent model adjustments and refinement in the cryo-EM density maps.

DNA G-quadruplexes (G4s), non-B-form DNA structures, stalls replisomes by arresting the CMG helicase. Cryo-EM structures of stalled CMG complexes, where *de novo* model building into the cryo-EM maps was performed using ModelAngelo, revealed a folded G4 lodged inside the central CMG channel, arresting translocation [37].

Unlike typical CRISPR-Cas systems that degrade foreign DNA, Type IV-A CRISPR-Cas systems suppress gene expression through a nuclease-independent pathway. Cryo-EM structures of two distinct Type IV-A complexes (IV-A1 and IV-A3) bound to target DNA, where the type IV-A1 initial model was generated *de novo* using ModelAngelo, provides a structural foundation for developing new genome-editing tools [38].

Humans detect odors using two G protein-coupled receptor families: odorant receptors (ORs) and trace amine-associated receptors (TAARs), with TAARs resembling aminergic receptors. Cryo-EM structure of murine mTAAR7f, bound to N,N-dimethylcyclohexylamine (DMCHA) and a G protein, at 2.9 Å resolution, where the *de novo* model of TAAR7f was generated from the cryo-EM density map and protein sequence using ModelAngelo, suggests TAAR activation mirrors β-adrenoceptors [39].

The type-1 ryanodine receptor (RyR1) is crucial for skeletal muscle excitation-contraction coupling, acting as an intracellular calcium release channel. Recent cryo-EM study demonstrates a key role of an additional transmembrane helix, S0, in the channel gating. ModelAngelo and PSI-BLAST were used to identify the sequence of S0 as part of RyR1 itself [40].

ApOR5-Orco heterocomplex is an odorant receptor (OR), ligand-gated ion channels, that detects and distinguish a wide variety of chemicals. During cryo-EM structure determination, DeepTracer (version 1.0), an automated deep learning method, first generated an initial backbone model. facilitating integration of AlphaFold2 predictions of the ApOR5 and ApOrco subunits in the complete reconstruction of the complex [41].

The N-degron pathway marks proteins for degradation, with yeast's Ubr1 enzyme playing a key role in tagging them with ubiquitin chains using partner enzyme Ubc2. Deep learning model building tool, DeepTracer, provided a complete model with Ubc2 correctly positioned and 80% of Ubr1 correctly built into the cryo-EM density map [42].

DeepTracer generated models were instrumental in determining high-resolution (2.7 to 3.9 Å) cryo-EM structures of the human small subunit processome, which mediates early maturation of the small ribosomal subunit [43].

Caveolin, a monotopic membrane protein composed of 11 protomers, is crucial for shaping cell membrane invaginations called caveolae. DeepTracer was used to generate an initial prediction of amino acid coordinates from the regions of the cryo-EM map with well-resolved secondary structures for one of the protomers to build the model of the caveolin complex [44].

Initial monomer models of chimallin (ChmA), a bacteriophage nuclear shell protein, were built in the density maps using DeepTracer web server, shedding light on its architecture and mechanisms for its nucleation, growth, and scaffolding role for phage factors involved in transport, cytoskeleton interactions, and viral maturation [45].

The SARS-CoV-2 protein Nsp2 has been implicated in a wide range of viral processes. DeepTracer web server generated a quality initial model from the high-resolution region of the cryo-EM density map and the full Nsp2 sequence which served as a template for generating full atomic model [46].

Inorganic polyphosphate (polyP), an ancient energy metabolite, is synthesized from ATP and translocated into the vacuolar lumen by the ubiquitous vacuolar transporter chaperone (VTC) complex.

In the cryo-EM map of VTC complex at 3 Å resolution, the overall structure of the complex was first *de novo* main-chain modeled by DeepTracer, resulting a backbone atomic structure followed by docking of AlphaFold predicted domains into this backbone atomic structure to aid in model building [47].

Retrons are bacterial genetic elements that provide anti-phage defense. From the cryo-EM density of the *E. coli* Ec86 retron complex, the density map of RNA/DNA or protein was identified using deep learning method Emap2sec+ v1.0 and the extracted protein densities were *de novo* main-chain modeled using DeepTracer v1.0 providing a structural basis for the optimization of retron-based genome editing systems [48].

Cryo-EM was used to analyze a raw human kidney microsomal lysate, simultaneously identifying and solving the structures of four distinct kidney enzymes, implicated in diabetes [49]. Notably, DeepTracer directly identified protein primary sequences from the high-resolution cryo-EM density maps, confirmed by NCBI protein blast and proteomics.

Noncoding RNAs, like human Alu RNA, can repress gene transcription by RNA polymerase II (Pol II). Cryo-EM structures of Pol II bound to Alu RNA reveal that Alu RNA mimics DNA and RNA binding during transcription elongation [50]. Here, model building for Pol II and the minimal Alu-RA RNA was achieved by combining DeepTracer for *de novo* building and manual secondary structure-based model building.

Integrating systems and structural biology is a promising new field, utilizing the potential of cryo-EM for atomic-level tissue proteomics. Single-particle cryo-EM was used to analyze three heterogeneous fractions from human liver mitochondrial lysate, simultaneously resolving high-resolution structures of nine essential mitochondrial enzymes involved in critical metabolic functions [51]. Notably, DeepTracer successfully revealed protein sequences from the high-resolution maps, confirmed using BLASTP against *Homo sapiens*.

Cryo-EM structure of *Francisella* protein FTN_1118 (*Francisella* Periplasmic Metalloprotein, FPM13), a novel 13 kDa periplasmic protein unique to the *Francisella* genus suggests its role in metal transport or detoxification. During the cryo-EM structure determination, a 3D model trace was generated using DeepTracer and analyzed against the *F. novicida* protein database using cryoID [28], thus identifying FTN_1118 (FPM13) [52].

3-methylcrotonyl-CoA carboxylase (MCC) is a vital mitochondrial enzyme for leucine catabolism. *Leishmania tarentolae* MCC (LtMCC) forms long, inactive filaments, observed in their structures at 3.4, 3.9, and 7.3 Å using cryo-EM thus providing new insights on the role of polymerization in carboxylase function. Partial model predictions from DeepTracer (using the 3.9 Å map) and subsequent cryoID [28] analysis (combined with mass spectrometry data) confirmed LtMCC α-subunits as a filament component [53].

Motor neurons form synapses with skeletal muscle, releasing acetylcholine (ACh) to activate nicotinic ACh receptors (AChRs) and initiate contraction. *De novo* models for high-resolution cryo-EM density maps of AChRs were generated using ModelAngelo providing structural insights into pathogenic mechanisms behind congenital myasthenic syndromes [54].

Mitochondria generate ATP via oxidative phosphorylation through five respiratory chain complexes. ModelAngelo, in combination with cryo-ET and cryo-EM, enabled visualization of the native structures and organization of major mitochondrial complexes of *Chlamydomonas reinhardtii*. Specifically, focus-refined cryo-EM maps (all at resolutions better than 3 Å) for complexes I, III, and IV were used as inputs to ModelAngelo for *de novo* sequencing and the resulting peptide chains were identified against the *Chlamydomonas reinhardtii* UNIPROT database using BLAST assisting retrieval of AlphaFold2 models [55].

The plastid-encoded RNA polymerase (PEP) transcribes most chloroplast photosynthesis genes, aided by nuclear-encoded PEP-associated proteins (PAPs). Cryo-EM structures of native PEP reveal PAPs encasing the core polymerase, promoting assembly and stability [56]. Here, a model of the entire PEP complex was built using ModelAngelo which enabled fitting of the AlphaFold predicted PAP subunits into the density maps via structural alignment to the PEP complex.

ModelAngelo can also successfully generate *de novo* RNA models from cryo-EM maps as exemplified by its application on generating 3D structures of large bacterial RNAs, GOLLD, ROOL, and OLE, from high resolution cryo-EM maps, thus providing insights into the biological importance of ornate, protein-free RNA quaternary assemblies [57].

Measles virus (MeV) is a highly contagious pathogen whose polymerase machinery (L and P proteins) is vital for replication and transcription, making it an antiviral drug target. Atomic models for MeV polymerase machinery were automatically built using ModelAngelo into the cryo-EM density maps of two distinct MeV polymerase complexes: Lcore-P and Lfull-P-C, offering architectural and molecular insights into MeV polymerase mechanisms [58].

*Pseudomonas aeruginosa* is a critical pathogen requiring new antimicrobials, a challenge increasingly addressed by phage therapy using bacterial viruses. High-resolution structural atlas of Pa193, a therapeutic *Pseudomonas* phage with a contractile tail, was determined using bioinformatics, proteomics, and cryo-EM analysis [59]. Specifically, out of the 21 distinct polypeptide chains comprising its icosahedral capsid, neck, contractile tail, and baseplate, ModelAngelo successfully identified and built the atomic models for scaffolding protein gp24 and tape measure protein gp41.

Mycobacteriophage Bxb1, a well-characterized double-stranded DNA virus, shows promise for treating *Mycobacterium* infections. Here, ModelAngelo was used to make primary assignments of protein subunits into the cryo-EM density and to establish stoichiometry, enabling determination of complete structure and atomic model of phage Bxb1 [60].

The complete degradation of heparan sulfate (HS), a glycosaminoglycan, relies on the essential acetylation of its terminal α-D-glucosamin catalyzed by Heparan-α-glucosaminide N-acetyltransferase (HGSNAT). ModelAngelo was used to build the initial HGSNAT model into the cryo-EM density map, providing structural insights into this critical acetylation reaction [61].

F1Fo ATP synthase generates ATP using a proton motive force. Its F1-ATPase catalytic core hydrolyzes ATP through conformational changes, involving central γ subunit rotation and catalytic β subunit opening/closing. The cryo-EM structure of an axle-less *Bacillus sp*. PS3 F1-ATPase, where an initial axle-less F1-ATPase model was built using ModelAngelo, suggest that the complete γ subunit is crucial for coordinating efficient ATP binding in F1-ATPase [62].

Pannexin-3 (PANX3) is an ATP-permeable channel involved in diverse biological processes. Notably, the initial PANX3 atomic models were built using ModelAngelo into cryo-EM maps of human PANX3 at 2.9–3.2 Å resolution, providing foundational insights into pannexin channel properties [63].

Cell division cycle 25 phosphatases (CDC25A, B, C) activate CDKs by dephosphorylating their glycine-rich loop, regulating cell cycle transitions. The 2.7 Å cryo-EM structure of CDK2-cyclin A in complex with CDC25A, where ModelAngelo was used for automated model building of the trimeric complex, details its architecture and critical protein-protein interactions thus highlighting the role of CDC25 in CDK regulation, potentially aiding anti-cancer drug development [64].

The unusual mitoribosome of the apicomplexan parasite *Toxoplasma gondii* assembles from over 50 extremely short rRNA molecules. In the cryo-EM structure of the mitoribosome, model building involved initial protein fragment assignment to the density map using ModelAngelo and querying the resulting *de novo* sequences against annotated proteins in ToxoDB. Finally, initial models from the AlphaFold database for likely hits were fitted against these *de novo* chains reveals functioning of this mitoribosome [65].
